# Structural dynamics and immunogenicity of the recombinant and outer membrane vesicle-embedded Meningococcal antigen NadA

**DOI:** 10.1101/2024.01.30.577382

**Authors:** Valeria Calvaresi, Lucia Dello Iacono, Sara Borghi, Enrico Luzzi, Alessia Biolchi, Barbara Benucci, Ilaria Ferlenghi, Ilaria Peschiera, Fabiola Giusti, Lucia E. Fontana, Zhong-Yuan Kan, Zaira Spinello, Marcello Merola, Isabel Delany, Kasper D. Rand, Nathalie Norais

## Abstract

The structure and conformation adopted by protein vaccine antigens significantly influence the exposure of their antigenic determinants. Structural knowledge of antigens in native state could drive the design of recombinant vaccines that resemble their cognate native forms, although such information is often difficult to obtain, particularly for membrane proteins. Here, we assessed the structural and functional features of the native Neisseria Adhesin A (NadA), a meningococcal trimeric outer membrane protein included as soluble recombinant antigen in the 4CMenB vaccine. We used hydrogen-deuterium exchange mass spectrometry (HDX-MS) to generate a structural model of NadA and to compare the fold and structural dynamics of the recombinant NadA as soluble vaccine form, and the native NadA *in situ*, as embedded in meningococcal outer membrane vesicles (OMVs), complementing the HDX data with electron microscopy imaging. While their overall structures are similar, conformational differences between the two forms were observed. Especially, OMV- embedded NadA appears more susceptible to trimer opening than its cognate soluble antigen, suggesting that NadA in its native membrane could display a larger antigenic surface. Accordingly, we show that mice immunized with OMV-embedded NadA elicited antibodies with superior bactericidal activity and capable of better preventing bacterial adhesion compared to the soluble antigen. Collectively, these data support the hypothesis that protein vaccine antigens presented in native-like environments can elicit a more potent immune response than recombinant forms.

## Introduction

Subunit vaccines are often based on recombinant proteins, which can be rationally designed with excellent efficacy and safety profiles and produced with high purity^1^. Membrane proteins often represent suitable surface antigens of bacterial pathogens. However, membrane proteins are challenging to be recombinantly produced in their ‘native’ conformation as extracting them from their membrane environment can compromise their native folding. The genome-based *reverse vaccinology* approach is a valuable technology to identify surface exposed bacterial membrane proteins suitable as vaccine candidates^2–4^ and their subsequent structural characterization (*structural vaccinology*) can enable a rational vaccine design^5,6^. In 2013, a reverse vaccinology-derived multicomponent subunit vaccine against *Neisseria meningitidis* serogroup B (MenB) was approved (4CMenB)^7,8^. One of the four protective components of 4CMenB is Neisseria adhesin A (NadA)^3,8^, a protein antigen able to induce high levels of bactericidal antibodies both in animal models^3,9,10^ and in humans^11^. NadA variant 3, the vaccine variant and hereinafter referred as NadA, is a flexible and elongated outer membrane protein that mediates adhesion to and invasion of host cells during bacterial infection and colonization of the human respiratory tract^12^. NadA is a member of trimeric autotransporter adhesins and shares with them the typical modular organization^13^. It has an N-terminal head region (aa 24-73) projected away from the bacterial surface that is involved in host recognition and binding, an elongated central stalk (aa 73-344), and a beta-barrel C-terminal anchor that embeds into the bacterial outer membrane (aa 345-405). NadA still lacks a complete structural determination, although the X-ray structure of a short yet trimeric construct (NadA24-170) has been solved^14^. The stalk of NadA24-170 adopts a coiled-coil organization and is made of canonical heptad repeats and two uncanonical undecad repeats. Traditionally, in coiled coils, heptad repeats, denoted *‘abcdefg’*, are characterized by hydrophobic residues at the first *‘a’* and fourth *‘d’* positions, which are packed inside the coiled coil^15^. In the uncanonical undecads *‘abcdefghijk’*, hydrophobic residues are present also at the eighth position *‘h’*^15^. NadA head domain is a small globule resulting from the partial opening of the trimer, and is shaped in a tripartite wing-like structure flanking an N-terminal heptad coiled-coil region^14^. The head and stalk domains of NadA, referred together as passenger domain, spontaneously assemble into a stable trimer^16^, allowing it to be included as recombinant soluble antigen in the current 4CMenB vaccine (NadA24-350), without the C-terminal beta-barrel anchor. The recombinant soluble NadA form has already proven to elicit an effective immune response, with anti-NadA antibodies promoting bacterial killing and preventing the initial bacterial colonization^11^. However, it has been shown that not all anti-NadA antibodies isolated from vaccinated individuals are able to recognize the native antigen exposed on the bacterial membrane^11^, which may suggest that the protein conformation and epitope exposure of the soluble vaccine NadA and the bacterial NadA could differ or that not all epitopes are accessible in the native membrane bound form. In this work, we thus aimed at comparing the conformational dynamics of the soluble antigen NadA, included in the current vaccine, and NadA *in situ*, in its native meningococcal membrane, as well as evaluating the immune response elicited by both NadA forms, in order to identify its optimal immunogenic state. To study NadA *in situ*, we leveraged the capability of outer membrane vesicles (OMVs) to faithfully reproduce the bacterial membrane environment, hence of carrying antigens in an identical state as that adopted onto the bacterial surface. OMVs are spherical buds of outer membrane naturally released by Gram-negative bacteria, therefore their lipid, outer membrane protein and lipopolysaccharide (LPS) composition are the native ones^17,18^. For interrogating the conformational dynamics of NadA, we applied hydrogen-deuterium exchange coupled to mass spectrometry (HDX-MS), a technology that has proved suitable for the analysis of membrane proteins in complex lipid environments^19–24^, including OMVs^25^. In HDX-MS, a protein is exposed to a deuterated buffer and the exchange of its back-bone amide hydrogens with deuterium atoms is measured as a function of time using mass spectrometry. The HDX rate of these labile hydrogens is a highly sensitive reporter of the local structure and conformational dynamics^26^, such that HDX data can be of use to fill in missing structural data from electron microscopy (EM) or X-ray crystallography, or combined to them, to predict protein structure^27^.

By using HDX-MS data, we generated a structural model of the full-length NadA, gaining new information on its entire structure, and unveiled that NadA can adopt an open trimer conformation. While the overall structures of the soluble vaccine antigen and the OMV-embedded form appear similar, we reveal differences at their conformational level, and that the open trimer conformation is favored in OMVs. Finally, we show that OMV-embedded NadA elicits in mice antibodies of higher functional quality, capable of better preventing meningococcal adhesion to epithelial cells and with superior bactericidal activity compared to the recombinant antigen. As a whole, our data offer a structural basis for a potential optimization of the MenB antigen NadA and, in a broader perspective, showcase that antigen functional immunogenicity can be improved by displaying antigens onto OMVs.

## Results

### Structural model of NadA

The recombinant vaccine antigen NadA (NadA_24-350_), hereinafter referred as rNadA, fails to crystalize, likely due to its flexibility^14,28^. We thus lack a complete determination of its structure, which prompted us to use HDX-MS to infer local dynamic information linkable to its secondary structural elements^29,30^. Our HDX-MS analysis yielded a map of 181 peptides, with 100% sequence coverage and high spatial resolution (10.24 of redundancy) (**Fig. S1**, table 1, uptake plots in Supplementary dataset 1). Deuterium labelling was performed for several time intervals and under different temperature conditions to expand the dynamic and time window studied, thus sampling the HDX with high temporal resolution^31^. The results showed that rNadA contains regions that undergo rapid, medium, and slow HDX kinetics both in the head and stalk domains (**Figs. 1a and S2**), and interestingly, that specific regions manifested bimodal HDX behavior, hence a low- and high-mass envelope distribution (**Fig. 1b**) (see paragraph: *Identification of an open trimer conformation in NadA*). By using HDsite, we deconvolved peptide-level HDX data to residue-resolved kHX values of individual or groups of few back-bone amides^32^. The *k*HX provides information on the degree of engagement of back-bone amide hydrogens in intramolecular hydrogen-bonding, hence on the presence of stable secondary/tertiary structure^33^, which we combined with the structural information available for NadA24-170 (PDB: 6EUN) to model the part of the stalk that still lacks structural determination. *k*HX values were extracted at single residue level for 19% of the sequence and with an average resolution of 3.92 amino acids for the rest (**Fig. S3**, table 2 and *k*HX plots in Supplementary dataset 2). In NadA_24-170_, log*_k_*_HX_ values less than -1 (protection factor > 250 and ranging from 10^3^ to 10^6^) were extracted for residues within the coiled coil head and stalk, whereas log*_k_*_HX_ values greater or equal to -1 (protection factor ≤ 250) were calculated for residues located in non-coiled coil regions, namely the wings (aa 65-74) and the N-terminal part of the head domain (residues 24-28), which in part lacks electron density in the X-ray structure previously determined^14^. For the part of the stalk of unknown structure (residues 168-350), most backbone amides displayed log*_k_*_HX_ < -1, indicative of coiled coil elements, except for the regions spanning residues 252-254, 264-267 and 288-293, which suggests that the coiled coil stalk of NadA is interrupted by flexible structural elements. Based on the HDX data and the analysis of NadA primary sequence and coiled coil periodicities, from L168 the most likely assignment of NadA stalk resulted in fourteen heptad repeats and two uncanonical undecad repeats, one spanning residues 210-220 and the other one spanning residues 270-280; with two flexible regions of non-coiled coil structure assigned to residues 252-265 and 284-297 (**Fig. 1c**, Table 2). The log*_k_*_HX_ values of the amides comprised in the C-terminus (aa 329-350) indicate a highly flexible and unstructured region (log*_k_*_HX_ near to 1 and protection factors < 10), corroborated by the presence of numerous disruptive and polar amino acids, typical of intrinsically disordered segments^34^. We could thus infer that this part, which we denominated NadA tail, lacks a stable coiled coil structure (**Fig. 1d**). A low-resolution Cryo-EM structure confirmed that rNadA predominantly adopts an elongated and flexible structure (30 nm in length), with a clearly resolved rod-like organization. A globular density with a diameter of 5 nm, fitting the N-terminal head domain, was visible at one end, whereas a hook-like bend with a dimeter of 3.7 nm was observable at the opposite end. The latter corresponds to the NadA tail, for which also Cryo-EM data supports the lack of a coiled coil organization (**Figs. 1d and S4**).

**Fig. 1.**
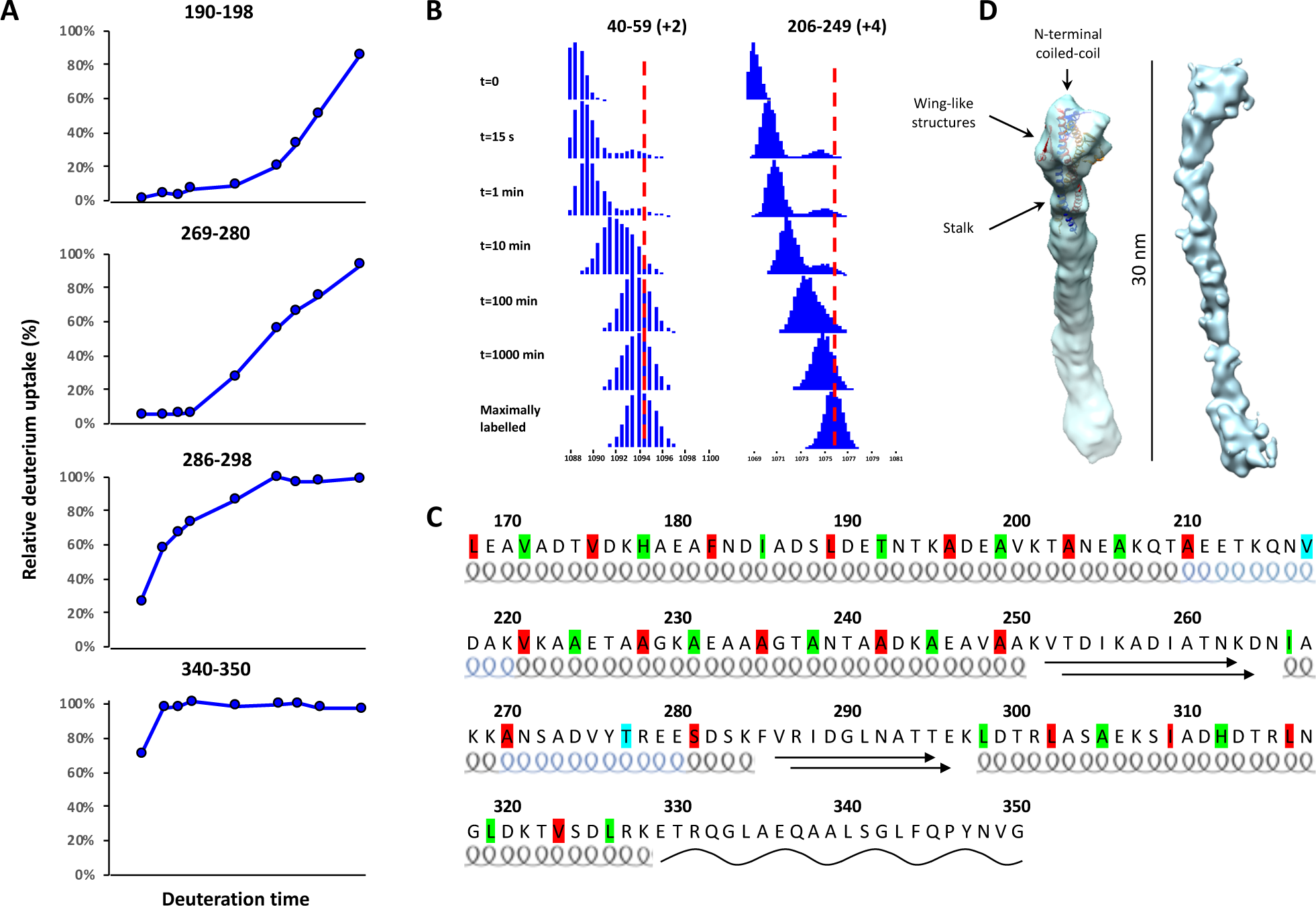
Structural information on rNadA. **A**). Deuterium uptake plots of representative peptides spanning distinct structural motifs of NadA and with distinct HDX rates (deuterium labelling times: 4 s, 15 s, 30 s, 1 min on ice and 1 min, 10 min, 30 min, 100 min, 1000 min at room temperature). Uptake values are normalized by the MaxD and express as % of deuteration. **B**) Bimodal HDX kinetics of NadA. Isotopic distributions of peptides 40-59 (coiled coil, head) and 206-249 (coiled coil, stalk) upon deuterium labelling at room temperature. The dashed red line indicates the centroid mass of the maximally labelled state. **C**) Proposed structural model of the stalk (aa 168-350). Black helices represent coiled coil heptad repeats, blue helices represent coiled coil undecad repeats, arrows indicate the flexible domains, and the black wavy line represents the disordered region of NadA tail. Residues colored in red are located in *a* position of heptad and undecad repeats, residues colored in green are located in *d* position in heptads, and residues colored in cyan are located in *h* position in undecads. **D**) Different views of the cryo-EM map of rNadA (left, front view; right, side view) fitted with the crystal structure of NadA24- 170 (PDB 6EUN).

### Conformational differences between rNadA and OMV-embedded NadA

Successively, we aimed at investigating NadA structure *in situ*, as exposed on the meningococcal membrane surface. To do so, we analyzed meningococcal OMVs overexpressing full-length NadA. In this native form, the passenger domain of the protein extends outwards from the beta-barrel translocator domain, which is anchored and inserted into the outer membrane^12^. By label-free proteomic analysis, 296 proteins were identified in the OMVs, with predicted cytoplasmatic proteins representing 5.5% of the total protein content, revealing the high quality of the OMVs. NadA monomer was measured at 8.7% of the total protein content, corresponding to 2.9% of trimeric NadA (table 3). In negative staining transmission electron microscopy (NS-TEM), the OMVs appeared as spherical and intact particles with a diameter ranging from 30 to 100 nm, surrounded by a thick membrane carrying several copies of NadA (**Fig. 2a**). NadA inserted onto the membrane appeared as an elongated appendage of approximately 30 nm in length, formed by flexible rod-like stalk decorated by a small globular head of 5 nm in diameter, projected toward the extracellular *milieu*. The stalk contains cavities and, interestingly, presents an enlargement close to the C-terminus (at around 20 nm distant from the head), which could be the coiled coil interruptions predicted by HDX-MS (**Fig. 2b**). The NadA tail is visible between the membrane-spanning region and the enlargement along the stalk. Furthermore, we compared the conformational dynamics of rNadA and OMV-embedded NadA by HDX-MS (**Figs. 2c and 3a**). We carried out an HDX-MS experiment with integrated size-exclusion chromatography (SEC-HDX-MS), which enabled efficient lipid removal and allowed monitoring the HDX of 36 peptides spanning 72.8% of sequence^35^ (table 4, uptake plots in Supplementary dataset 3). We observed that peptides of OMV-embedded NadA encompassing the insertion within residues 284-297 and the wings did not reveal significant differences in HDX compared to rNadA, indicating that these elements of flexibility are present both in the recombinant and native state. They could thus be pivotal structural elements. The tail of NadA, located upstream the beta-barrel anchor, showed characteristics of a structural reorganization *in situ*. In rNadA, peptide 327-340, spanning the tail, was found to be highly dynamic (fully exchanged at early time points), whereas in OMV-embedded NadA, we clearly observed that the peptide isotopic envelope was characterized by HDX bimodal behavior and overall reduced HDX (**Fig. 3b**). This phenomenon can be reasonably attributed to a structural rigidification of the NadA tail due to the proximity of the membrane anchor. Compared to rNadA, OMV-embedded NadA manifested reduced HDX in several segments of the stalk (within residues 78-93, 155-163, 172-211, 270-278) and in the head domain (residues 38-59) (**Figs. 2c and3a**), indicative of a long-range propagation of the membrane stabilizing effect. Like rNadA, also OMV-embedded NadA displayed bimodal HDX behavior in its head domain (see paragraph *Identification of an open trimer conformation in NadA*), and the observed reduced HDX was due to the low-mass distribution **(Fig. 3b)**. Considering the observed long-range propagation of the membrane stabilizing effects, we investigated whether there could be in NadA a bidirectional allosteric crosstalk between the head and stalk domains. We thus performed HDX-MS analysis of rNadA in complex to a known interactor hypothesized to bind to the head domain, Hsp90^36–38^. We confirmed that Hsp90 directly associates with the NadA head domain (residues 40-51 showed decreased HDX) and observed that its binding induced significantly increased HDX, meaning structural destabilization, in the NadA stalk (from residue 94) (**Fig. S5**, table 5, uptake plots in Supplementary dataset 4). This destabilization resulted pronounced until the predicted coiled-coil interruption at residues 252-265, whilst appearing highly diminished downstream. Taken together, these data corroborate that an allosteric crosstalk exists in NadA and suggest two mechanistic features of this adhesin: 1) the coiled coil stalk presumably bends, twists or rigidifies when both termini are subjected to stimuli, like the membrane constrain at the C-terminus or interactors to the N-terminal head domain, giving rise to a bidirectional head-anchor crosstalk; and 2) the predicted coiled-coil interruptions likely play a role in releasing the torsional stress of the trimeric stalk when this is perturbed.

**Fig. 2.**
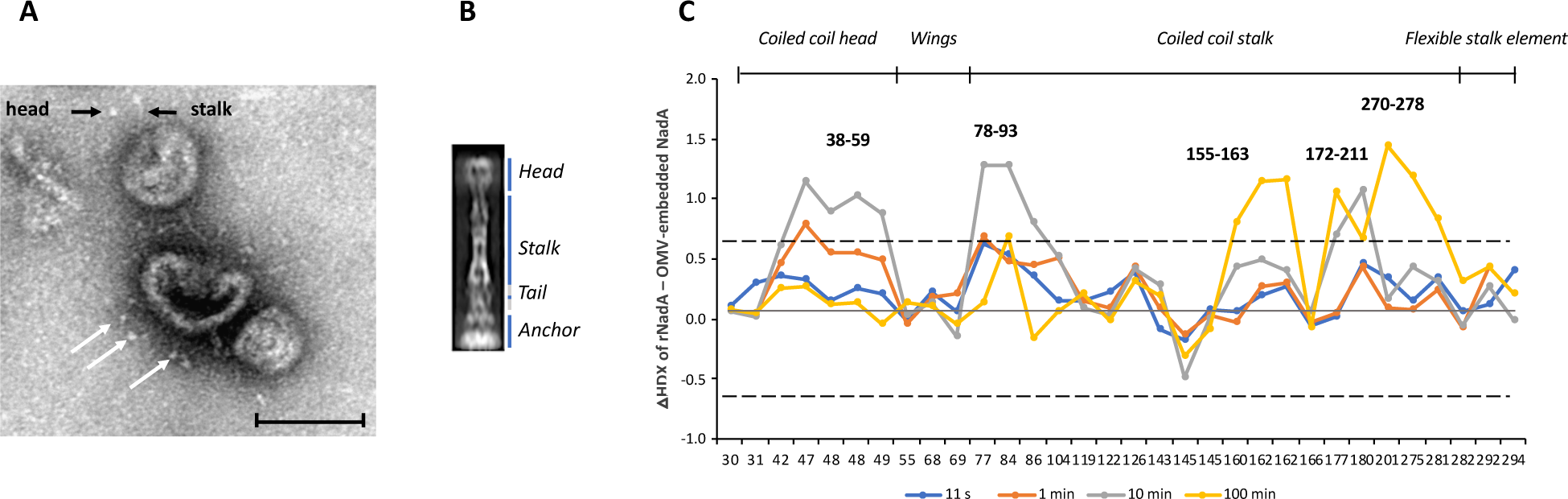
Structural information on NadA *in situ* (native NadA exposed on the OMV surface). **A**) Representative raw of Negative Staining TEM image of meningococcal OMVs carrying on the surface several NadA copies (indicated with white arrows). NadA head and stalk are marked with black arrows (scale bar 60 nm). **B**) Representative 2D class average of NS-TEM data showing a small portion of OMV membrane carrying NadA. Each class is composed by at least 100 individual boxed particles. **C**) Difference in HDX between rNadA and OMV-embedded NadA. The dashed black line indicates 0.61 Da (99% CI) as a threshold for statistical significance. On the x-axes, peptides are listed according to the peptide centre residue (peptides 24-39, 40-58, 40-59 and 327-340 are not plotted as they manifest bimodal HDX behavior; their isotopic envelopes are represented in fig. 3).

**Fig. 3.**
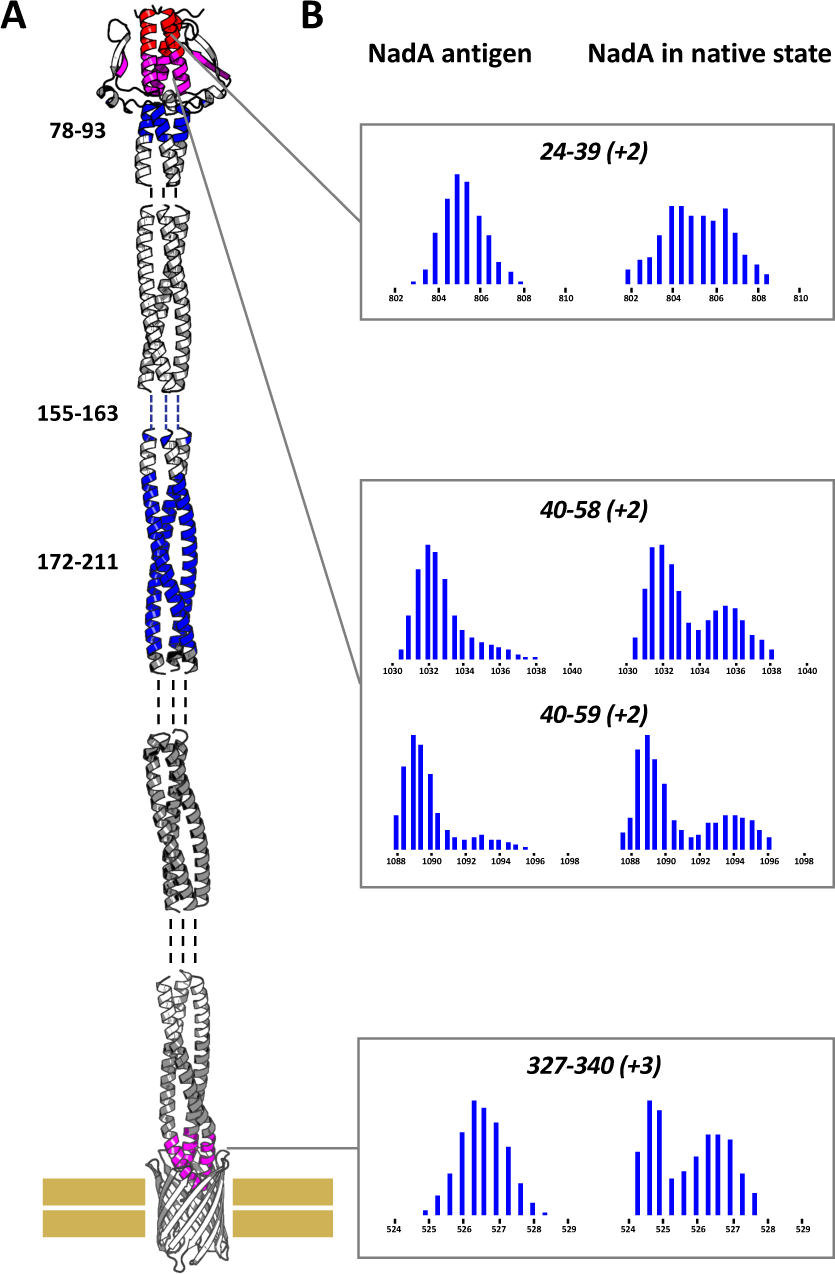
Bimodal HDX of NadA *in situ.* **A**) Differences in HDX between rNadA and OMV-embedded NadA are superimposed on the in-silico model of full-length NadA^28^. In red: HDX bimodality manifesting in OMV-embedded NadA only with overall increased HDX compared to rNadA. In purple: HDX bimodality in both states, with high mass envelope higher in OMV-embedded NadA and low mass envelope with decreased HDX; or bimodality with appearance of low mass envelope in OMV- embedded NadA. In blue: decreased HDX in OMV-embedded NadA compared to rNadA. In dark grey: no HDX information in OMV-embedded NadA. **B**) Illustrative isotopic distributions upon deuteration of peptide showings differences in HDX bimodality between rNadA and OMV-embedded NadA. From top to bottom: peptide 24-39 (+2), 11 s; peptide 40-58 (+2), 1 min; peptide 40-59 (+2), 11 s; peptide 327-340 (+3), 1 min.

### Identification of an open trimer conformation in NadA

As aforementioned, several regions of the coiled coil head and the stalk of rNadA exhibited bimodal isotopic envelopes (**Figs. S6 and S7**), which we clearly observed also in OMV-embedded NadA (**Figs. 3b and S8**). Bimodal HDX envelopes are characterized by a low- and a high-mass distribution distinguishable on the *m/z* scale, indicative of the presence of two different populations in solution: one in which the protein backbone amides are more prone to exchange compared to the other one^39^. In a pulse labelling HDX experiment on rNadA, both isotopic distributions displayed unvaried centroid mass and intensity, disproving irreversible unfolding or dissociation of the trimer in solution over time (**Fig. S9**). For rNadA, the two populations are not observed to interconvert during the HDX time course studied, indicating them as conformationally distinct, with a very slow interconversion rate. For OMV-embedded NadA, the two populations do show a minor interconversion during the HDX time course thus indicating a somewhat faster interconversion rate^40^ (**Figs. S6 and S7**). The alternative more accessible population mainly manifested in coiled coil regions flanking flexible elements, i.e. in proximity of the discontinuity between 252-265 of the stalk (peptides 206-249, 254- 274, **Figs. S2 and S7**) and of the wings in the head domain (peptides 36-58, 38-59, 40-59, 40-50, 74- 93, **Figs. S2 and S6**). These data denote that these flexible elements give a certain degree of conformational freedom to the NadA trimer, yielding the coiled coil to occupy co-existing more closed (tight) and more open (relaxed) trimeric states. The analysis of a shorter form of NadA (NadA24-149), containing approximately 70% of dissociated monomeric form^14^, suggested that the open trimer conformation in rNadA does not adopt a fully monomeric state, as the high-mass distributions of rNadA display lower centroid masses compared to NadA24-149 at early time points (**Fig. S6**). This experiment also rules out the presence of dissociated monomers in the rNadA sample, given that NadA24-149 manifested a clear HDX bimodality in the coiled coil segment downstream of residue 87, unlike rNadA. In rNadA, the closed and open trimer conformations (low-/high-mass distributions) appear to be distributed in approximately 9/1 ratio respectively (**Figs. S6 and S7**). Intriguingly, in OMV-embedded NadA, the HDX bimodal behavior of the head domain appeared to favor the high-mass distribution compared to rNadA (**Figs. 3b and S8**) and bimodal HDX was also observed in the extreme N-terminus (peptide 24-39), a phenomenon absent in rNadA. We could not confirm the presence of bimodal HDX in the stalk of OMV-embedded NadA because of the lack of sequence coverage, but we clearly show that the head domain of NadA *in situ* appears to manifest a more populated open trimer conformation compared to rNadA. Considering that the head domain is primarily responsible for binding as well as adhesion function and harbors the main epitopes^11^, it is reasonable to speculate that the open trimer conformation could influence the adhesive and immunogenic properties of NadA through expansion of its exposed surface.

### Evaluation of the immune response elicited by rNadA and OMV-embedded NadA

Owing to the conformational differences highlighted by the HDX-MS analysis, we compared the immunogenicity of rNadA and OMV-embedded NadA. CD-1 mice were immunized with 5 μg (total protein content) of MenB or *E. coli* OMVs overexpressing NadA or 20 μg of rNadA, the three formulated with aluminum hydroxide as adjuvant. MenB OMVs from a NadA knock-out strain (NadA KO) were used as negative internal control. The elicited antibody titers were evaluated by ELISA using post-III immune sera of individual mice. rNadA and both NadA-OMV-based formulations elicited comparable antibody titers that result significantly higher than the negative control (**Fig. 4a**).

**Fig. 4.**
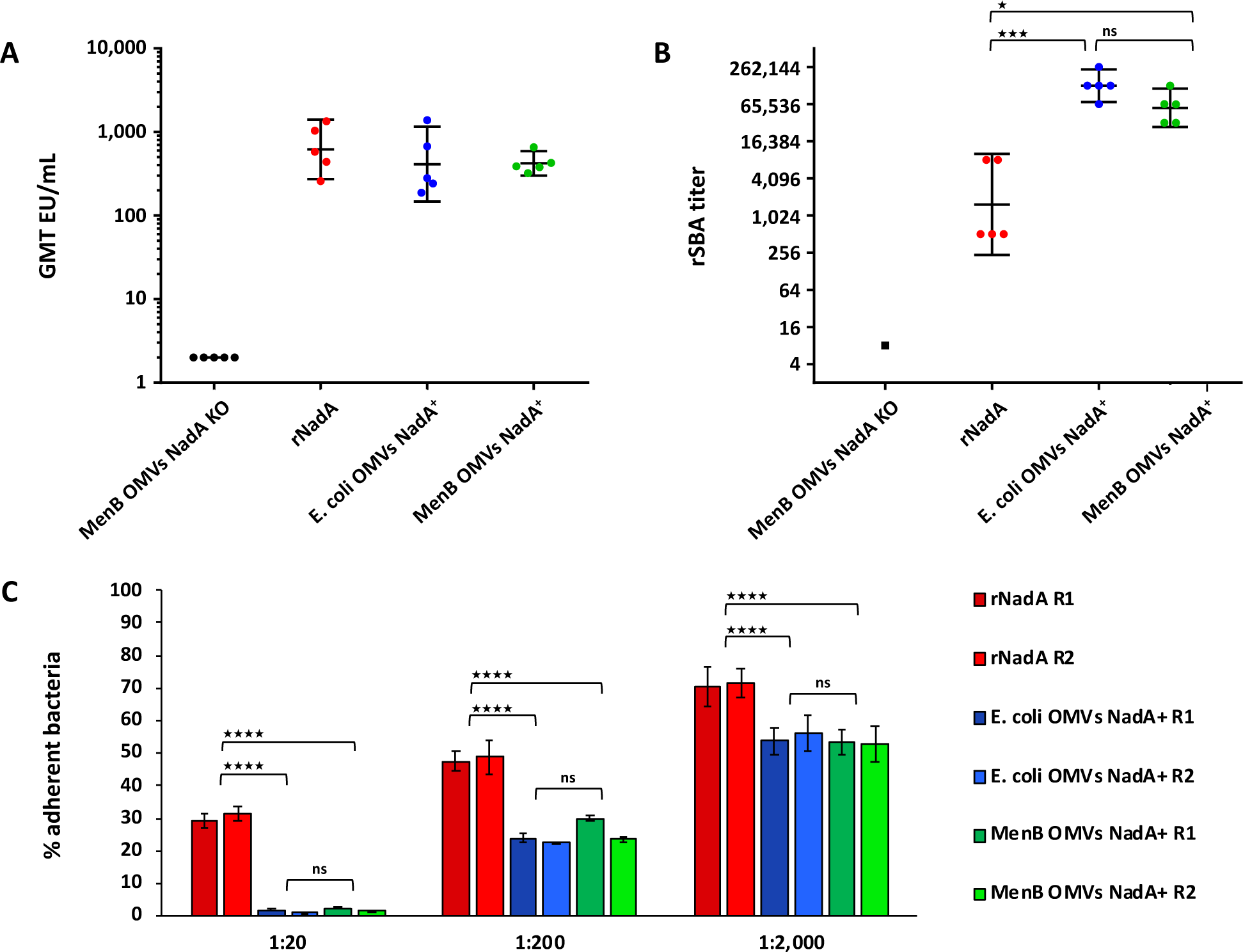
Immune response elicited by rNadA and OMV-embedded NadA. **A**) Antibody titers of sera from mice immunized with NadA-OMVs (5 µg) or with rNadA (20 µg). IgG α-NadA antibody titers elicited measured by ELISA, where each dot represents an individual mouse serum while lines indicate the median value within each immunization group. **B**) Serum bactericidal antibody (SBA) titers against BZ83 meningococcal strain. Dots represent SBA titers of individual mouse sera against the defined strain. Statistical analysis was performed using Kruskal-Wallis multiple comparisons test (ns: not significant; *p<0.05; ***p<0,001). **C**) Inhibition of NadA-mediated adhesion of the sera raised against rNadA and NadA-OMVs formulations in two biological duplicates (R1 and R2). Statistical analysis was performed using Two-way ANOVA followed by uncorrected Fisher’s LSD multiple comparison (****p<0.0001; ns: not significant).

To estimate the serum bactericidal activity (SBA) of the elicited antibodies, mice sera were tested against the BZ83 meningococcal natural strain. Expectedly, no killing was achieved with the serum derived from mice immunized with MenB OMVs from the NadA KO, whilst a highly functional response was achieved with sera derived from mice immunized with NadA-OMVs (SBA titers exceeding 65,000) compared to the response raised by rNadA (**Fig. 4b**). These data indicate that, even though the recombinant and OMV-based formulations elicit similar NadA-specific IgG titers, the OMV formulations show improved functional immune response and at lower dose of antigen administrated.

Additionally, we evaluated the activity of antisera raised against the distinct NadA formulations in a NadA-specific adhesion assays in which *E. coli* cells overexpressing NadA (*E. coli*-NadA^+^) exhibit NadA-mediated adhesion to Chang cell monolayers. Pooled mouse antisera from the rNadA or the OMV-NadA groups were pre-incubated with *E. coli*-NadA^+^ before adding to Chang cell monolayers to investigate their ability to inhibit bacterial adhesion, which is mediated by NadA head domain^12^ (**Fig. 4c**). Sera raised against MenB OMV NadA KO, as expected, did not show interference with bacteria adhesion and resulted in 100% adhesion (**Fig. S10**). Conversely, inhibition of bacterial adhesion was observed for both recombinant and OMV-NadA antisera and at the three dilutions tested (1:20, 1:200 and 1:2000), with significantly higher inhibition from sera obtained from OMV-NadA compared to rNadA at each dilution. These data suggest that NadA-specific antibodies elicited by the OMV formulations are more neutralizing for the adhesion function than the rNadA-elicited antibodies.

## Discussion

*In situ* structural biology of membrane proteins is still a largely unconquered area. Effective methods for the expression or extraction of membrane proteins as detergent-solubilized or completely soluble forms can lead to solving of their structure. Nevertheless, juxtaposing this information with their conformation in native state remains of paramount importance to elucidate the intricate interplay between their structure and function. However, the paucity of established methodologies for probing their structure and conformation *in situ* -in their native membrane environment- has so far hampered the acquisition of this knowledge. In this study, we compare the structural dynamics of the recombinant soluble meningococcal NadA and NadA *in situ,* revealing that, although overall very similar in structure, they show subtle conformational disparities that may influence epitope accessibility and molecular interactions with biological receptors.

The MenB NadA antigen, like other trimeric autotransporter adhesins, has posed several challenges to crystallization in its recombinant form, such that only half of the NadA structure has been determined by X-ray crystallography^14^. In this study, we applied HDX-MS to propose a structural model of the full-length NadA (NadA24-350) and analyze the conformational dynamics of NadA *in situ*, as natively exposed into OMVs, complementing the HDX data with Negative Staining and Cryo-EM imaging of the antigen. There are currently limited examples and no general consensus on how HDX-MS data are best used to provide structural information for modelling^41–44^. Here, we used a residue-level HDX approach in order to discriminate regions of stable helical structure (coiled coil regions) and disordered segments, based on the fact that detailed studies using NMR to measure individual amide HDX in proteins have indicated significantly lower HDX rates for amides in alpha-helical or beta-stranded regions compared to amides in unstructured regions^30^.

Using such an approach, we successfully modelled the region of the NadA stalk that lacks X-ray structure determination (downstream residue 168), and our results are compatible with fourteen coiled coil heptads, two undecad repeats and the presence of two flexible elements interrupting the coiled coil stalk. Notably, the segments predicted as undecad repeats were omitted in a previous *in-silico* model of NadA, as they indeed showed low heptad coiled coil probability^28^. Low resolution Cryo-EM data collected on the full-length rNadA antigen support the HDX-derived observation of a certain degree of flexibility of NadA stalk, since some bending at the level of the stalk region is clearly visible. So far, two other proteins of the trimeric autotransporter adhesins family (BpaA and EibD) have been observed with similar interruptions along the continuity of the stalk^13^. Although their function remains unknown, it has been proposed that these interruptions may be involved in releasing the constraint caused by local stalk twisting or that they may increase stalk flexibility by adding extra degrees of freedom^13^.

Additionally, we compared the conformations sampled by NadA antigen as soluble vaccine protein and when expressed on the surface of OMVs. NadA shows conformational heterogeneity in proximity of the wings in the head domain and of the predicted flexible interruptions of the coiled coil stalk, indicating that these elements give additional degrees of freedom to the trimer. Such a conformational heterogeneity is underlined by the presence of two distinct populations, corresponding to the expected tightly folded trimeric protein, and a more relaxed form likely manifesting upon transient opening (breathing) of NadA trimer. We observed that the open conformation is present to a low extent (10% of total population) in the recombinant NadA, whilst it is significantly more populated in the OMV state. We infer that the conformational constraint induced in the trimer by the anchoring to the bacterial membrane might be propagated along the entire stalk up to the distant head domain, with the constraint released at the level of the flexible elements enhancing trimer opening.

Similar allosteric effects were also identified, in the opposite direction, upon binding of Hsp90 to the NadA head, evidencing a bidirectional crosstalk along NadA stalk. In this case, a destabilization of the NadA stalk was observed up to the first flexible stalk interruption, whereas the effect in the downstream region was highly attenuated, highlighting again a role for the flexible stalk elements in releasing stalk torsional stress. So far, the role and the mechanism adopted by Hsp90 in binding NadA has not been fully clarified. While the chaperone function of Hsp90 in remodeling unfolded proteins or binding to intrinsically disordered regions in the cellular cytosol is fully elucidated^45–47^, the establishment of its presence in the extracellular space is more recent and functionally less defined. Extracellular Hsp90 has been found to be involved in the entry of some viruses^48^ as well as to modulate rNadA internalization and trafficking in epithelial cells and monocytes/macrophages stimulation^36–38^. It cannot be excluded that NadA conformational changes and dynamic behavior of the stalk upon binding to Hsp90 herein described could contribute to NadA-mediated cell entry.

Overall, all the structural data collected in this work highlight that NadA is conformationally heterogeneous, therefore we wondered whether the observed transition from a closed to an open trimer conformation, resembling a ‘molecular breathing’, could have functional implications. Although poorly explored for bacterial antigens, the concept of trimer breathing is not completely new and is reminiscent of a trimer-monomer equilibrium described for many viral antigens, i.e. HIV Env gp120/gp4^49^, RSV glycoprotein F^50^, SARS-CoV-2 spike^51^. It has been demonstrated that conformational switches contribute to virus receptor binding, whereas more drastic conformational changes (i.e. transition from pre-fusion to post-fusion state) promote virus-host cell membrane fusion. Notably, for viral proteins, conformation-specific antibodies for the open trimer and antibodies able to promote trimer opening have been identified^54–56^, leading us to hypothesize that similar mechanisms of antibody engagement could also be adopted for the bacterial trimer NadA. Therefore, we evaluated the functional immune response elicited by NadA in its recombinant form and when embedded in OMVs, in the attempt to rationalize the impact of the observed open and closed states of the head domain on its functional activity.

The NadA head domain is an important antibody target^11^ and is responsible for the adhesion function^28^. By comparing the immune response elicited by recombinant and OMV-embedded native NadA, we demonstrated that antibodies elicited by the native NadA showed higher functional activity. We cannot exclude that the increased bactericidal activity in the OMV formulations may be due to a shift in specific NadA IgG subclasses due to the adjuvant effect of LPS^52,53^ and particularly by an increase of IgG2a, which in mice has higher complement fixing effector activity^54^. However, the superior capacity of antibodies raised against OMV-embedded NadA of inhibiting bacterial adhesion to epithelial cells shows that native NadA elicits better neutralizing antibodies, suggesting that the more populated open trimer conformation of the native protein could underlie an expansion of its antigenic surface *in situ*, and exposure of epitopes that are otherwise partially occluded in rNadA. An enhanced immunogenicity has been analogously observed when NadA is genetically fused to *H. pylori* ferritin^55^, suggesting that locking the NadA extremity via its C-terminus to ferritin nanoparticle, as in the case of the bacterial membrane, improves its functional immune response. This strengthens the hypothesis of a correlation between the higher abundance of the open trimer conformation in the head domain of OMV-NadA antigen and the higher neutralizing functional response.

Taken together, our work represents a first attempt to explain differences in functionality of antibodies raised against recombinant and native bacterial antigens by changes in antigen conformational dynamics. Indeed other studies have reported higher efficacy of OMV-embedded antigens compared to purified recombinant proteins formulated with alum^56–59^. To note, further studies are necessary to unambiguously demonstrate whether the dynamic behavior of NadA and its breathing at the cell surface may be functionally relevant to promote MenB-host cell adhesion, similarly to what previously observed in the context of viral infections. Additionally, epitope mapping studies with monoclonal and polyclonal antibodies elicited after immunization with rNadA and OMV-embedded NadA will clarify the plasticity of the humoral response and assess the relationship between the transient opening of NadA trimer on the cell surface and immunogenicity. All together, these studies will be useful to unveil whether engineering strategies aiming at relaxing recombinant trimer antigens and promoting the open form may be considered for future vaccine design.

In general, our data indicate that it is possible to obtain structural insights on vaccine antigens *in situ*, in their native bacterial environment. HDX-MS delivered precise information on the conformational dynamics of NadA as displayed on the meningococcal surface, overcoming the extreme complexity of the bacterial membrane. The structural information is paralleled with antibody functional data of the humoral response elicited by NadA when in its native conformation, as that assumed when embedded in OMVs. Our data support the potential of OMVs as an antigen-presenting platform and highlight the importance of structural knowledge of antigens to drive vaccine design.

## Data availability

According to the community-based recommendations^60^, to allow access to the HDX data of this study, the HDX data tables and the proteomics data table are included as Supplementary tables 1–5, and all the deuterium uptake plots and the *k*HX plots are included as Supplementary datasets 1-4.

## Material and methods

*Preparation of OMVs*. The meningococcal OMVs were prepared from 5/99 strain (PorA subtype P1.5,2 and ΔlpxL1, ΔSynx,) deleted of NadA repressor (ΔNadR) to overexpress NadA^61^. The *N. meningitidis* strain was first pre-inoculated into 7 mL of Meningitidis Chemical Define Medium I-medium (MCDMI) at an OD600 ranging from 0.15-0.2 and incubated at 37 °C at 180 rpm until mid-exponential phase (=0.8-0.9 OD). The mid-exponential pre-cultures were inoculated into 50 mL medium in 250 mL baffled flasks and grown over night (16-18 h) until late stationary phase at 37°C and 180 rpm. Successively, the bacterial cells were pelleted by centrifugation for 30 min at 3,400 × g and 4°C, and the culture supernatants filtered (pore size 0.22 µm). OMVs were collected from the filtered supernatants by ultracentrifugation for 2 h at 9,6000 × g and 4 °C, and an additional washing step with PBS was performed. The OMV pellets were re-suspended in PBS at a concentration of 1 µg/µL.

*Identification of rNadA peptides*. Peptide identification was performed by liquid chromatography– tandem mass spectrometry (LC-MS/MS) analysis of peptides generated upon on-line pepsin digestion, setting buffer conditions and chromatographic gradient as for deuterated samples. Peptide ions were fragmented by collision-induced dissociation using argon as collision gas and MS/MS spectra were acquired in data independent acquisition mode (MS^E^). MS/MS data were processed with ProteinLynx Global Server (PLGS) version 3.0 (Waters). DynamX version 3.0 (Waters) was used to filter peptides by selecting those fragmented at least 2 times, presenting 0.2 fragments per amino acid, and found at least in 4 out of 5 acquired MS/MS files, to ensure confident peptide identifications. Additionally, the MS trace of the identified peptides was visually inspected and peptides with insufficient signal-to-noise ratio were discarded. The identification of the peptides for OMV- embedded NadA was not performed due to the lack of sufficient material.

*Continuous labelling of rNadA*. rNadA was diluted in PBS at room temperature or on ice and the temperature let equilibrate for 30 min prior labelling. HDX was then initiated by 4.17-fold dilution with deuterated PBS buffer (at 95% D2O), yielding to 72.2% of final deuterium content in the reaction mixture. The labelling was performed at room temperature (for 1 min, 10 min, 30 min, 100 min, and 1,000 min) and on ice (for 4 s, 15 s, 30 s, 1 min). At selected time intervals, an aliquot was withdrawn and quenched 1:1 with ice-cold 300 mM phosphate buffer at 4 M Urea (final pH/D 2.5). After quenching, samples were immediately frozen in liquid nitrogen and stored at -80 °C until LC-MS analysis. The labelling for 30 min at room temperature and 1 min on ice was performed two additional times, producing triplicates. A maximally labelled sample was associated to this experiment.

*Pulse labelling of rNadA*. rNadA was diluted in PBS and let in a thermomixer set at 23°C. At various incubation times (15 s, 1 min, 10 min, 100 min, and 1,000 min), an aliquot was withdrawn and HDX was initiated by 4.17-fold dilution with deuterated PBS buffer (at 95% D2O), yielding to 72.2% of final deuterium content in the reaction mixture. After 15 s of deuterium labelling, the reaction was quenched 1:1 with ice-cold 300 mM phosphate buffer at 4 M Urea (final pH/D 2.5). After quenching, samples were immediately frozen in liquid nitrogen and stored at -80 °C until LC-MS analysis. A maximally labelled sample was associated to this experiment.

*Continuous labelling of rNadA and rNadA24-149*. rNadA and rNadA24-149 were diluted alone in PBS and let in a thermomixer set at 23°C for 30 min. HDX was then initiated by 4.17-fold dilution with deuterated PBS buffer (at 95% D2O), yielding to 72.2% of final deuterium content in the reaction mixture. At various time intervals (15 s, 1 min, 10 min, 100 min, and 1,000 min), an aliquot was withdrawn and quenched 1:1 with ice-cold 300 mM phosphate buffer at 4 M Urea (final pH/D 2.5). After quenching, samples were immediately frozen in liquid nitrogen and stored at -80 °C until LC-MS analysis. A maximally labelled sample was included in the data set.

*Continuous labelling of rNadA and rNadA-Hsp90*. rNadA was incubated alone in PBS or with Hsp90 (ratio 1:3.8 NadA: Hsp90) for 30 min at room temperature or on ice, in order to allow the complex to form and the temperature to equilibrate prior labeling. HDX was then initiated by 4.17-fold dilution with deuterated PBS buffer (at 95% D2O), yielding to 72.2% of final deuterium content in the reaction mixture. At various time intervals (15 s, 1 min, 10 min, 30 min, 100 min, and 1,000 min at room temperature; 15 s, 30 s, 1 min on ice), an aliquot was withdrawn and quenched 1:1 with ice-cold 300 mM phosphate buffer at 4 M Urea (final pH/D 2.5). After quenching, samples were immediately frozen in liquid nitrogen and stored at -80 °C until LC-MS analysis. The labelling for 30 min was performed two additional times, producing triplicates for that time point. A maximally labelled sample was associated to this experiment.

*Continuous labelling of rNadA and OMV-embedded NadA*. rNadA and OMV-containing NadA were diluted in PBS and let in a thermomixer set at 23°C for 30 min prior labelling. HDX was then initiated by 2.33-fold dilution with deuterated PBS buffer (at 95% D2O), yielding to 66.5% of final deuterium content in the reaction mixture. At various time intervals (11 s, 1 min, 10 min and 100 min at 23°C), an aliquot was withdrawn and quenched 1:1 with ice-cold 300 mM phosphate buffer at 4 M Urea (final pH/D 2.5). After quenching, samples were immediately frozen in liquid nitrogen and stored at -80 °C until LC-MS analysis. The labelling for rNadA was performed in triplicates for each time point.

*Maximally labeled control*. rNadA was diluted in Solvent A (0.23% formic acid), injected onto the LC system and digested on-line. The solvent eluting from the pepsin column, containing peptides generated upon enzymatic cleavage, was collected and evaporated to dryness. Peptides were resuspended in deuterated PBS buffer at equal deuterium content as in the protein labelling mixture and the exchange was allowed for 2 h. The reaction was quenched 1:1 with ice-cold 300 mM phosphate buffer at 4 M Urea (final pH/D 2.5). Samples were immediately frozen in liquid nitrogen and stored at -80 °C until LC-MS analysis.

*Liquid chromatography method for soluble recombinant proteins.* Frozen quenched samples were rapidly thawed with a bench top centrifuge and injected onto a refrigerated UPLC system (NanoAcquity, Waters) with all chromatographic elements held at 0 °C. Except for the maximally labelled peptides, all protein samples passed through an in-house packed column containing pepsin immobilized on agarose resin beads at 20 °C. The generated peptides were trapped on a Vanguard column (BEH C18, 130 Å, 1.7 µm, 2.1 mm × 5 mm; Waters) and desalted for 2.5 min in solvent A (0.23% formic acid, pH 2.5) at a flow rate of 200 µl/min. Subsequently, the peptides were separated over an Acquity UPLC column (C18, 130 Å, 1.7 µm, 1.0 mm × 100 mm; Waters), with a 9 min-linear gradient rising from 8 to 40% of solvent B (0.23% formic acid in acetonitrile, pH 2.5) at 40 µl/min.

*Size-exclusion chromatography coupled to liquid chromatography method for rNadA and OMV- embedded NadA*. Frozen quenched samples were rapidly thawed and injected onto the refrigerated UPLC system with all chromatographic elements held at 0 °C. Protein and protein-lipid samples were analyzed by modifying the LC apparatus with the integration of a short size-exclusion chromatography (SEC) column in the HDX refrigerated chamber, placed after the pepsin column^35^. Samples were digested through a pepsin column held at 20 °C, then filtered through the SEC column (BEH SEC guard column, 125 Å, 1.7 µm, 4.6 x 30 mm, 1-80K; Waters). OMV lipids, when present, were retained in the SEC, whereas peptides passed and were trapped and desalted on a C18 Vanguard column for 4 min in solvent A, at a flow rate of 150 µl/min. Subsequently, peptides were separated over a C18 analytical column as described in the previous session. The SEC was cleaned on-line during the chromatographic gradient with four subsequent injections of 100 µL of 6M Urea in phosphate buffer (pH 2.5), whereas the pepsin column was removed, and a metal union mounted in its place.

*Mass spectrometry method*. Following the chromatographic separation, peptides were analyzed using a hybrid ESI-Q-TOF mass spectrometer (Synapt G2-Si, Waters). The instrument was set in positive ionization mode with spray voltage of 3 kV and the ions were further separated by ion mobility for enhancing peak capacity. The MS spectra were acquired in range from 200 to 2000 m/z, performing a scan every 0.5 s. Human Glu-fibrinopeptide (Sigma-Aldrich, St. Louis, MO) served as an internal standard and the peptide signal was recorded throughout all the analysis.

*Data analysis*. DynamX version 3.0 was used to calculate deuterium incorporation at peptide level. LogkHX values were calculated with HDsite software, by applying the centroid fitting and normalizing the data for the maximally labelled control. Time points of the labelling on ice (≌0°C) were normalized to in-exchange time intervals at room temperature as elucidated in^62^.

*Statistical approach*. The statistical significance of a difference in HDX between two states was determined by calculating a confidence interval (CI) based on the pooled standard deviation of the peptide deuterium uptake (SD) in time points performed in triplicates following Weis’ recommendations^63^. For each state, the SDs were pooled using the root-mean-square, according to equation 1:

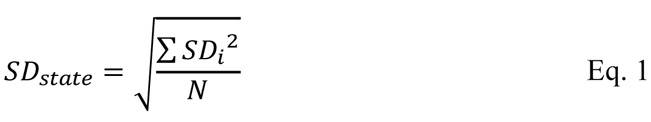

where 𝑁 is the number of peptides considered. A pooled SD for the states performed in triplicates, was subsequently calculated using equation 2 for the labelling rNadA-Hsp90:

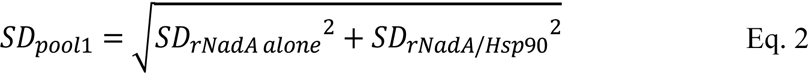

The restricted volume of OMV sample allowed to produce only singlets, while triplicates were carried out for the state rNadA at every time points. The propagated SD (SDpool) was thus calculated as follow:

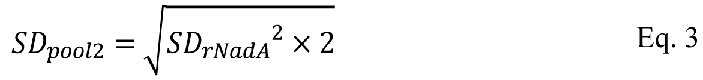

The pooled SD calculated by eq. 2 and 3 were used to calculate the CI at significance level of 98% and 99% respectively, with a zero-centered average difference in deuterium uptake considering a two-tailed distribution with two degrees of freedom (*n*=3), by using equation 4 and 5, respectively:

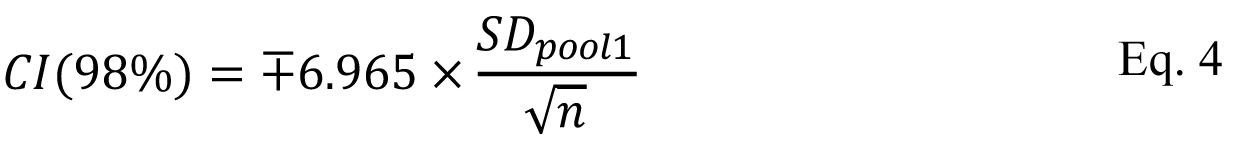

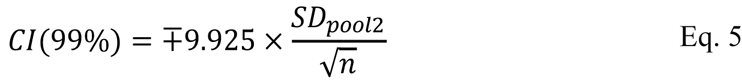

where n is the number of replicates performed, equal to 3. A higher CI was set for the comparison of rNadA/OMV-embedded NadA to compensate for the lack of triplicates in one state. Additionally, for peptides showing significantly different deuterium content between rNadA and OMV-embedded NadA, we checked that the ΔHDX was at least >5xSD calculated for rNadA in that time point (all peptides pass this threshold, and, in most instances, their ΔHDX is highly above this threshold).

*Label free proteomics analysis of the OMVs*. Five micrograms of OMV proteins were digested with trypsin using the iST kit (Preomics) according to the manufacturer’s protocol. Data acquisition was performed on a Q Exactive Plus (Thermo Scientific) mass spectrometer equipped with an electrospray interface (ESI) and coupled to an Acquity UPLC I-Class (Waters). Peptides were injected and separated on an Acquity UPLC CSH Peptide C18 130 Å column (1 mm X 150 mm, 1.7 μm, Waters) set at 50 °C using a flow rate of 0.05 mL/min in gradient mode (mobile phase A consisted of 3% (v/v) acetonitrile 0.1% (v/v) formic acid in water and a mobile phase B of 0.1% (v/v) formic acid in acetonitrile). The following gradient was used: 0–28% B in 40 min, 28–85% B in 5 min, holding at 85% B for 5 min and re-equilibration at 0% B for 10 min. Acquisition was performed in Data Dependent Acquisition (DDA) mode (Top5). Data base search was performed using Peaks software by Bioinformatics solutions, version X, using the public data base containing the 1950 annotated proteins of the reference strain NZ-05/33 of *Neisseria meningitidis*. LFQ was performed using the “quantification Peaks module”. The identified proteins were first classified as lipoproteins using Signal P^64^ and the cellular localization of the other proteins was assigned by PsortB 3.0^65^. For quantification, the measured proteins were normalized respect the total amount of identifiable proteins and expressed as percentage of the total protein amount.

*Negative staining electron microscopy.* OMVs diluted at 80 ng/μL in PBS were loaded onto a copper 200-square mesh grid of carbon/formvar rendered hydrophilic by glow discharge (Quorum Q150R S). The excess solution was blotted off after 30 s using Whatman filter Paper No.1 and then the grids were negatively stained with NanoW (Nanoprobes) for 30 s and then blotted using Whatman filter Paper No.1 and let air dry. Micrographs were acquired using a Tecnai G2 Spirit Transmission Electron Microscope (final magnification of 120000x), equipped with a CCD 2kx4k camera. For data analysis, a total number of 20 NS-TEM images were collected by using Relion 3.0 software and 800 individual NadA particles selected on the OMVs surface, by a box of 200 pixel (3.84 Å/pixel used for processing). After performing 2D classification. The best class average, containing 128 images, was selected for preliminary structure understanding.

*Cryo-electron microscopy.* Data collection was performed at the FEI (The Netherland). Purified recombinant NadA variant 3 concentrated 1 mg/ml was diluted 1/15 times to a final concentration of 0.067 mg/ml with buffer composed of 20 mM Tris and 150 mM NaCl, pH 7. Quantifoil R2/2 grid (400 mesh Cu, Electron Microscopy Sciences) was rendered hydrophilic with 15 mA current for 90 second by glow discharge in a EmiTech K100X. 2.5 µl of the specimens were deposited onto the grid and vitrified using a Mark IV Vitrobot (FEI Company) with a blotting time of 4 second and humidity of 100% at 4°C. The plunge- frozen samples were imaged with a Titan Krios electron microscope operating at 300kV in focus with a Volta Phase plate with focus determined manually at a single offset position. A prototype Falcon 3 detector was used at 47,000 X nominal magnification with a pixel size of 1.75 A/pixel. A total dose of 120 electrons/Å^2^ was applied to each image space. The image processing was performed within Scipion Software platform (http://scipion.cnb.csic.es)^70^, which provides several 3DEM software packages unified into an interface. Particles in the images presented a clear preferred orientation with mainly lateral views, due probably to the length of the molecule that exceeds the ice thickness. Moreover, due to the probable difficult classification of “heads” and “tails”, the particle picking was performed on half of the particles and 15273 “heads- tails” were extracted. The particles were classified several times to prune the set in 2D using Relion^71^ and the best classes obtained were used to generate an initial model within EMAN2^72^.This map was subjected to 3D classification in Relion resulting in two different models with distinct features: one with a globular extremity and a upright stalk while the other presented a straight part ended with a broaden bent part, presumably corresponding to the head and the tail, respectively. Particles were cleaned again using reference free 2D classification in Relion and the best 6270 particles were re extracted from the original micrographs with a bigger box size of 320 pixels in order to obtain the full-length protein. Since the recombinant NadA full-length was known, an initial model of the correct length was created using Chimera program^73^ joining the initial volume of the head and the tail and extending the central part and low filtered in order not to generate bias. This raw initial model was 3d classified in Relion against the latest set of 4439 particles generating a structure of around 25 Å calculated using FSC=0.143 criteria^74^. To improve the resolution of this model 7327 particles were manually selected with a smaller box size of 240 pixels to avoid an overlapping of particles in the same box. This latest dataset was pre-processed and 3D classified using Relion to generate two maps and a new set of 5735 particles formed by the best ones that were later re-extracted with improved centering to meliorate the alignment of the particles. The best 3D volume obtained from classification was processed with Xmipp3 tools to be filtered map and to create a tight soft mask. Finally, this masked density was used as an initial template for the 3D refinement in projection matching with the 5735 particles. The final resolution reached 14 Å measured with FSC=0.143 criteria. Local resolution was calculated using MonoRes^75^ while the angular distribution was tested with Relion package. To confirm the overall structure a fitting with the crystal structure of NadAV5 (PDB: 6EUN) was performed using Chimera software removing the missing electron density part from the PDB.

*Mouse immunizations*. Animal studies have been approved by the Italian Ministry of Health and were ethically reviewed by the local Animal Welfare Body. Animal studies were carried out at GSK facility in accordance with national/European legislation and the GSK Policies on the Care, Welfare and Treatment of Animals. Groups of five between 6- to 8-week old female CD-1 mice (Charles River) were immunized intra-peritoneally (IP). For each injection, the mice received a total dose of 5 μg of OMV vaccines or 20 μg of NadA recombinant proteins. The OMVs or the recombinant protein vaccines were absorbed with aluminum hydroxide (Alum, 3 mg/ml) and administered in three doses at day 1, 21 and 35. Blood was taken at day 0, 20, 34 for further analysis and bleed out was performed on day 49.

*IgG antibody titers elicited against NadA*. Serum antibody titers against NadA were measured by ELISA, which was performed as described elsewhere^76^. Microtitter plates were coated overnight at 4°C with 0.015 μM of purified NadA recombinant protein variant 3. Plates were incubated with single mice sera followed by alkaline phosphatase-conjugated anti-mouse antibodies. After addition of p-nitrophenyl phosphate, optical density was analyzed using a plate reader at a dual wavelength of 405/620-650 nm. Antibody titers were quantified via interpolation against a reference standard curve.

*Serum Bactericidal Assay (SBA)*. Serum bactericidal activity against *N. meningitidis* strain BZ883 (PorA subtype P1.5,2) was evaluated as previously described^77^ with pooled baby rabbit serum (Cederlane) used as the complement source (rSBA). Bacteria were grown in Mueller Hinton broth (MH), plus 0.25% (w/v) glucose, at 37°C with shaking until early exponential phase (OD600 of ∼0.25) and then diluted in Dulbecco’s saline phosphate buffer (Sigma) with 0.1% (w/v) glucose and 1% (w/v) BSA (Bovine Serum Albumin) to approximately 10^5 CFU/ml. The incubation with 25% of baby rabbit complement with or without polyclonal mouse sera at different dilutions was performed at 37°C (or 30°C as specified for respective experiments) with shaking, for 60 min. Serum bactericidal titers were defined as the serum dilution resulting in a at least 50% decrease in the CFU/ml after 60 min of incubation of bacteria with the reaction mixture, compared to the control CFU/ml at time zero.

*Inhibition of the binding assay*. *E. coli* BL21(DE3)T1R (New England Biolabs) was used to express full-length NadA variant 3 following transformation with pET21-NadA, as previously described^12^, or the empty vector (*E. coli*-pET21) as control. *E. coli* was cultured at 37°C in Luria–Bertani (LB) broth supplemented with ampicillin (100 μg/mL) and chloramphenicol (10 μg/mL). Surface protein expression for NadA was achieved without addition of IPTG, exploiting the expression due to the leakage in the induction system. Chang epithelial cells (Wong-Kilbourne derivative, clone 1-5c-4, human conjunctiva, ATCC CCL-20.2) were maintained in Dulbecco’s modified Eagle’s medium (DMEM) supplemented with antibiotics and 1% heat-inactivated fetal bovine serum (iFBS). Cells were grown at 37°C in 5% CO2. Chang cells were seeded in a 24-well plate (1.5 x 10^5 cells/well) in antibiotics-free medium with 1% iFBS (Infection Medium) and were allowed to grow for 24 h. For adhesion analysis, *E. coli*-pET21 or expressing NadA were added at a multiplicity of infection (MOI) of 100 and allowed to adhere for 3 h at 37°C and 5% CO2. Unattached *E. coli* were removed by extensive washing, and bacteria were counted using serial dilution plating. The number of cfu obtained with *E. coli* expressing NadA in absence of sera was assigned to be 100% of adhesion while the percentage of cfu obtained infecting cells with *E. coli*-peT21 indicated the maximum level of inhibition. For inhibition of the binding assay, *E. coli* expressing NadA was pre-incubated in rotation with the sera at three serial dilutions (1:20; 1:200; 1:2,000) or infection medium alone for 1 hour at 4°C before addition to the cells. The experiment was performed twice and each time in triplicates.

## Acknowledgments

K.D.R. and N.N. would like to gratefully acknowledge funding for this project from the European Union’s Horizon 2020 research and innovation programme under the Marie Skłodowska-Curie grant agreement VADEMA No 675879. I.P. and I.F. thank Kasim Sader for Cryo-EM data collection, José María Carazo (Biocomputing Unit (CNB-CSIC, Madrid) for the valuable scientific conversations and Roberto Melero (Biocomputing Unit (CNB-CSIC, Madrid) for the technical support and discussion regarding Cryo-EM data processing.

## Author contribution

N.N., K.D.R. and I.D. conceived the study. V.C. performed HDX-MS experiments. V.C., L.D.I., Z.Y.K., K.D.R. and N.N. analyzed HDX-MS data. S.B., E.L., A.B. and B.B. performed functional assays. I.F., I.P. and F.G. performed electron microscopy experiments. L.E.F. performed proteomics analysis. Z.S. and M.M. provided Hsp90 and OMVs. V.C., L.D.I., S.B., I.F., F.G., L.E.F., M.M., I.D.,

K.D.R. and N.N. wrote the manuscript with inputs from all authors.

---

**Fig. S1.**
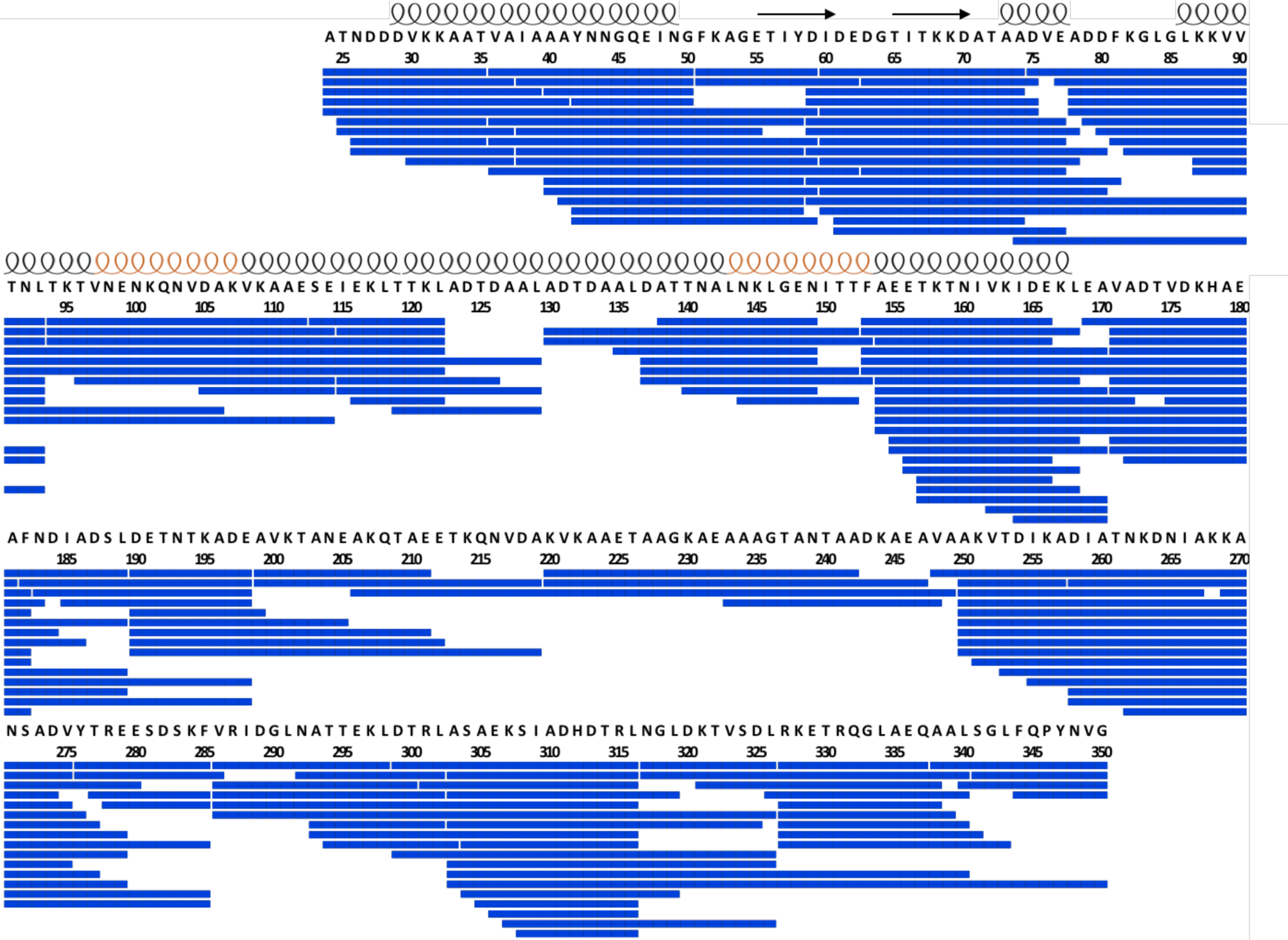
Peptide whose HDX was followed for the analysis of rNadA. The secondary structural elements determined from crystal structure of NadA24-170 (PDB: 6EUN) are depicted along the protein sequence: black helices represent coiled coil heptad repeats, orange helices represent coiled coil undecad repeats, arrows indicate the wing-like structures of the head domain.

**Fig. S2.**
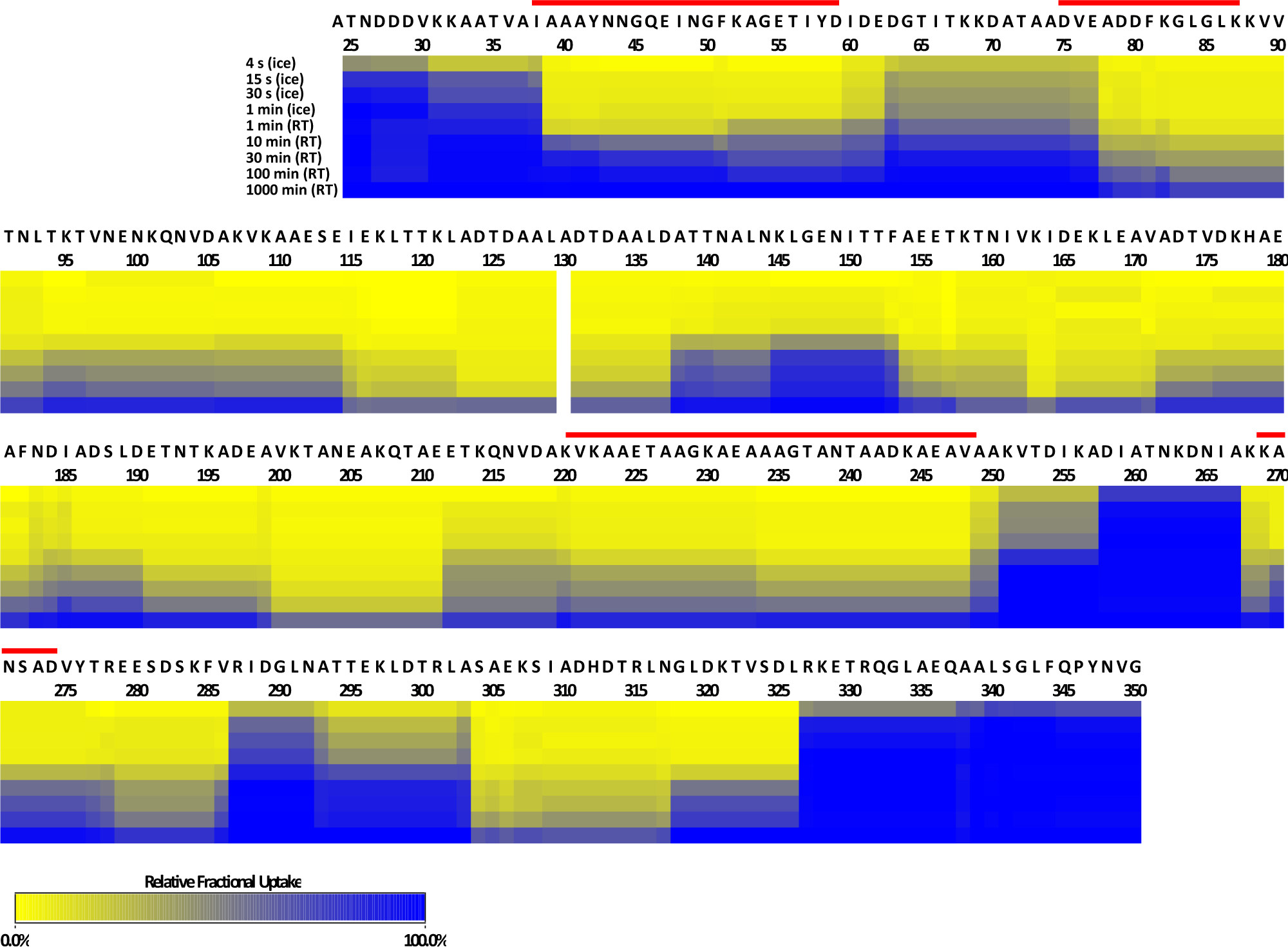
Relative fractional deuterium uptake of rNadA. The direct outcome (heat map) of DynamX v. 3.0 is reported. The scale indicates as 100% the relative uptake of the maximally labelled state, and as 0% the uptake of the undeuterated reference. The fractional uptake of individual peptides at individual time points has been normalized by the maximally labelled control. For the heat map, DynamX superimposes the information on uptake of intense peptides to that of less intense peptides and does not subtract the fractional uptake of overlapping peptides. Therefore, the heatmap represents an approximate map of the local HDX of rNadA (residue-level HDX information has been deconvolved by HDsite and is shown in fig. S3 and table 2). Red line indicates the regions manifesting bimodal HDX behavior.

**Fig. S3.**
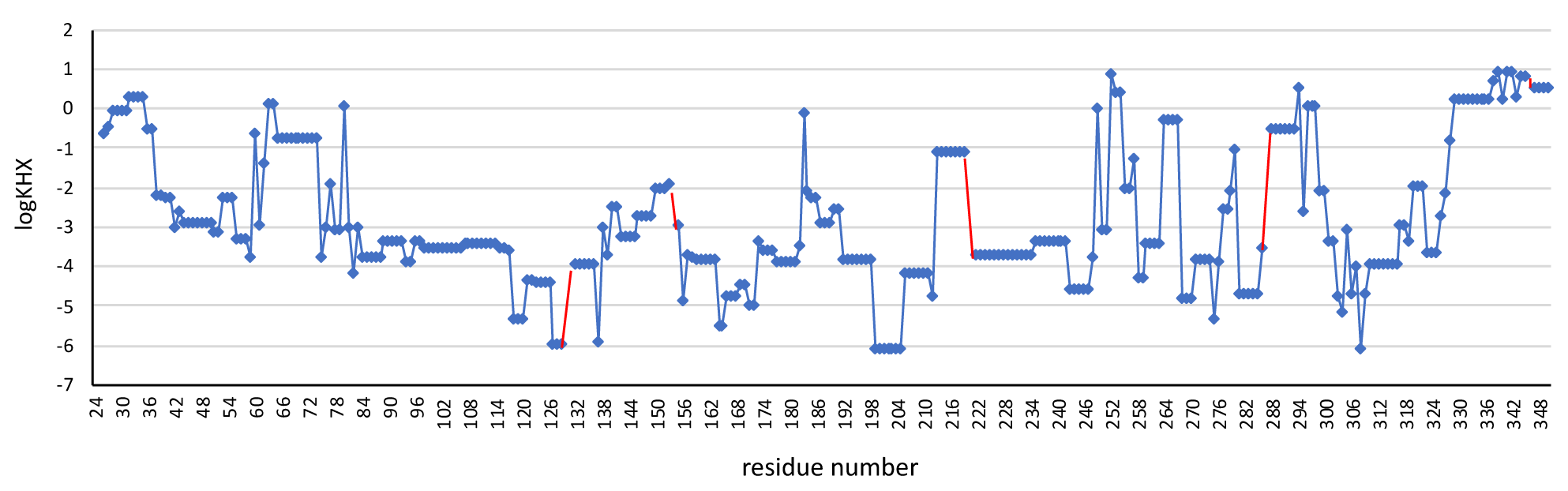
Log*K*HX values calculated by HDsite are plotted as blue dots along the protein sequence of rNadA. Red lines indicate missing values. On the x-axes, the residue number from 24 to 350.

**Fig. S4.**
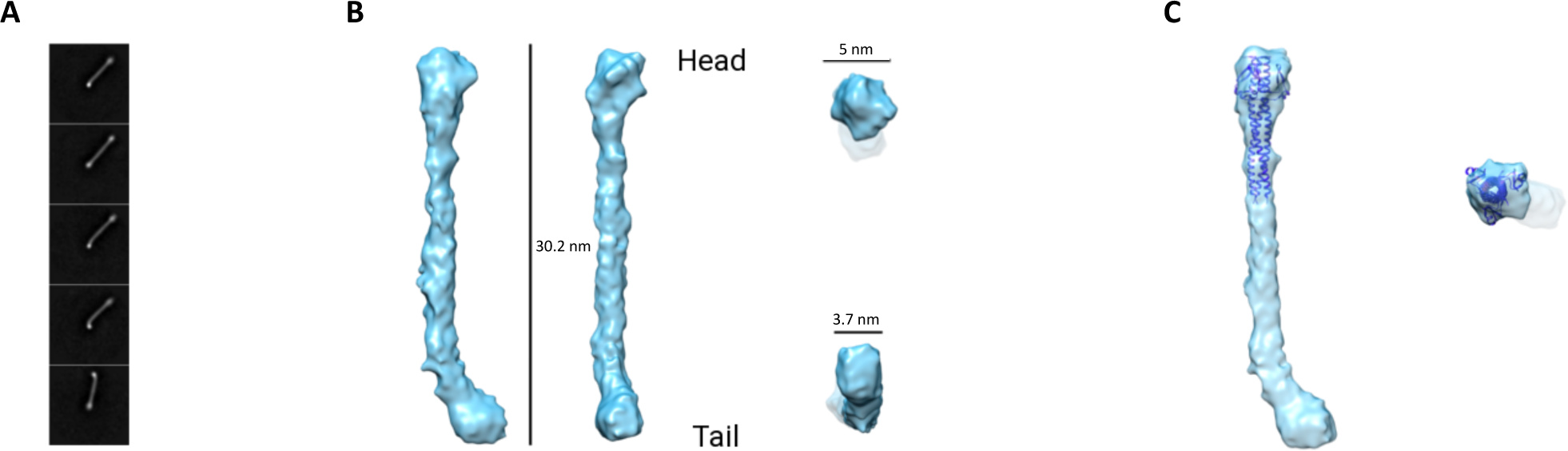
Cryo-EM map of rNadA. A) 2D class averages of rNadA. B) Different views of the Cryo-EM map of rNadA. C) Cryo-EM map of rNadA fitted with the crystallographic structure of NadA24-170 (PDB: 6EUN), top and lateral views.

**Fig. S5.**
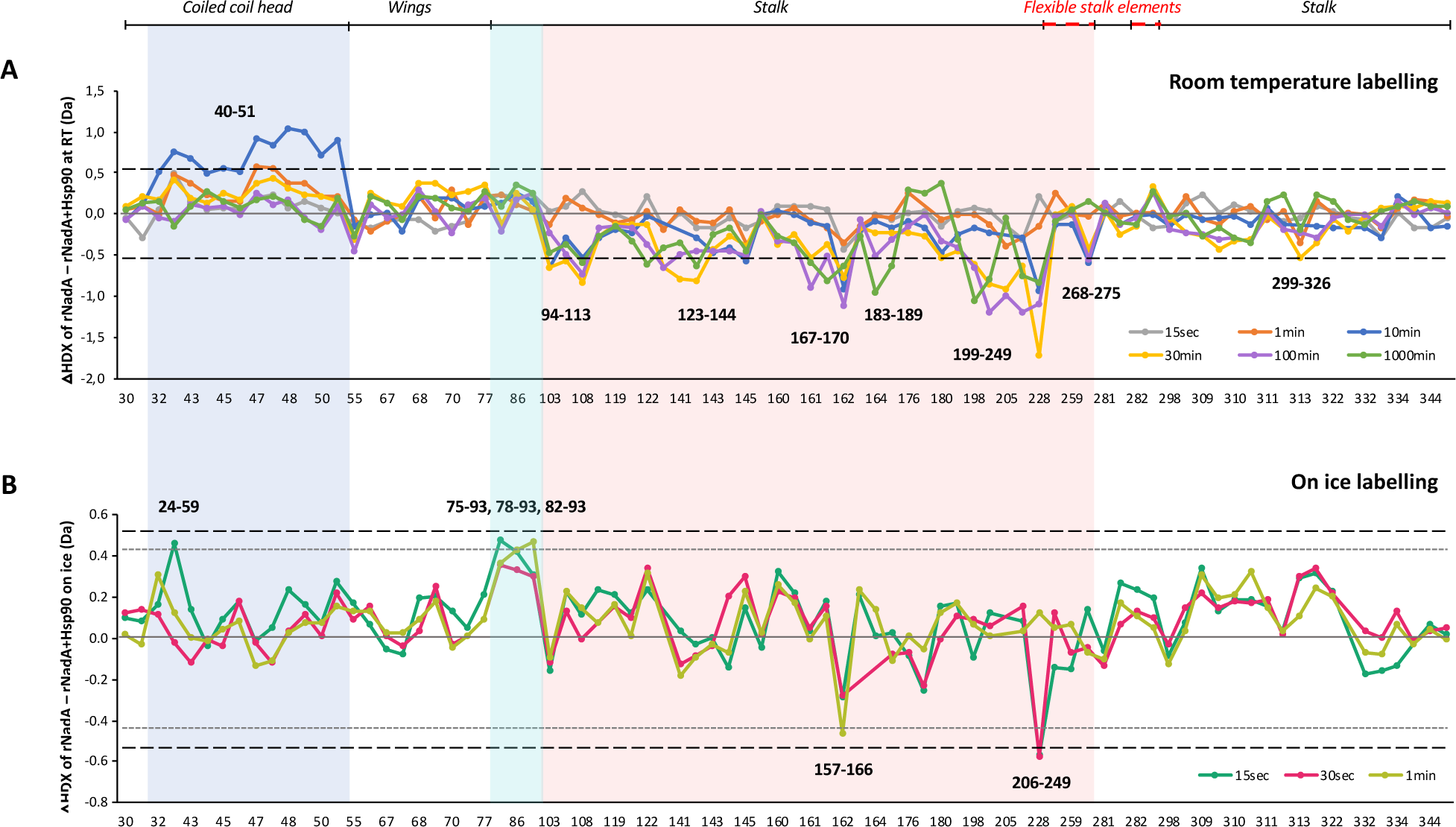
Conformational impact of Hsp90 on rNadA. A) Difference in HDX between rNadA alone and in complex with Hsp90 labelled at room temperature (RT). The dashed black line indicates 0.52 Da (98% CI) as a threshold for statistical significance. B) Difference in HDX between rNadA alone and in complex with Hsp90 labelled on ice. The dashed black line indicates 0.52 Da (98% CI) as a threshold for statistical significance. The dot grey line indicates △HDX=0.43 Da, considered relevant effects on ice. On the x-axes, peptides are listed according to the peptide centre residue.

**Fig. S6.**
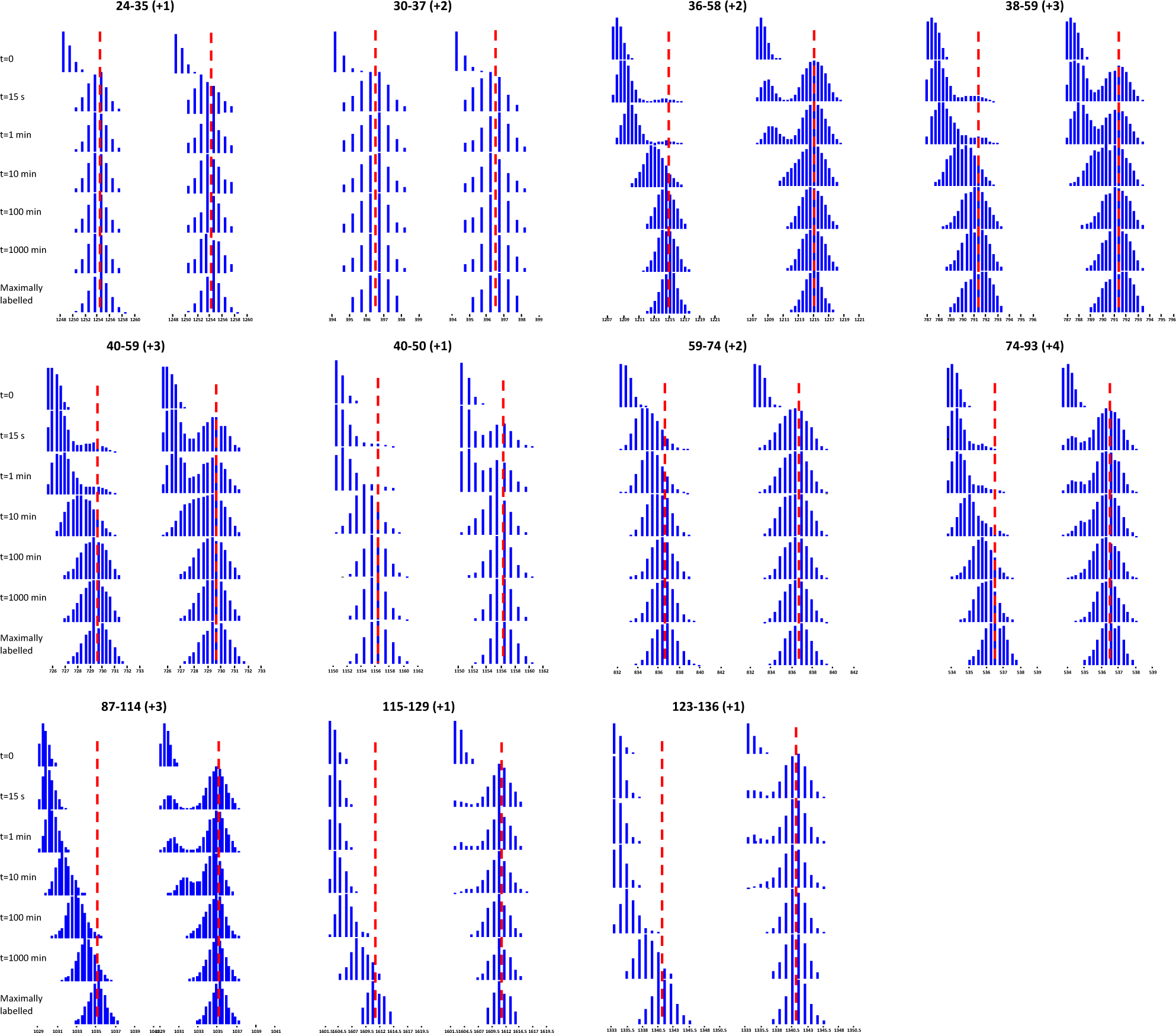
Bimodal HDX behavior of NadA (from aa 24 to 136). Illustrative isotopic distributions of peptides spanning sequence 24-136. Panels on the left show the spectra of rNadA; panels on the right show the spectra of NadA24-149 (partially trimeric form). The dashed red line indicates the centroid mass of the maximally labelled state.

**Fig. S7.**
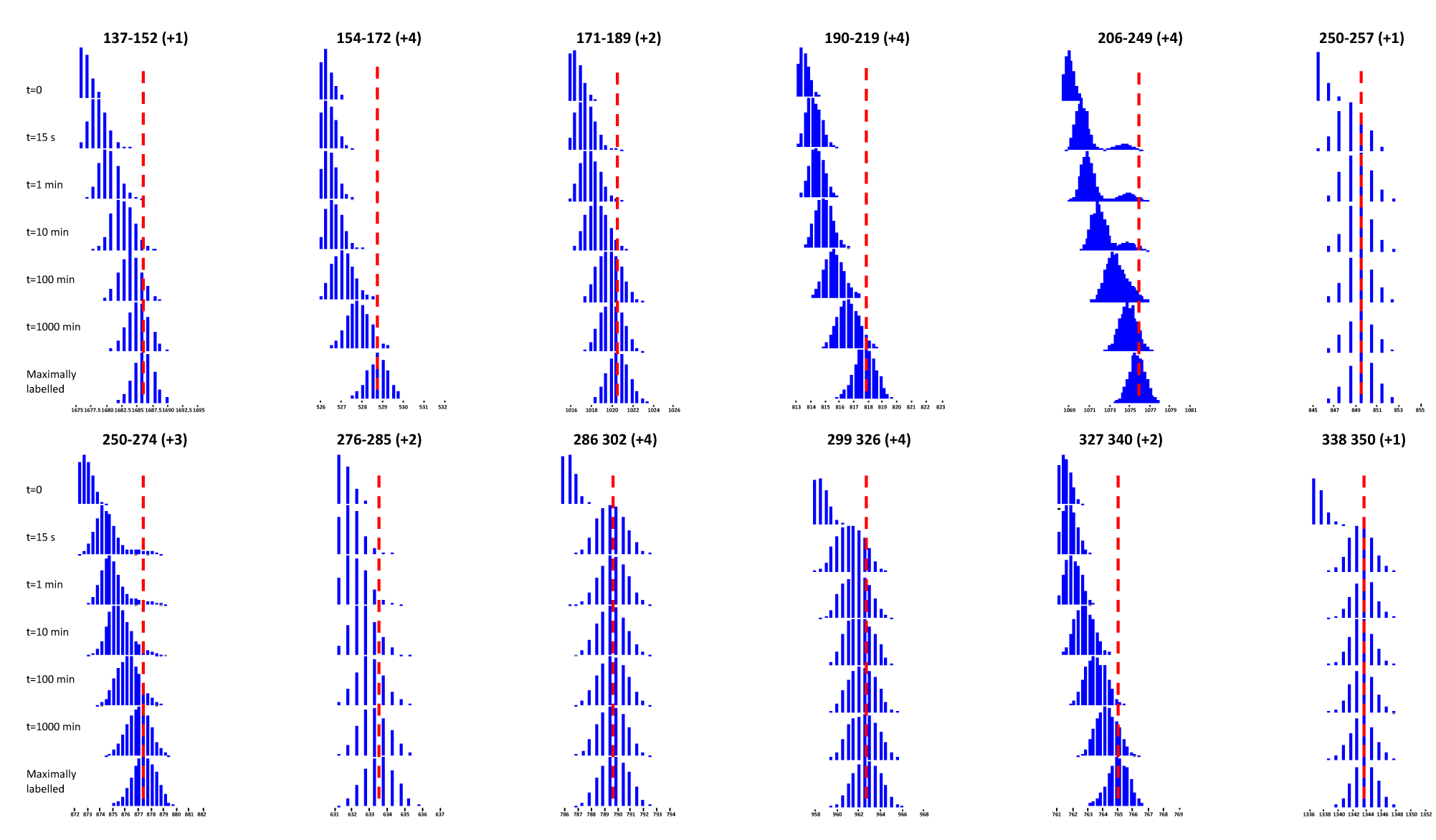
Bimodal HDX behavior of rNadA (from aa 137 to 350). Illustrative isotopic distributions of peptides spanning sequence 137-350. The dashed red line indicates the centroid mass of the maximally labelled state.

**Fig. S8.**
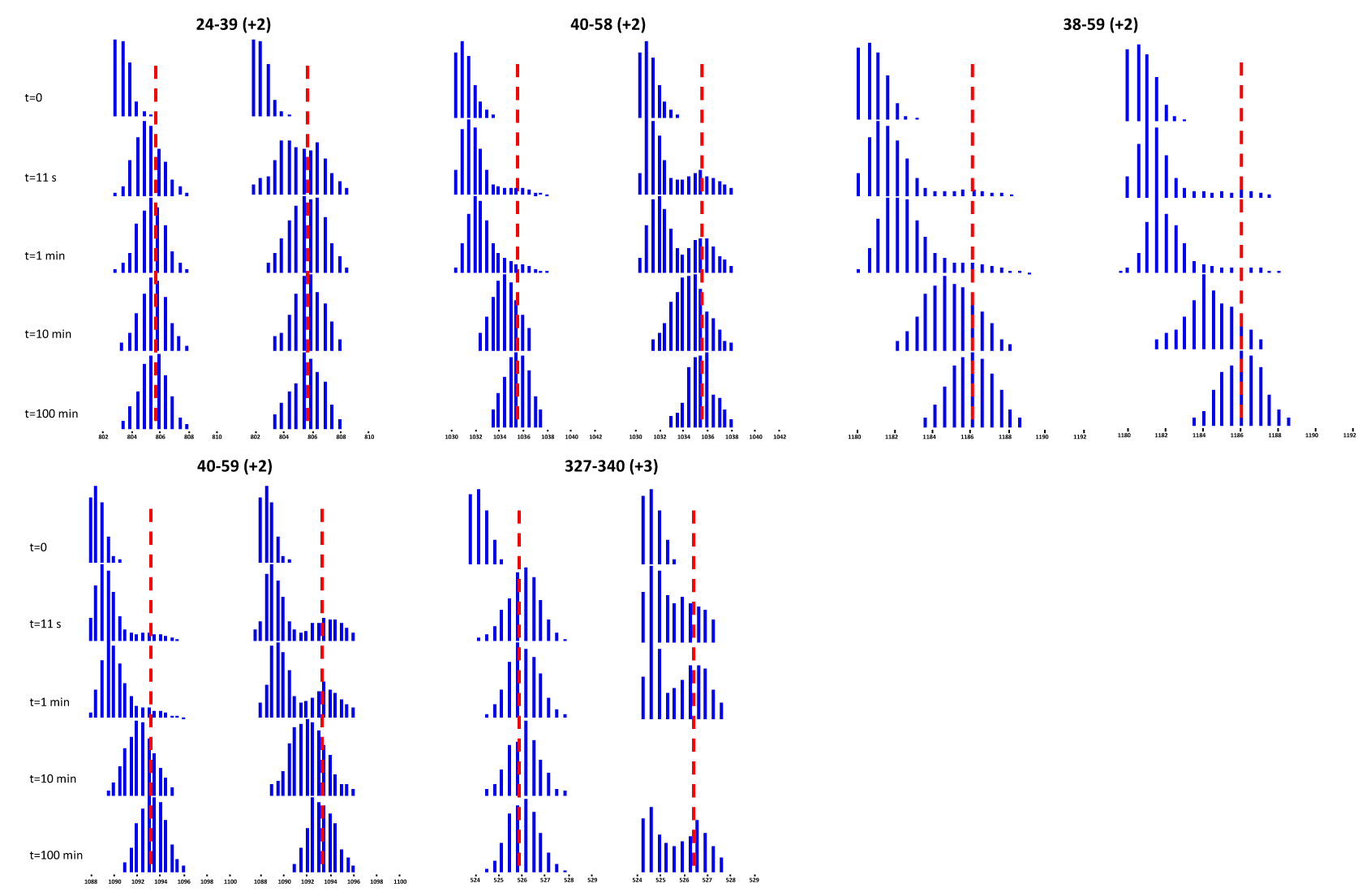
Bimodal HDX behavior of rNadA compared to OMV-embedded NadA. The dashed red line indicates the centroid mass of the isotopic envelope at 100 min. Panels on the left show the spectra of rNadA; panels on the right show the spectra of OMV-embedded NadA.

**Fig. S9.**
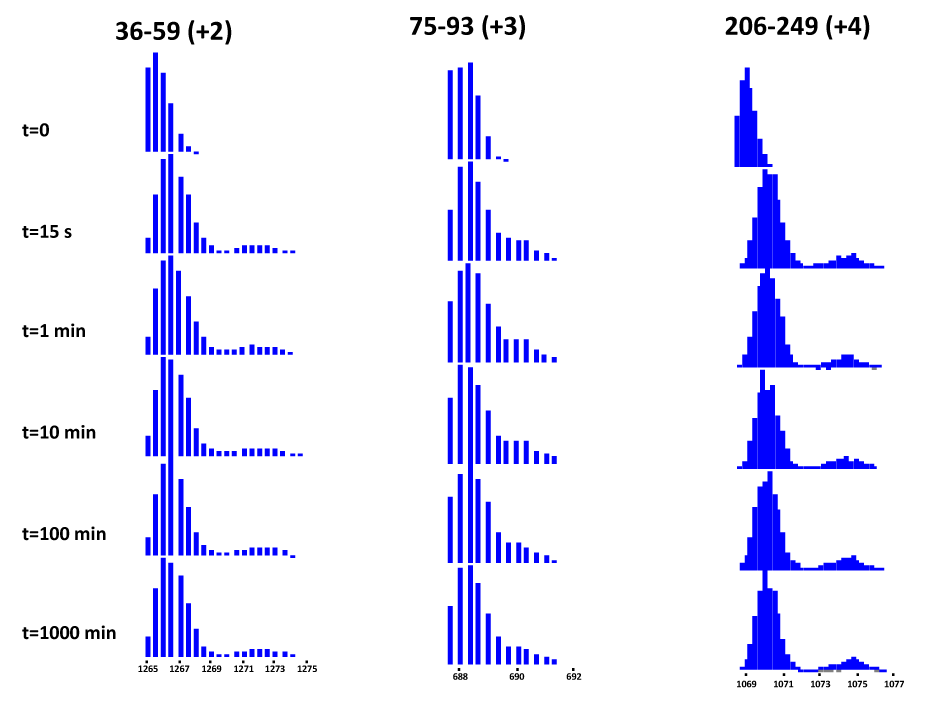
Pulse labelling HDX experiment of rNadA. Illustrative isotopic distributions showing unvaried centroid masses of both low- and high-mass envelopes. The peptides selected span the whole protein sequence displaying bimodality.

**Fig. S10.**
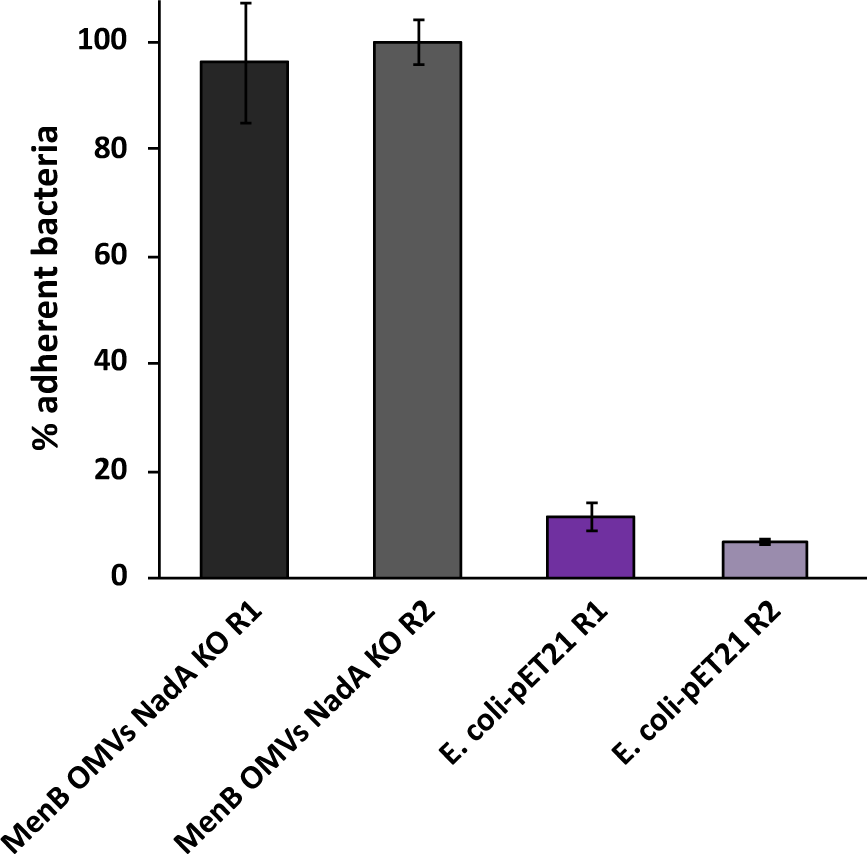
Inhibition of NadA-mediated adhesion of the sera raised against MenB OMVs NadA KO (1:20 dilution) in duplicates (R1 and R2) and adhesion of E. coli cells not expressing NadA (pET21) in duplicates. Sera raised against MenB OMVs NadA KO did not interfere with the adhesion mediated by *E. coli*-NadA^+^ cells, resulting in 100% adhesion relative to the adhesion in absence of sera, whereas *E. coli*-pET21 resulted in minimal adhesion.

## HDX summary tables

|  |  |  |
| --- | --- | --- |
| HDX reaction details | phosphate buffered saline (PBS), pH<sub>read</sub> 7,4 |  |
| HDX time course (min) | 11sec, 1min, 10min, 100min (RT) |  |
| HDX control samples | Maximally labelled control not performed |  |
| Back-exchange (mean / IQR) | - |  |
| # of Peptides | 36 |  |
| Sequence coverage | 72,8% |  |
| Average peptide length / Redundancy | 16,75 / 2,60 |  |
| Data Set | NadA recombinant | NadA OMV-embedded |
| Replicates (biological or technical) | 3 (technical) x time point | - |
| Repeatability | 0,0660 (average STD) | - |
| Significant differences in HDX (delta HDX ) | ≥0.62 D (99% CI) and ≥5 x STD of the time point |  |
| HDX reaction details | phosphate buffered saline (PBS), pH<sub>read</sub> 7,4 (RT) |  |
| HDX time course (min) | 15sec, 30sec, 1min (on ice) and 15sec, 1min, 10min, 30min, 100min, 1000min (at RT) |  |
| HDX control samples | Maximally-labeled control |  |
| Back-exchange (mean / IQR) | 29,5% / 14,1% |  |
| # of Peptides | 82 |  |
| Sequence coverage | 100% |  |
| Average peptide length / Redundancy | 16,21 / 4,06 |  |
| Data Set | NadA alone | NadA + Hsp90 |
| Replicates (biological or technical) | 3 (technical) | 3 (technical) |
| Repeatability | 0,0416 (average STD) | 0,1002 (average STD) |
| Significant differences in HDX (delta HDX > X D) | 0.52 D (98% CI) |  |

