## Supplementary figures and images for "Structural dynamics and immunogenicity of the recombinant and outer membrane vesicle-embedded Meningococcal antigen NadA"

### Supplementary dataset 1

#### Supplementary dataset 1

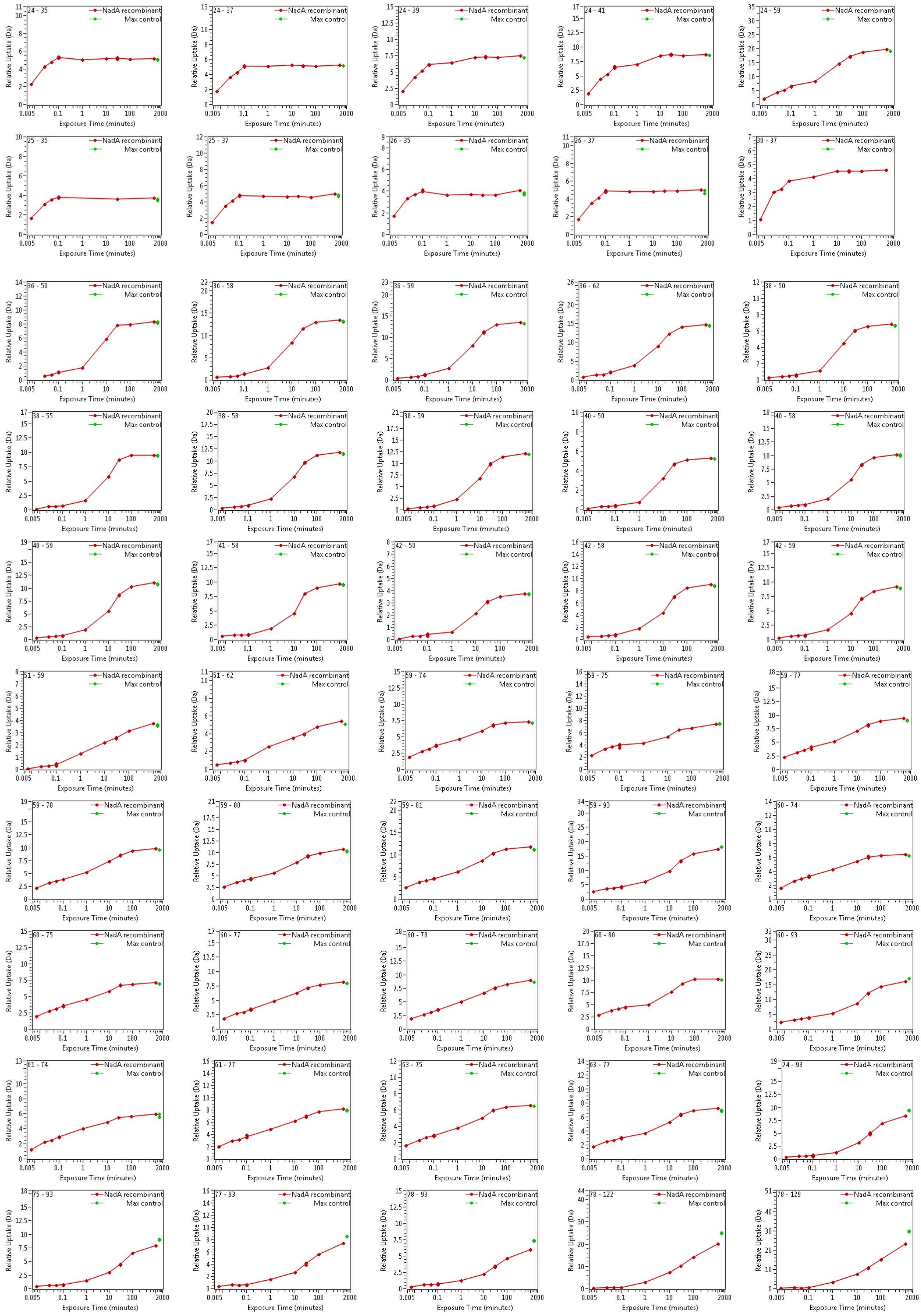

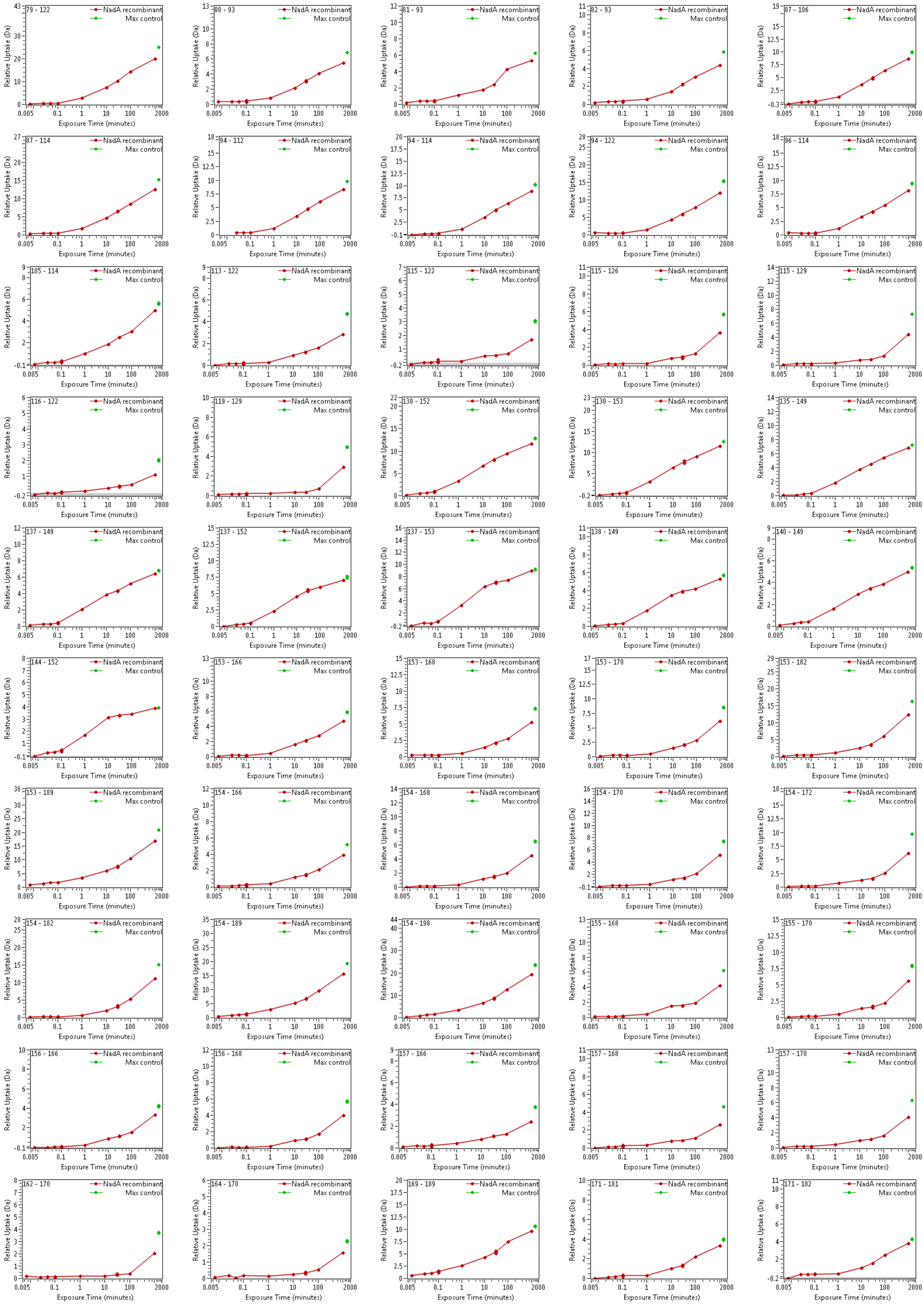

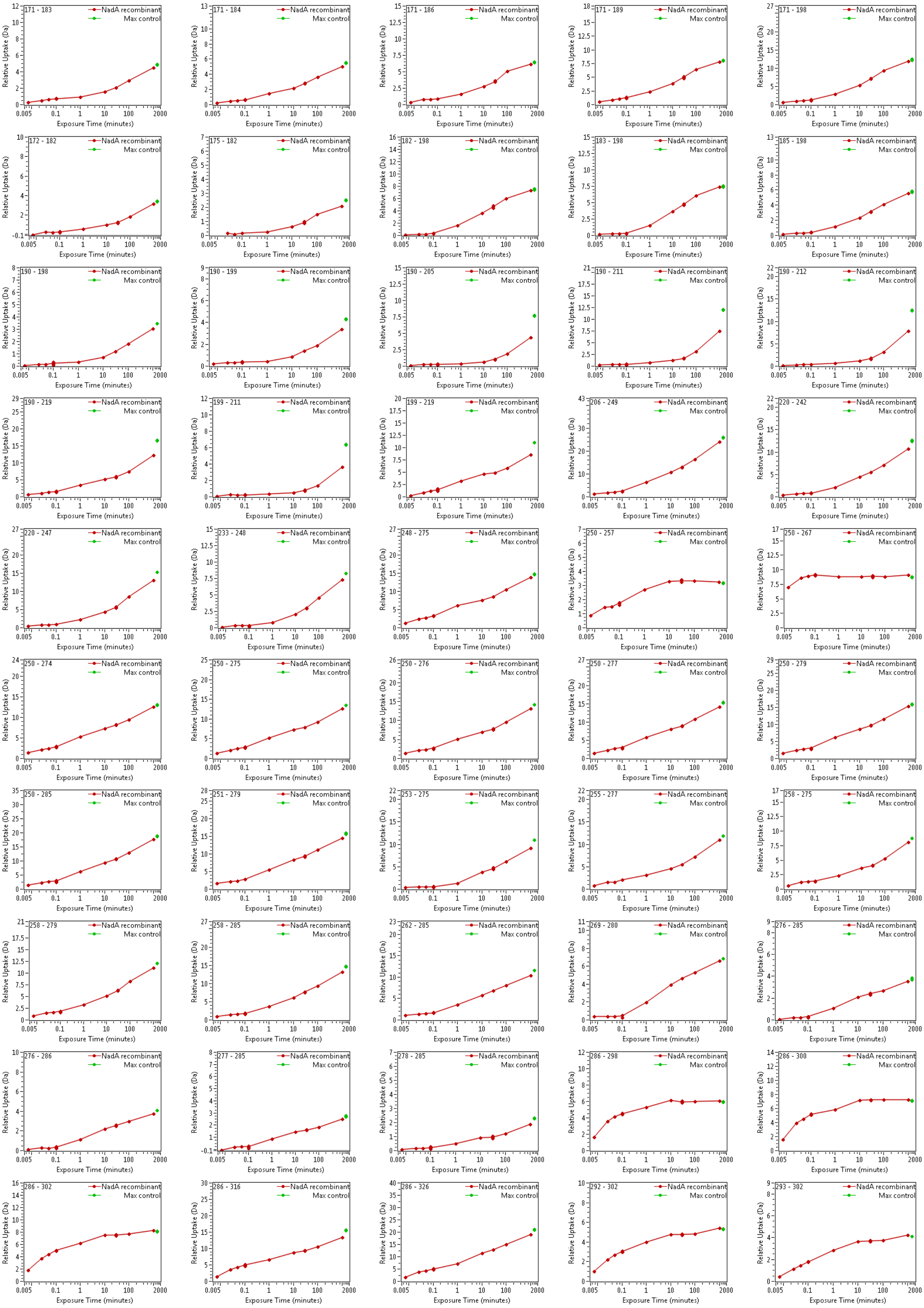

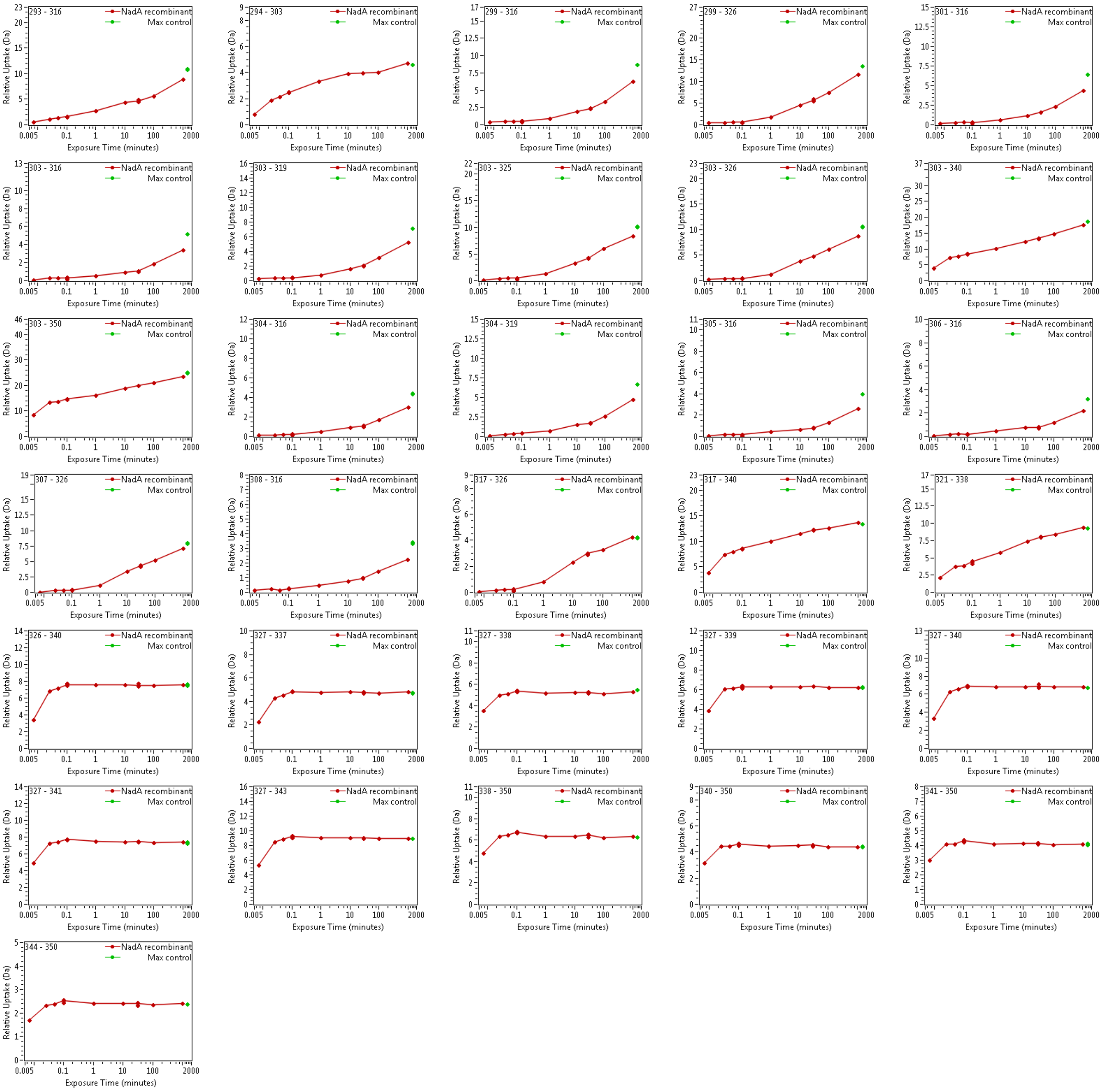

### Supplementary dataset 3

#### Supplementary dataset 3

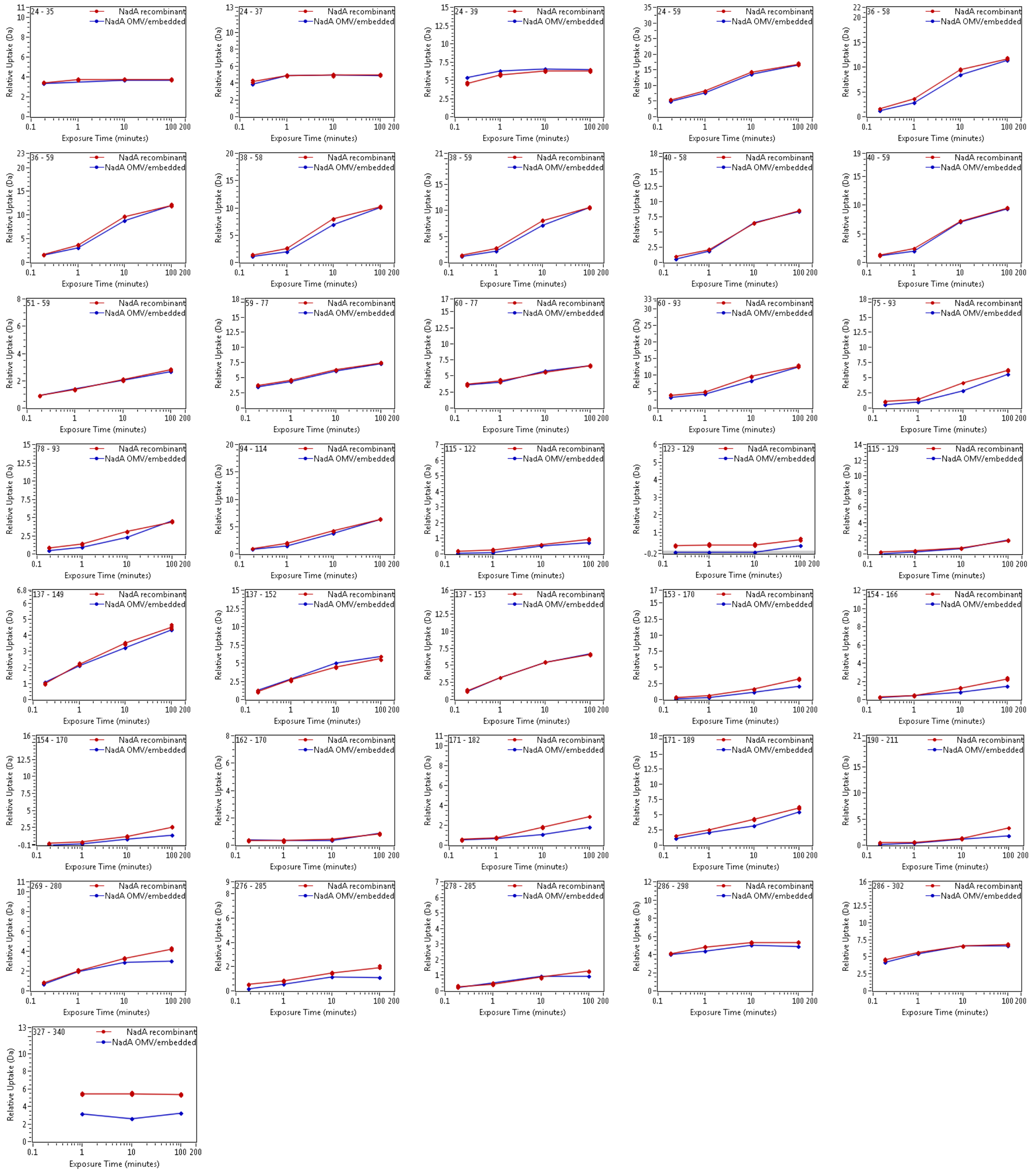

### Supplementary dataset 4

#### Supplementary dataset 4

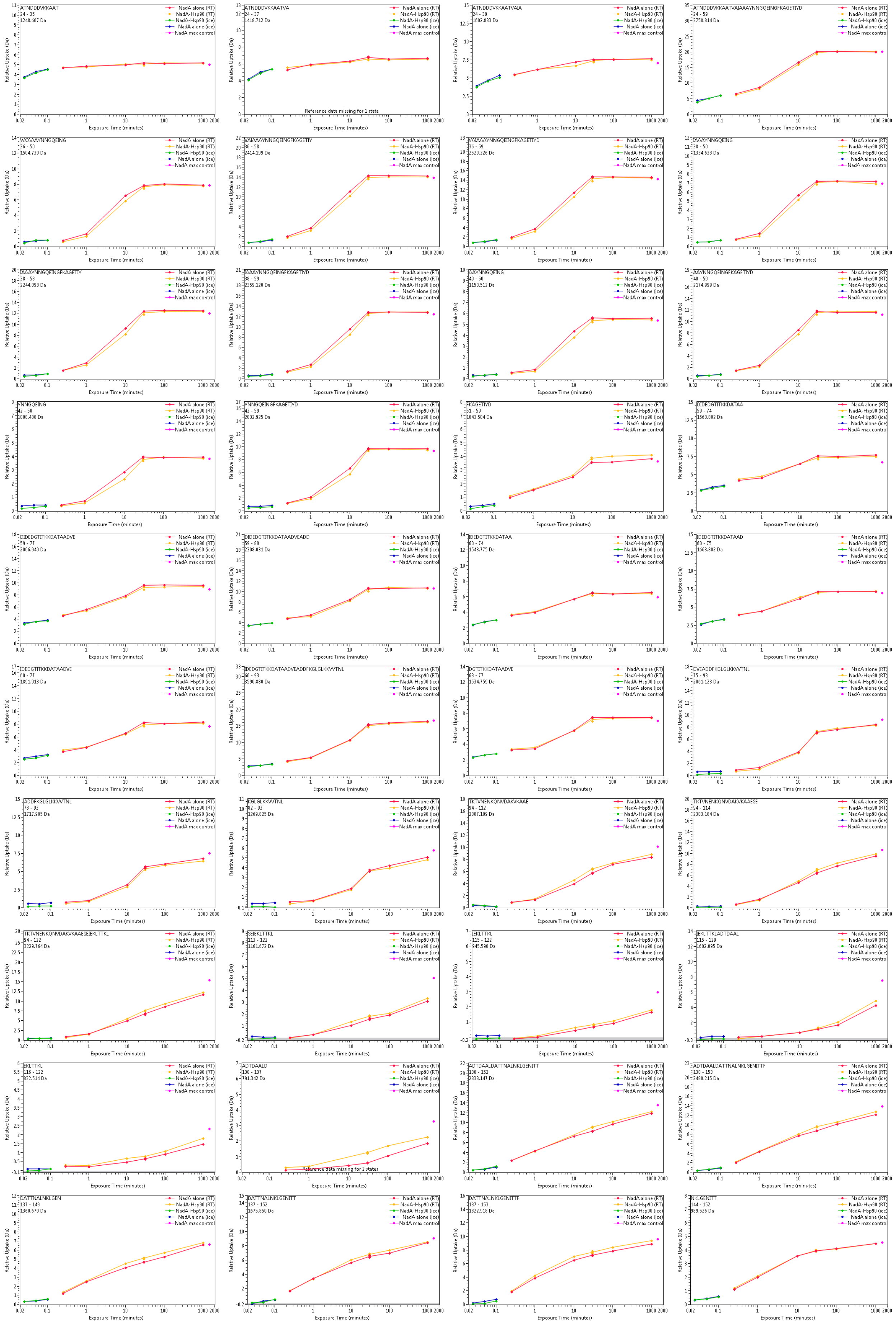

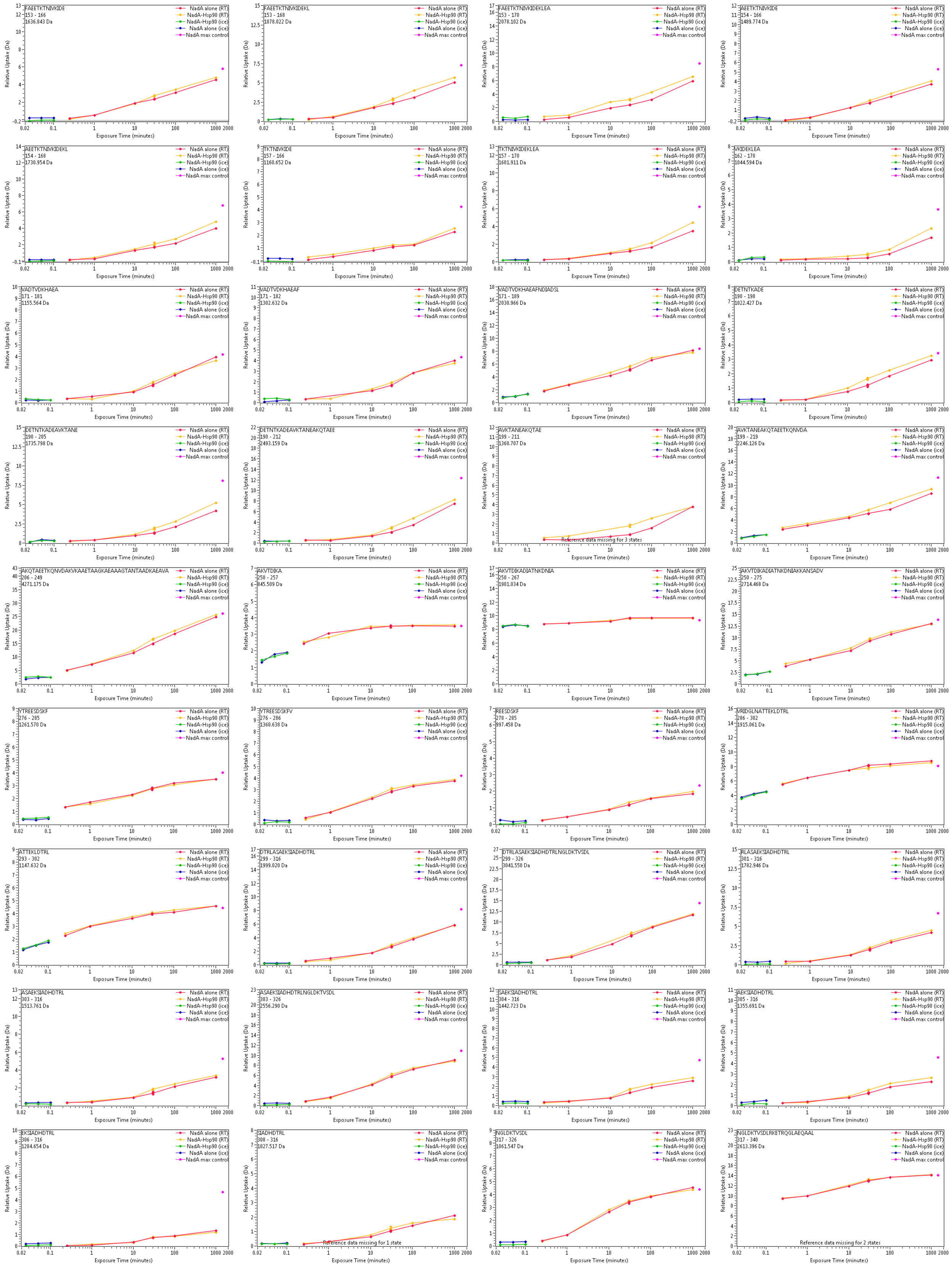

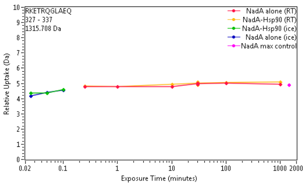

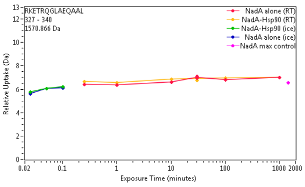

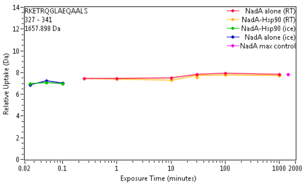

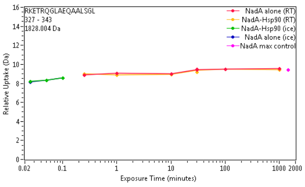

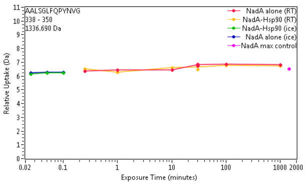

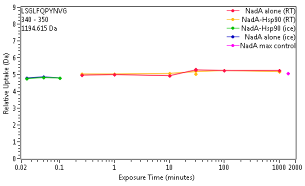
