## Supplementary dataset 2 for "Structural dynamics and immunogenicity of the recombinant and outer membrane vesicle-embedded Meningococcal antigen NadA"

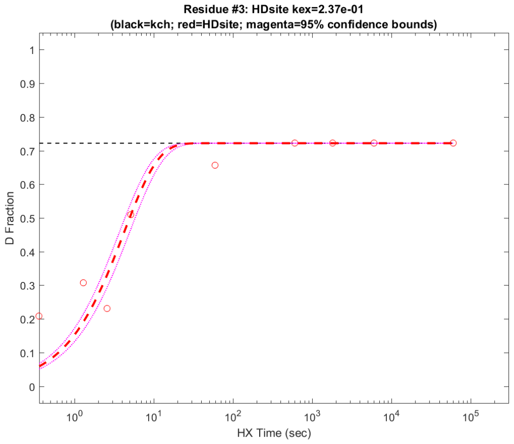


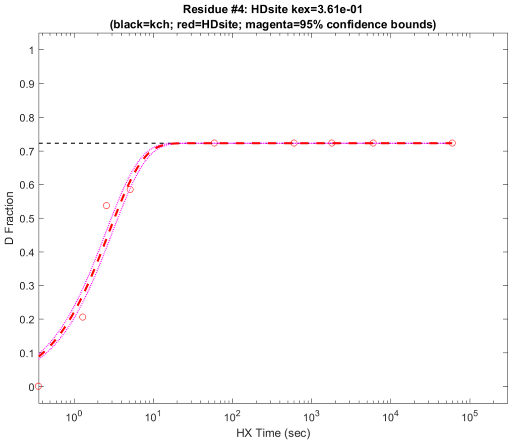


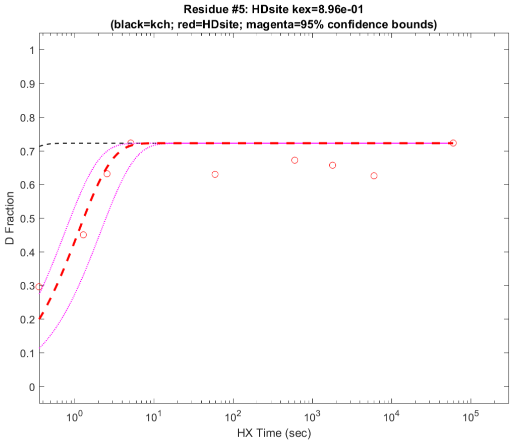


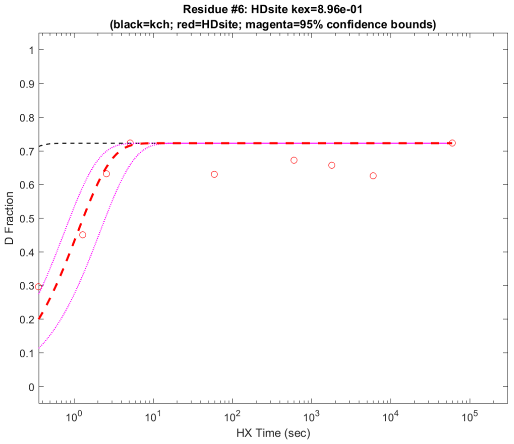


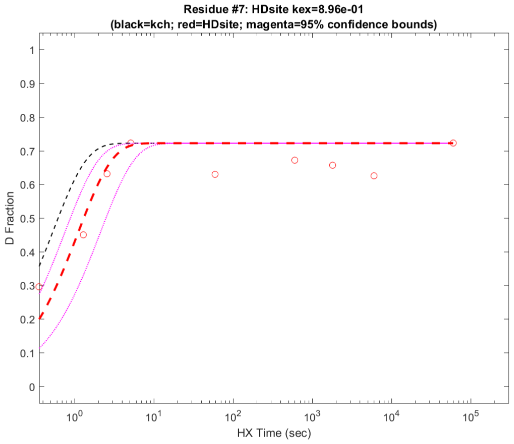


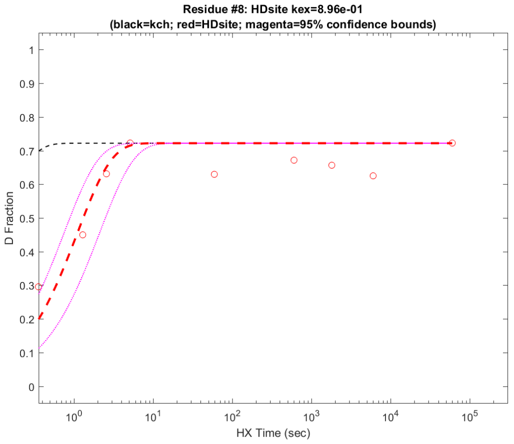


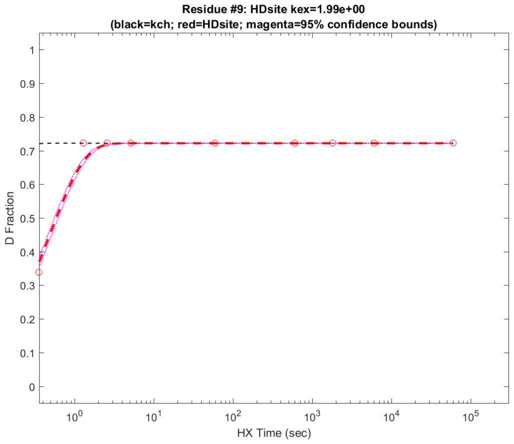


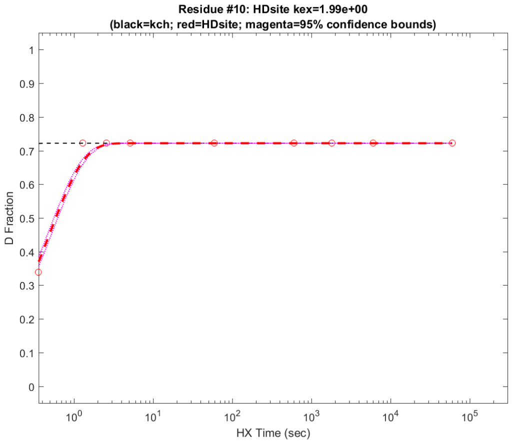

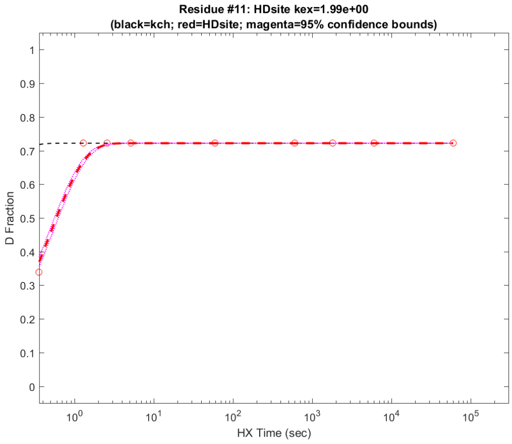
Residue #11: HDsite kex=1.99e+00
(black=kch; red=HDsite; magenta=95% confidence bounds)
D Fraction
HX Time (sec)

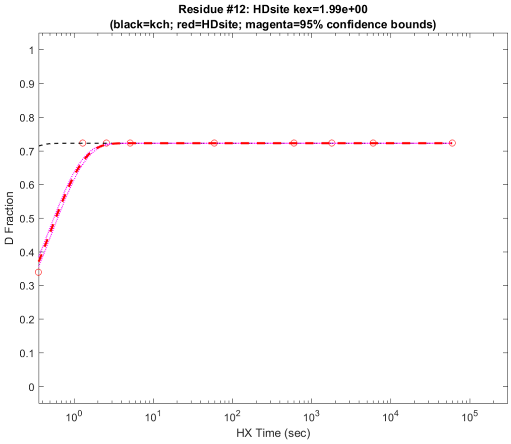
Residue #12: HDsite kex=1.99e+00
(black=kch; red=HDsite; magenta=95% confidence bounds)
D Fraction
HX Time (sec)

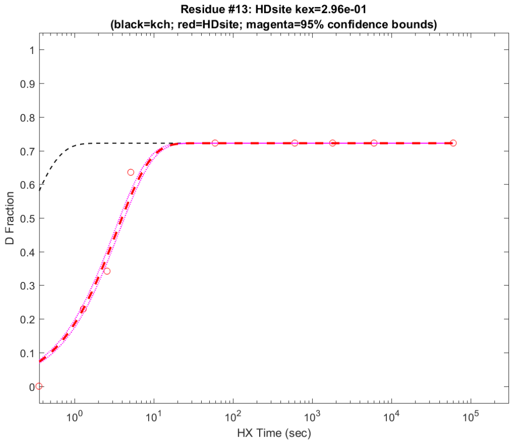
Residue #13: HDsite kex=2.96e-01
(black=kch; red=HDsite; magenta=95% confidence bounds)
D Fraction
HX Time (sec)

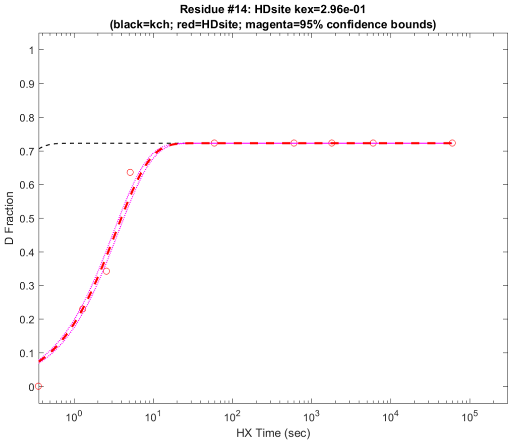
Residue #14: HDsite kex=2.96e-01
(black=kch; red=HDsite; magenta=95% confidence bounds)
D Fraction
HX Time (sec)

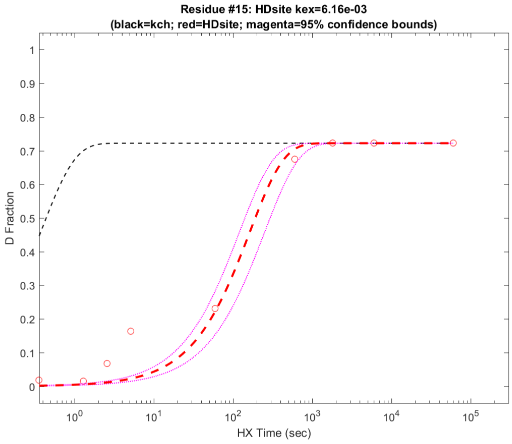
Residue #15: HDsite kex=6.16e-03
(black=kch; red=HDsite; magenta=95% confidence bounds)
D Fraction
HX Time (sec)

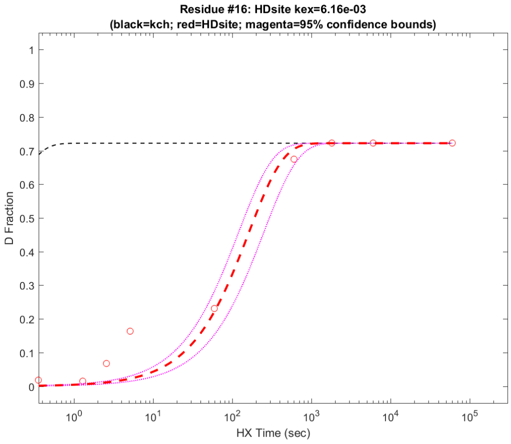
Residue #16: HDsite kex=6.16e-03
(black=kch; red=HDsite; magenta=95% confidence bounds)
D Fraction
HX Time (sec)

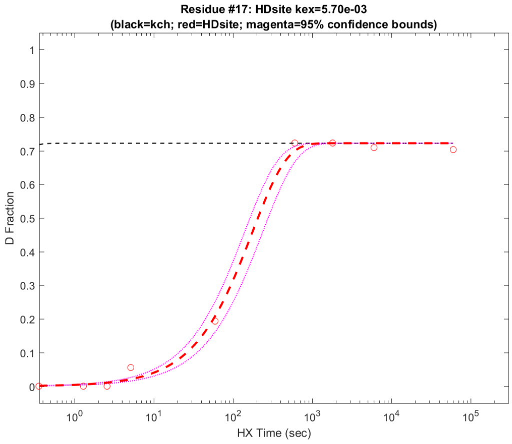
Residue #17: HDsite kex=5.70e-03
(black=kch; red=HDsite; magenta=95% confidence bounds)
D Fraction
HX Time (sec)

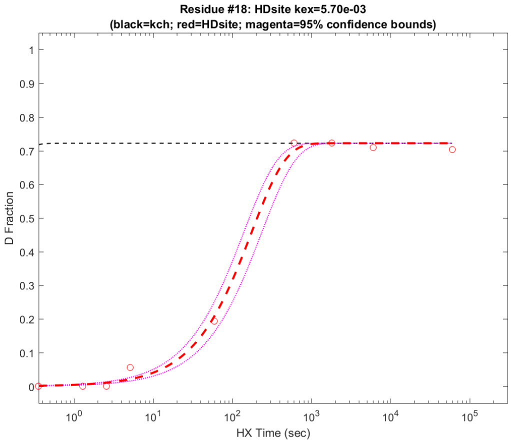
Residue #18: HDsite kex=5.70e-03
(black=kch; red=HDsite; magenta=95% confidence bounds)
D Fraction
HX Time (sec)

**Residue #19: HDsite kex=1.02e-03**
**(black=kch; red=HDsite; magenta=95% confidence bounds)**

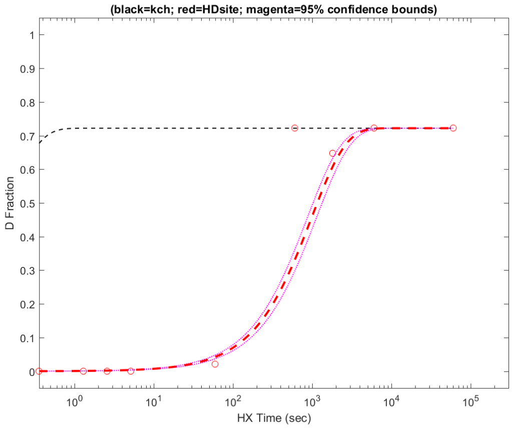


**Residue #20: HDsite kex=2.34e-03**
**(black=kch; red=HDsite; magenta=95% confidence bounds)**

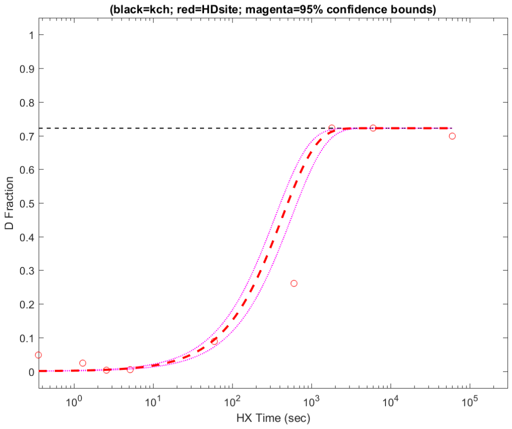


**Residue #21: HDsite kex=1.24e-03**
**(black=kch; red=HDsite; magenta=95% confidence bounds)**

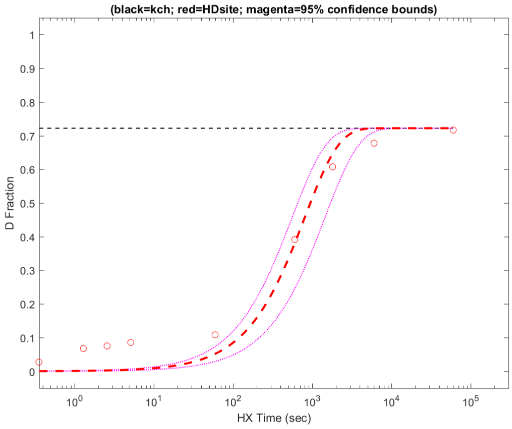


**Residue #22: HDsite kex=1.24e-03**
**(black=kch; red=HDsite; magenta=95% confidence bounds)**

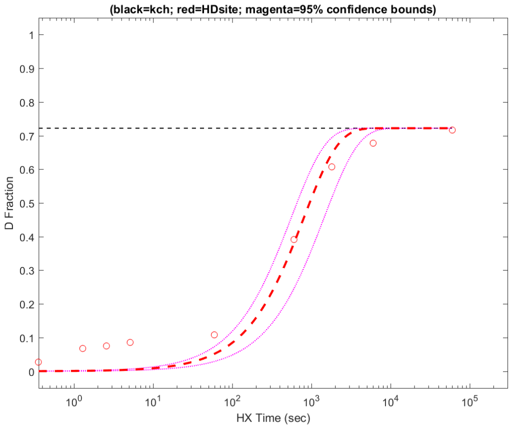


**Residue #23: HDsite kex=1.24e-03**
**(black=kch; red=HDsite; magenta=95% confidence bounds)**

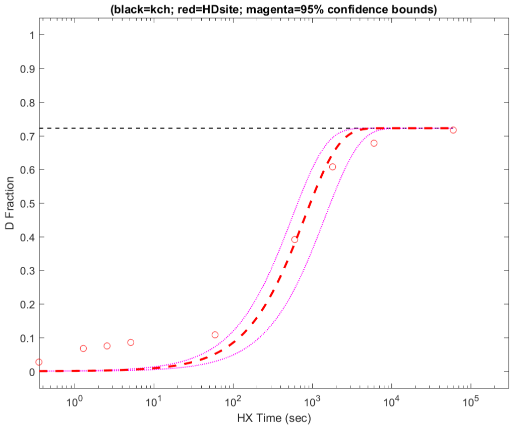


**Residue #24: HDsite kex=1.24e-03**
**(black=kch; red=HDsite; magenta=95% confidence bounds)**

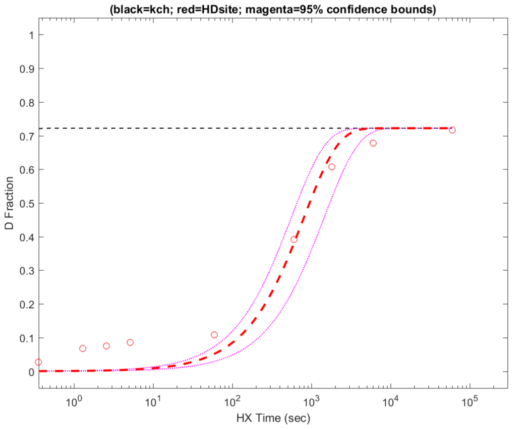


**Residue #25: HDsite kex=1.24e-03**
**(black=kch; red=HDsite; magenta=95% confidence bounds)**

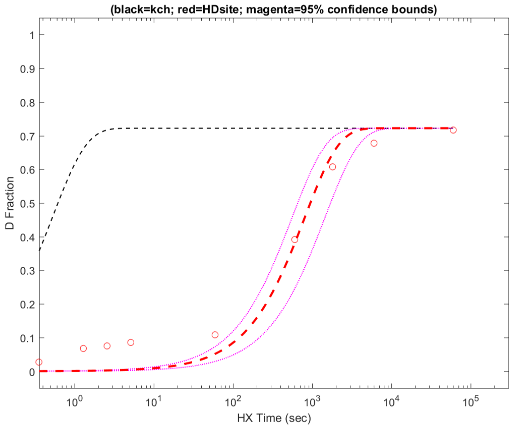


**Residue #26: HDsite kex=1.24e-03**
**(black=kch; red=HDsite; magenta=95% confidence bounds)**

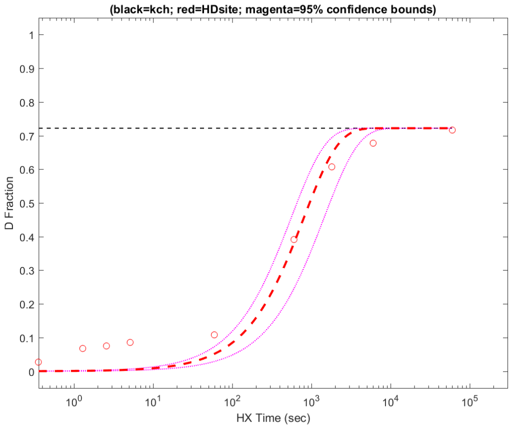

**Residue #27: HDsite kex=1.24e-03**
**(black=kch; red=HDsite; magenta=95% confidence bounds)**

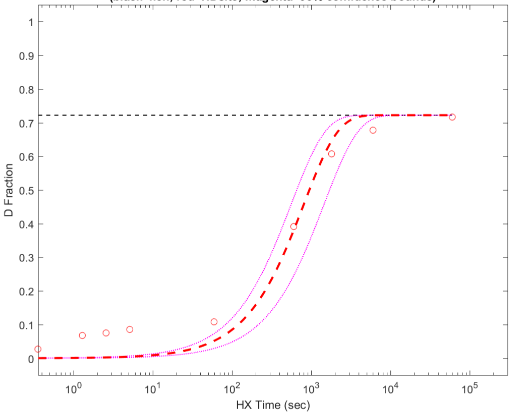


**Residue #28: HDsite kex=7.46e-04**
**(black=kch; red=HDsite; magenta=95% confidence bounds)**

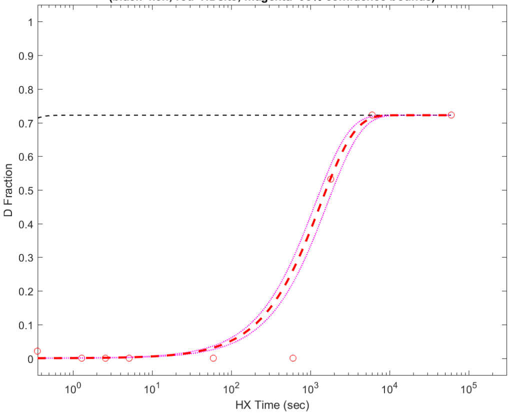


**Residue #29: HDsite kex=7.46e-04**
**(black=kch; red=HDsite; magenta=95% confidence bounds)**

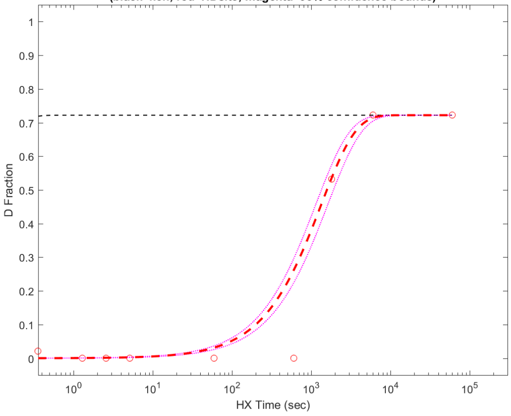


**Residue #30: HDsite kex=5.90e-03**
**(black=kch; red=HDsite; magenta=95% confidence bounds)**

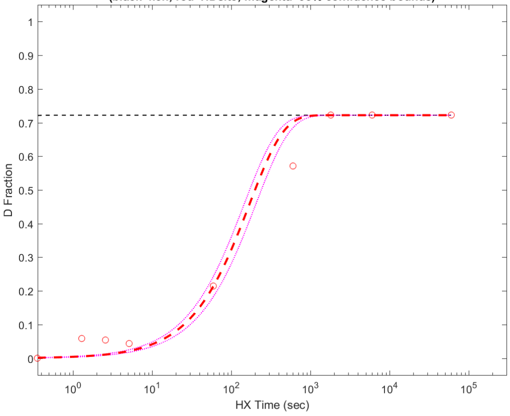


**Residue #31: HDsite kex=5.90e-03**
**(black=kch; red=HDsite; magenta=95% confidence bounds)**

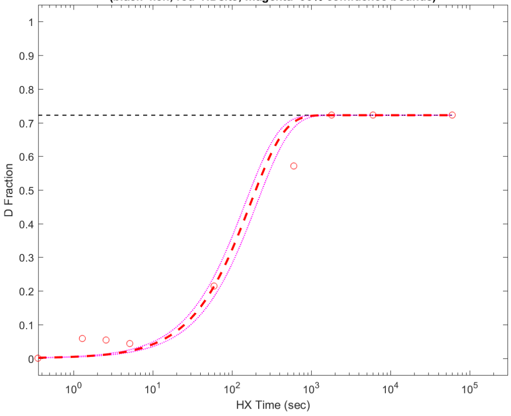


**Residue #32: HDsite kex=5.90e-03**
**(black=kch; red=HDsite; magenta=95% confidence bounds)**

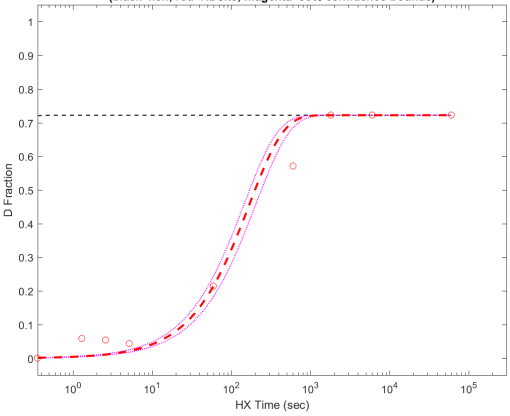


**Residue #33: HDsite kex=5.34e-04**
**(black=kch; red=HDsite; magenta=95% confidence bounds)**

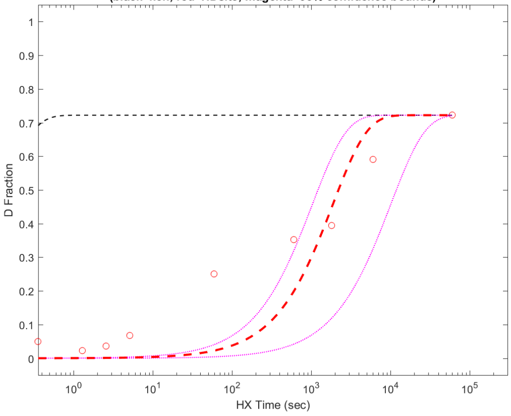


**Residue #34: HDsite kex=5.34e-04**
**(black=kch; red=HDsite; magenta=95% confidence bounds)**

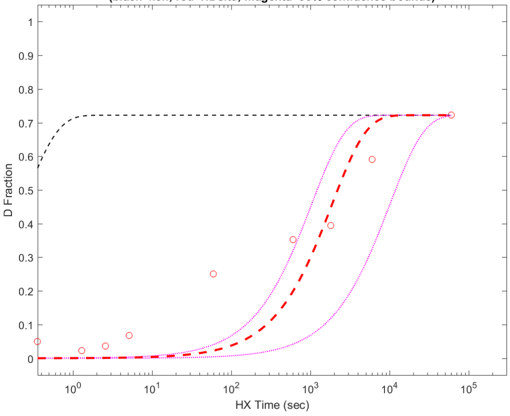

**Residue #35: HDsite kex=5.34e-04**
**(black=kch; red=HDsite; magenta=95% confidence bounds)**

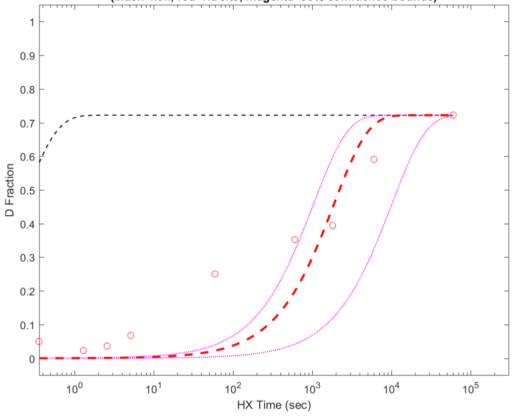


**Residue #36: HDsite kex=1.68e-04**
**(black=kch; red=HDsite; magenta=95% confidence bounds)**

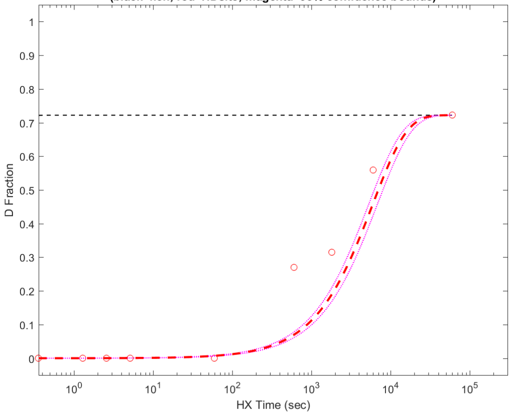


**Residue #37: HDsite kex=2.34e-01**
**(black=kch; red=HDsite; magenta=95% confidence bounds)**

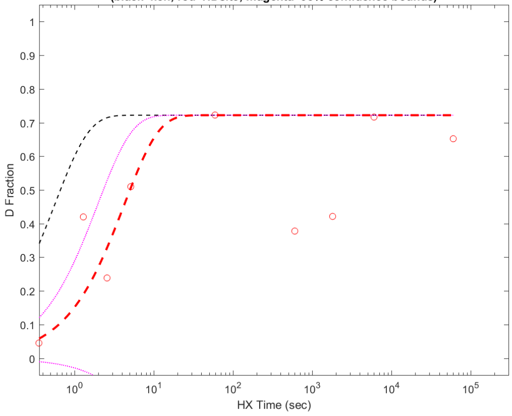


**Residue #38: HDsite kex=1.14e-03**
**(black=kch; red=HDsite; magenta=95% confidence bounds)**

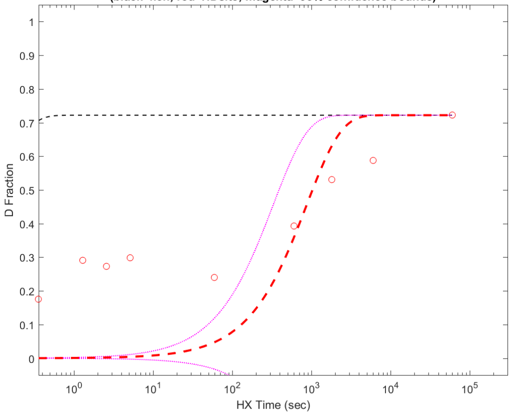


**Residue #39: HDsite kex=4.12e-02**
**(black=kch; red=HDsite; magenta=95% confidence bounds)**

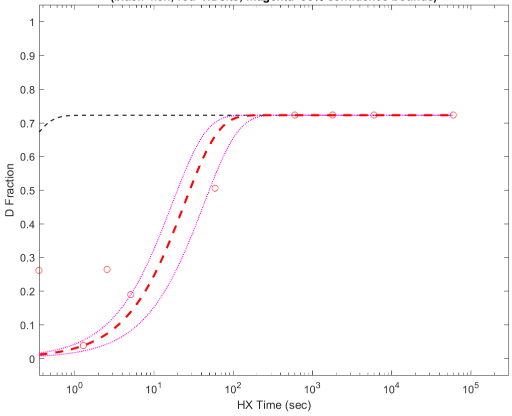


**Residue #40: HDsite kex=1.27e+00**
**(black=kch; red=HDsite; magenta=95% confidence bounds)**

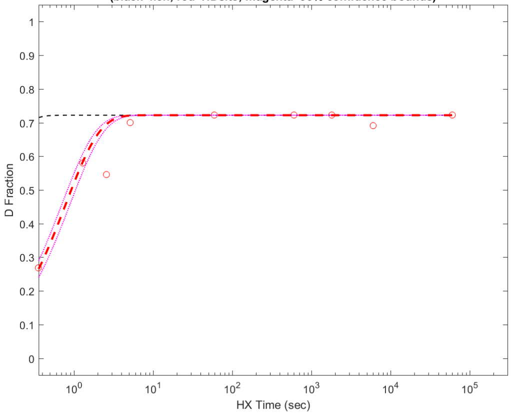


**Residue #41: HDsite kex=1.27e+00**
**(black=kch; red=HDsite; magenta=95% confidence bounds)**

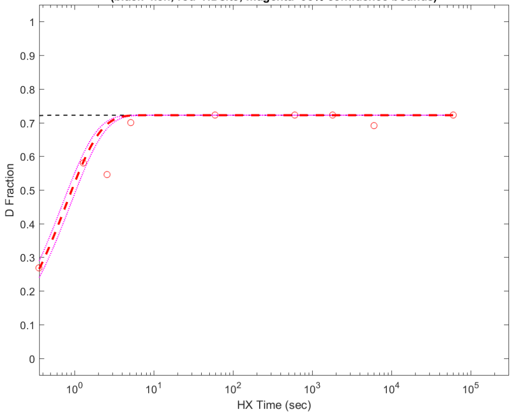


**Residue #42: HDsite kex=1.78e-01**
**(black=kch; red=HDsite; magenta=95% confidence bounds)**

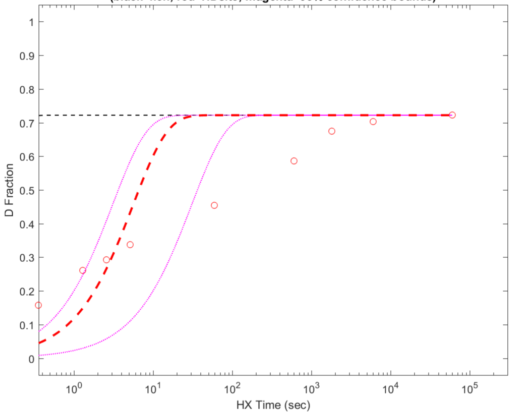

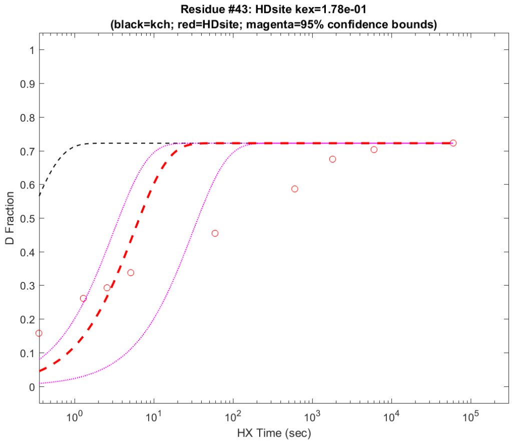
Residue #43: HDsite kex=1.78e-01
(black=kch; red=HDsite; magenta=95% confidence bounds)
D Fraction
HX Time (sec)

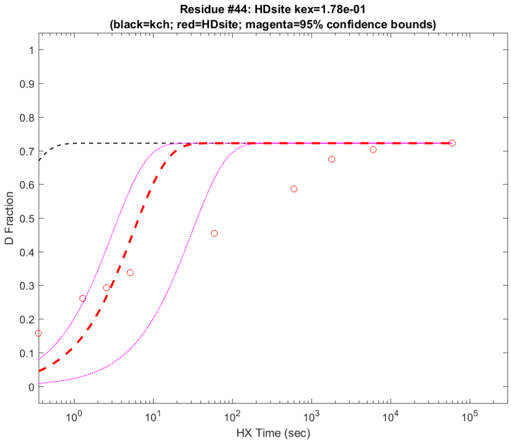
Residue #44: HDsite kex=1.78e-01
(black=kch; red=HDsite; magenta=95% confidence bounds)
D Fraction
HX Time (sec)

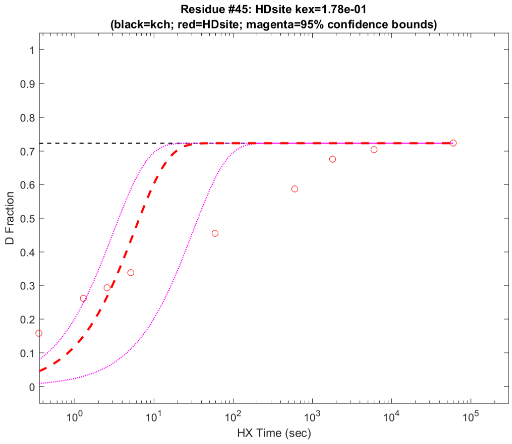
Residue #45: HDsite kex=1.78e-01
(black=kch; red=HDsite; magenta=95% confidence bounds)
D Fraction
HX Time (sec)

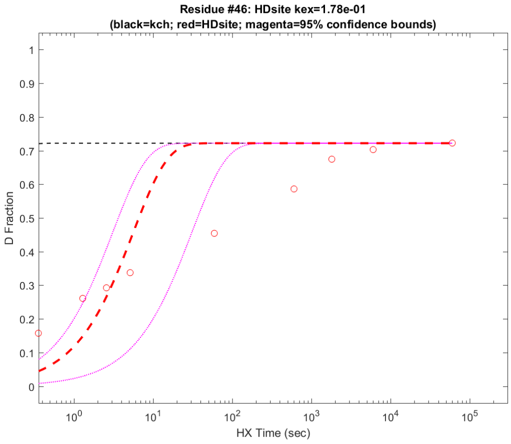
Residue #46: HDsite kex=1.78e-01
(black=kch; red=HDsite; magenta=95% confidence bounds)
D Fraction
HX Time (sec)

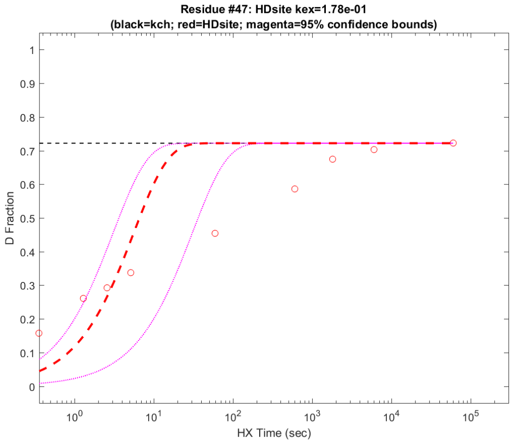
Residue #47: HDsite kex=1.78e-01
(black=kch; red=HDsite; magenta=95% confidence bounds)
D Fraction
HX Time (sec)

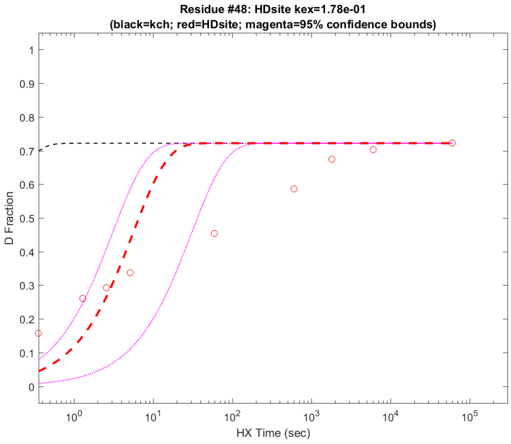
Residue #48: HDsite kex=1.78e-01
(black=kch; red=HDsite; magenta=95% confidence bounds)
D Fraction
HX Time (sec)

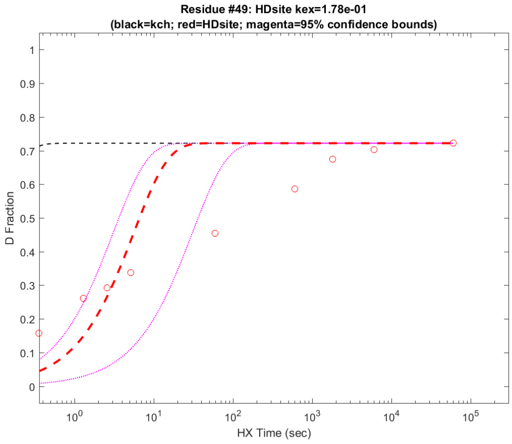
Residue #49: HDsite kex=1.78e-01
(black=kch; red=HDsite; magenta=95% confidence bounds)
D Fraction
HX Time (sec)

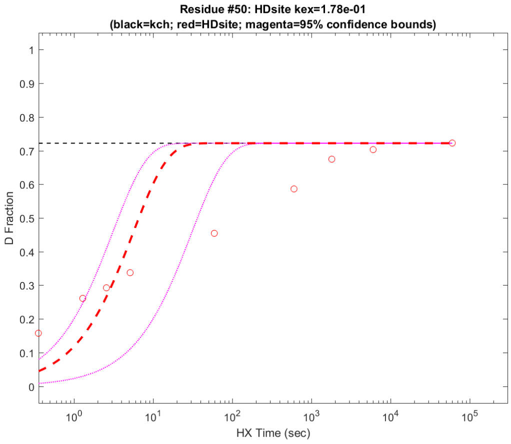
Residue #50: HDsite kex=1.78e-01
(black=kch; red=HDsite; magenta=95% confidence bounds)
D Fraction
HX Time (sec)

**Residue #51: HDsite kex=1.78e-01**
**(black=kch; red=HDsite; magenta=95% confidence bounds)**

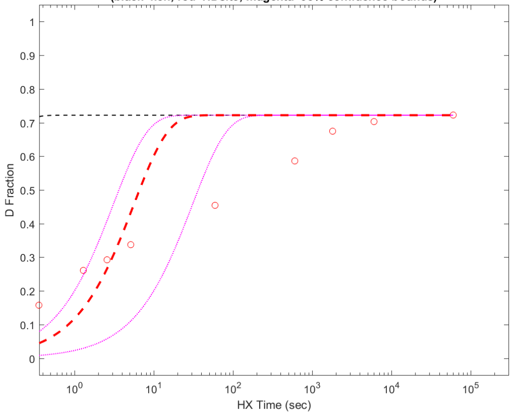


**Residue #52: HDsite kex=1.85e-04**
**(black=kch; red=HDsite; magenta=95% confidence bounds)**

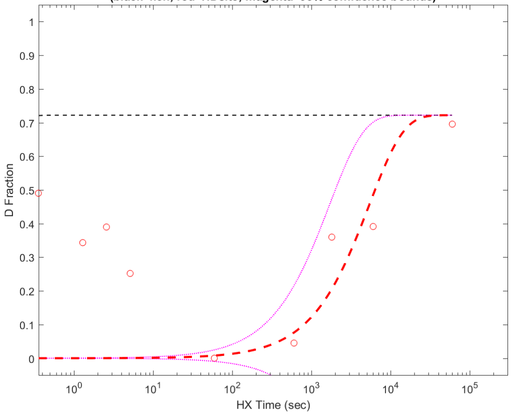


**Residue #53: HDsite kex=1.02e-03**
**(black=kch; red=HDsite; magenta=95% confidence bounds)**

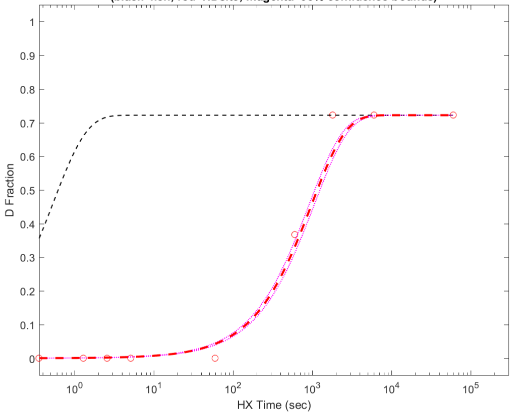


**Residue #54: HDsite kex=1.30e-02**
**(black=kch; red=HDsite; magenta=95% confidence bounds)**

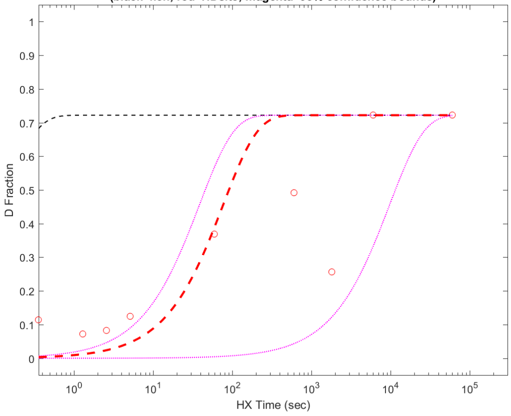


**Residue #55: HDsite kex=8.99e-04**
**(black=kch; red=HDsite; magenta=95% confidence bounds)**

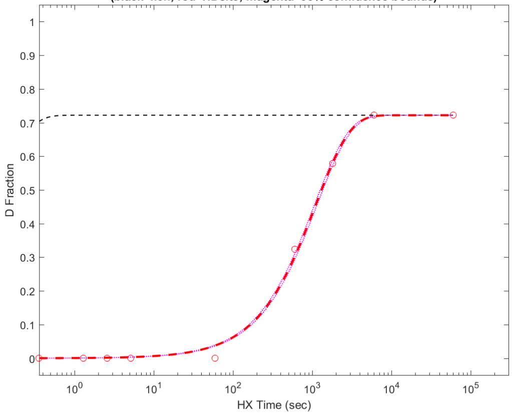


**Residue #56: HDsite kex=8.86e-04**
**(black=kch; red=HDsite; magenta=95% confidence bounds)**

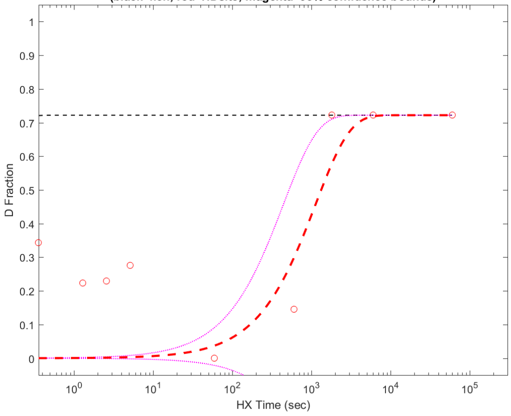


**Residue #57: HDsite kex=1.21e+00**
**(black=kch; red=HDsite; magenta=95% confidence bounds)**

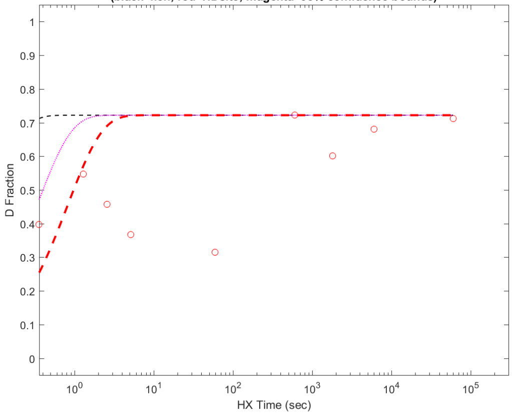


**Residue #58: HDsite kex=1.00e-03**
**(black=kch; red=HDsite; magenta=95% confidence bounds)**

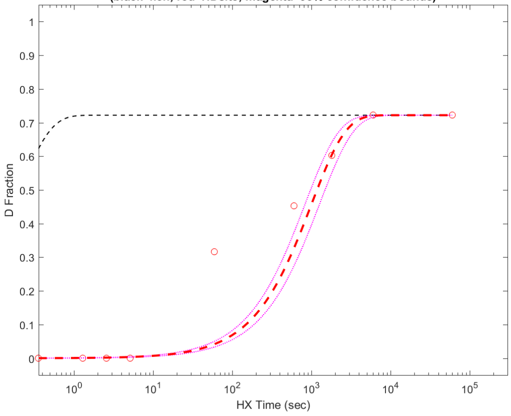

**Residue #59: HDsite kex=7.03e-05**
**(black=kch; red=HDsite; magenta=95% confidence bounds)**

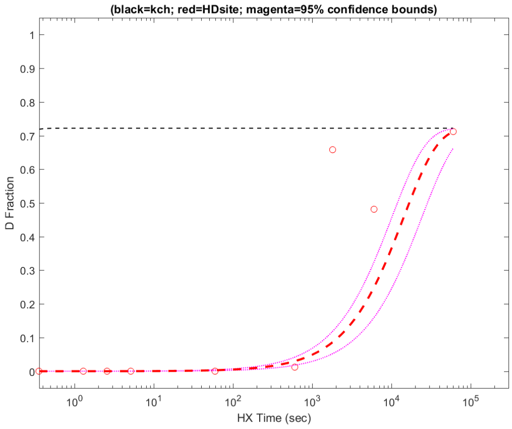


**Residue #60: HDsite kex=1.02e-03**
**(black=kch; red=HDsite; magenta=95% confidence bounds)**

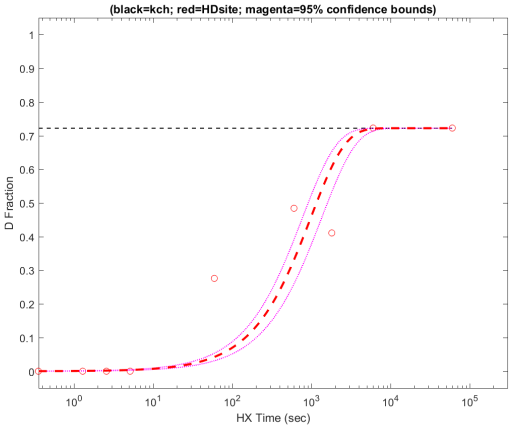


**Residue #61: HDsite kex=1.66e-04**
**(black=kch; red=HDsite; magenta=95% confidence bounds)**

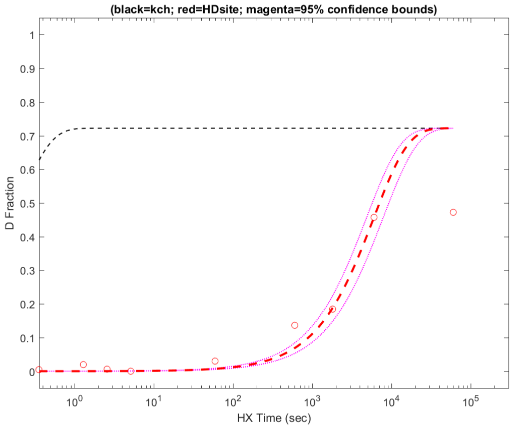


**Residue #62: HDsite kex=1.66e-04**
**(black=kch; red=HDsite; magenta=95% confidence bounds)**

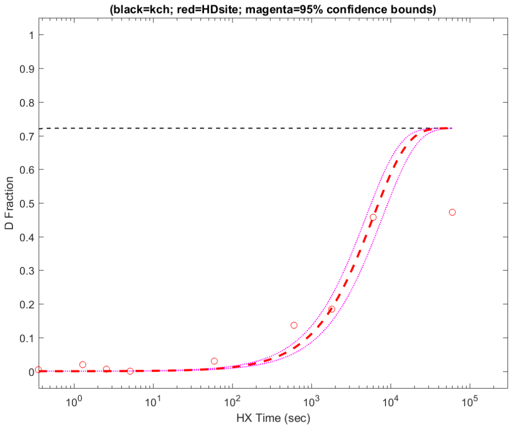


**Residue #63: HDsite kex=1.66e-04**
**(black=kch; red=HDsite; magenta=95% confidence bounds)**

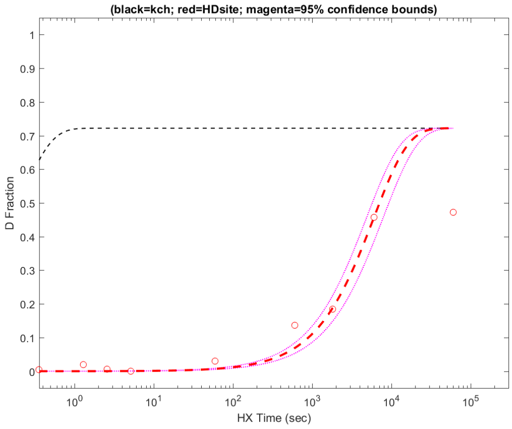


**Residue #64: HDsite kex=1.66e-04**
**(black=kch; red=HDsite; magenta=95% confidence bounds)**

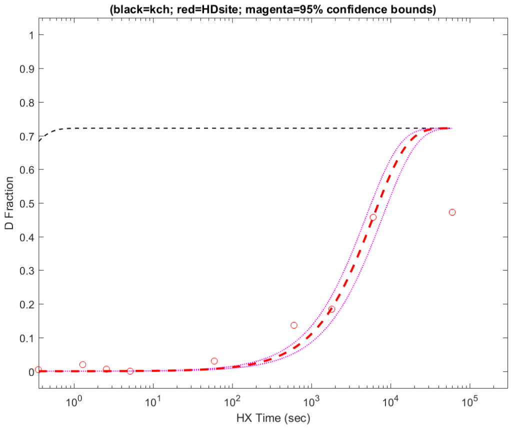


**Residue #65: HDsite kex=1.66e-04**
**(black=kch; red=HDsite; magenta=95% confidence bounds)**

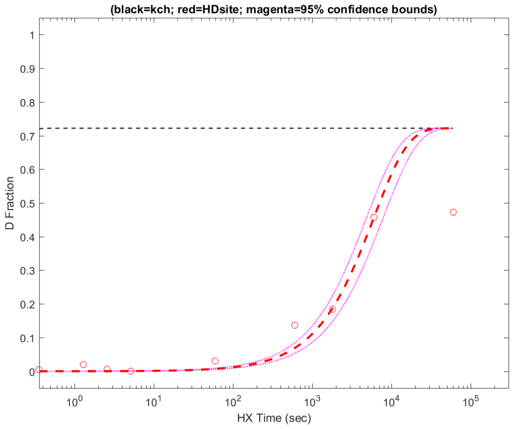


**Residue #66: HDsite kex=4.15e-04**
**(black=kch; red=HDsite; magenta=95% confidence bounds)**

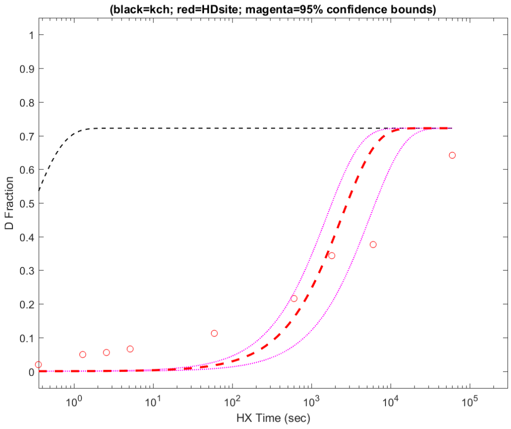

**Residue #67: HDsite kex=4.15e-04**
**(black=kch; red=HDsite; magenta=95% confidence bounds)**

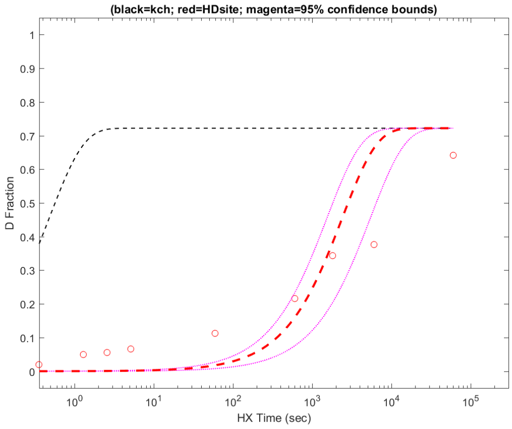


**Residue #68: HDsite kex=4.15e-04**
**(black=kch; red=HDsite; magenta=95% confidence bounds)**

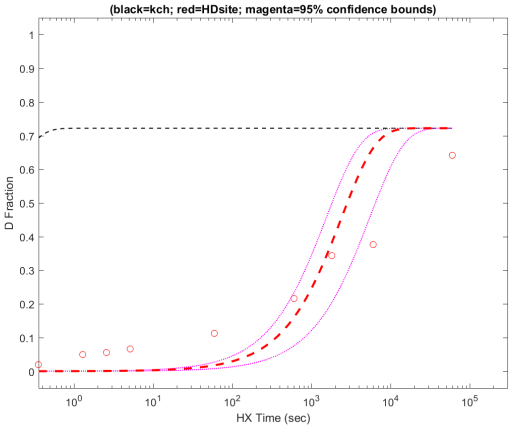


**Residue #69: HDsite kex=4.15e-04**
**(black=kch; red=HDsite; magenta=95% confidence bounds)**

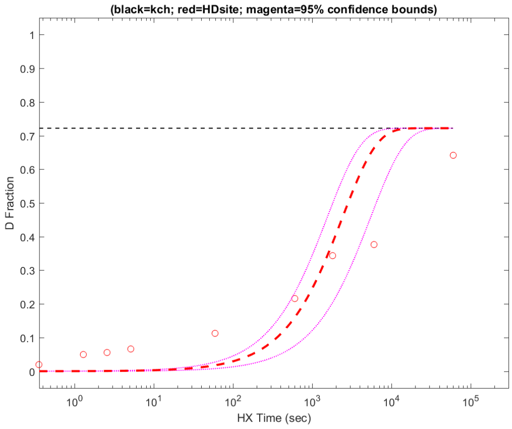


**Residue #70: HDsite kex=4.15e-04**
**(black=kch; red=HDsite; magenta=95% confidence bounds)**

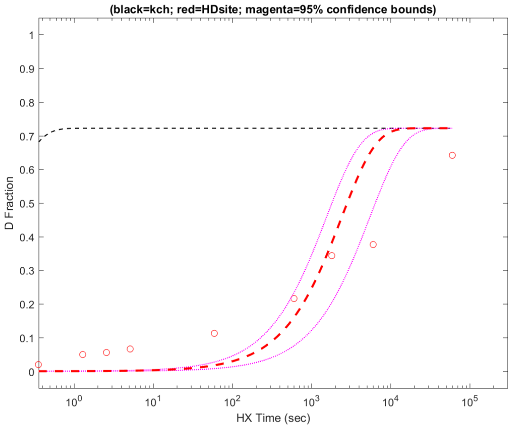


**Residue #71: HDsite kex=1.34e-04**
**(black=kch; red=HDsite; magenta=95% confidence bounds)**

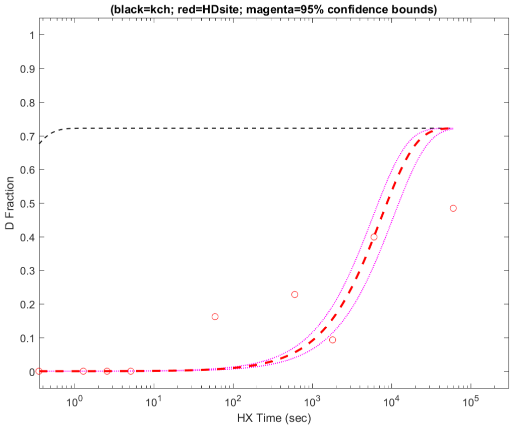


**Residue #72: HDsite kex=1.34e-04**
**(black=kch; red=HDsite; magenta=95% confidence bounds)**

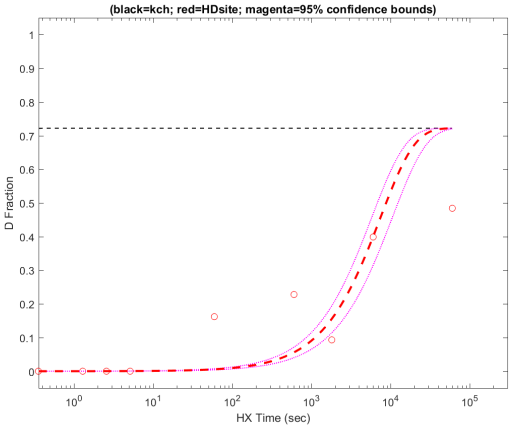


**Residue #73: HDsite kex=4.55e-04**
**(black=kch; red=HDsite; magenta=95% confidence bounds)**

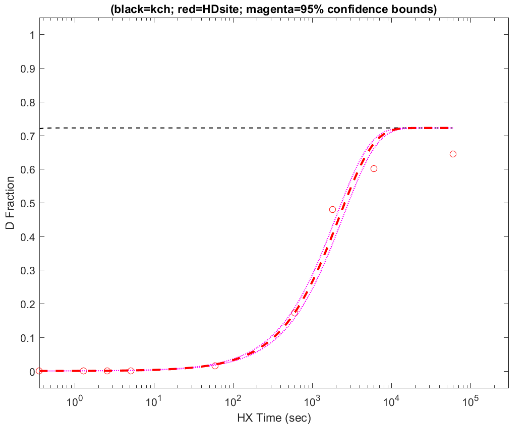


**Residue #74: HDsite kex=4.55e-04**
**(black=kch; red=HDsite; magenta=95% confidence bounds)**

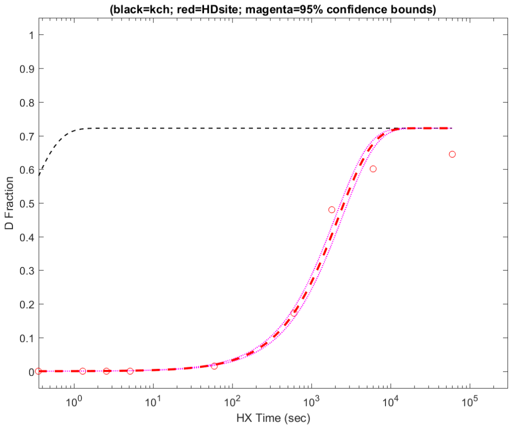

**Residue #75: HDsite kex=3.06e-04**
**(black=kch; red=HDsite; magenta=95% confidence bounds)**

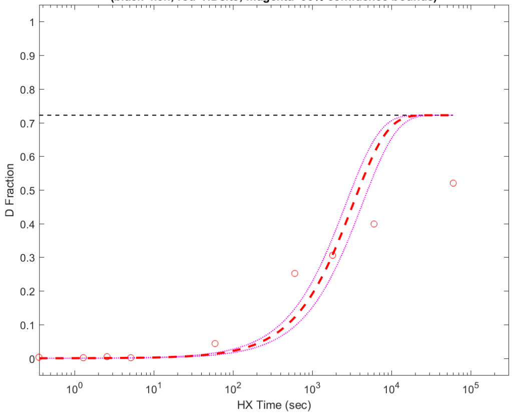


**Residue #76: HDsite kex=3.06e-04**
**(black=kch; red=HDsite; magenta=95% confidence bounds)**

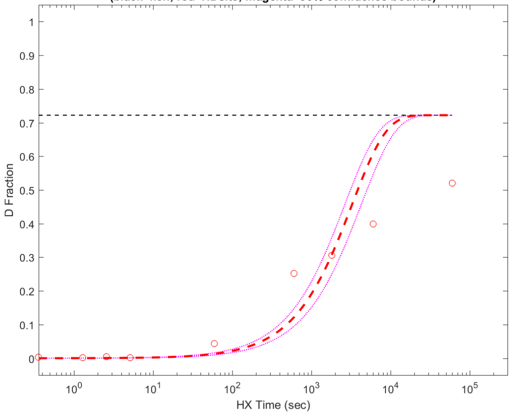


**Residue #77: HDsite kex=3.06e-04**
**(black=kch; red=HDsite; magenta=95% confidence bounds)**

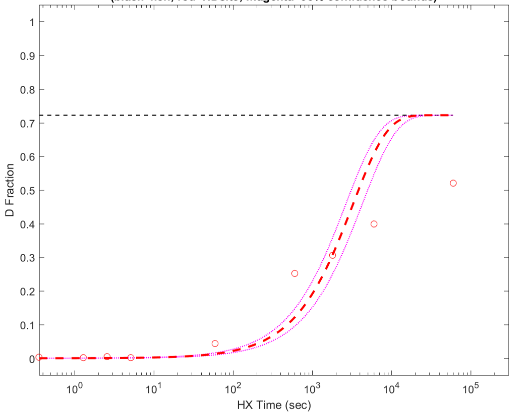


**Residue #78: HDsite kex=3.06e-04**
**(black=kch; red=HDsite; magenta=95% confidence bounds)**

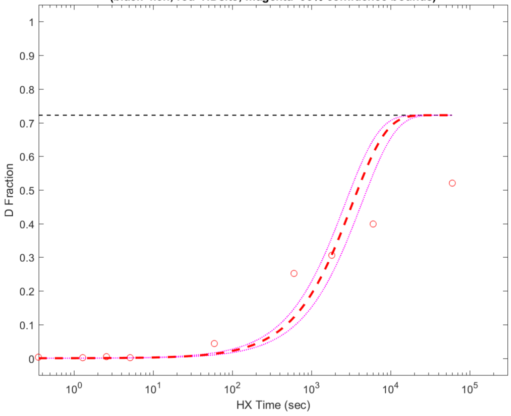


**Residue #79: HDsite kex=3.06e-04**
**(black=kch; red=HDsite; magenta=95% confidence bounds)**

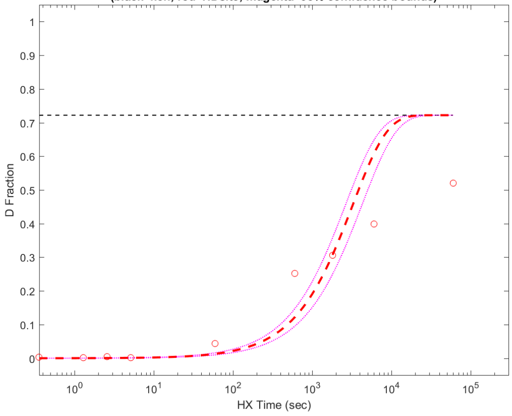


**Residue #80: HDsite kex=3.06e-04**
**(black=kch; red=HDsite; magenta=95% confidence bounds)**

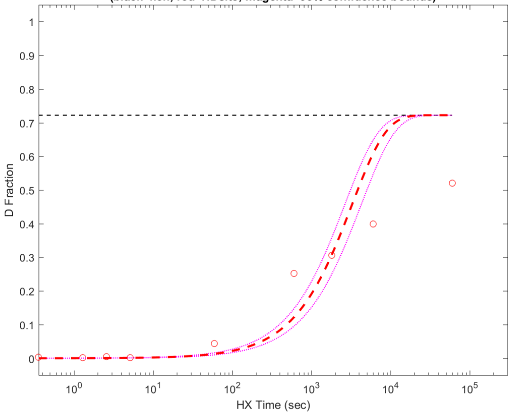


**Residue #81: HDsite kex=3.06e-04**
**(black=kch; red=HDsite; magenta=95% confidence bounds)**

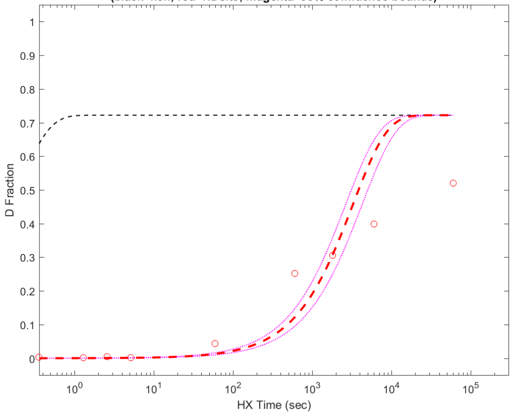


**Residue #82: HDsite kex=3.06e-04**
**(black=kch; red=HDsite; magenta=95% confidence bounds)**

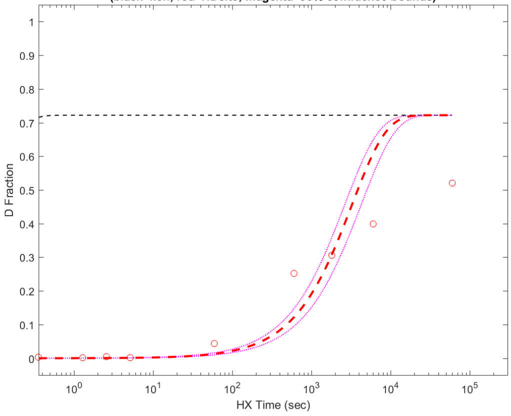

**Residue #83: HDsite kex=3.06e-04**
**(black=kch; red=HDsite; magenta=95% confidence bounds)**

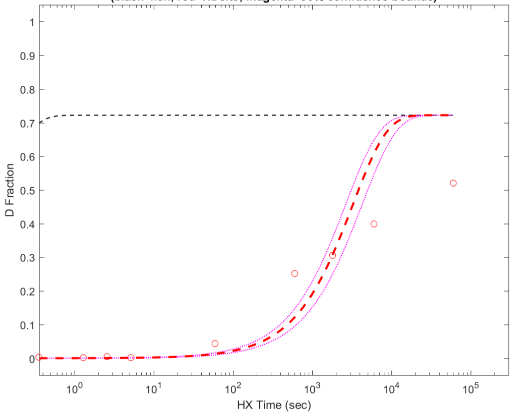


**Residue #84: HDsite kex=4.08e-04**
**(black=kch; red=HDsite; magenta=95% confidence bounds)**

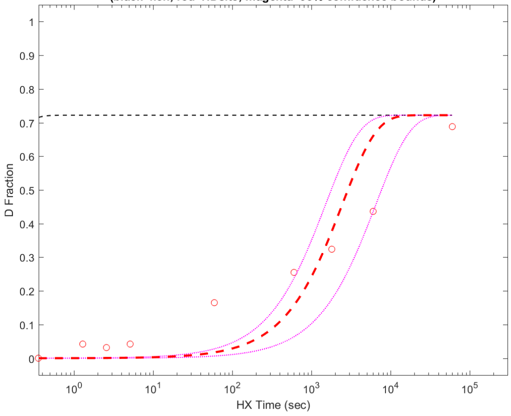


**Residue #85: HDsite kex=4.08e-04**
**(black=kch; red=HDsite; magenta=95% confidence bounds)**

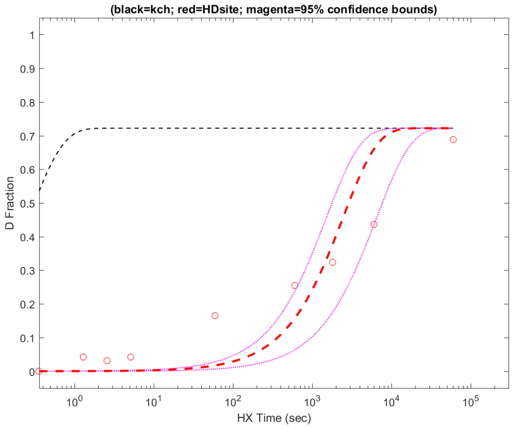


**Residue #86: HDsite kex=4.08e-04**
**(black=kch; red=HDsite; magenta=95% confidence bounds)**

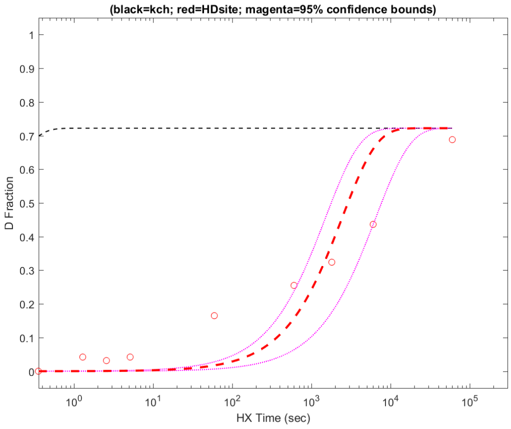


**Residue #87: HDsite kex=4.08e-04**
**(black=kch; red=HDsite; magenta=95% confidence bounds)**

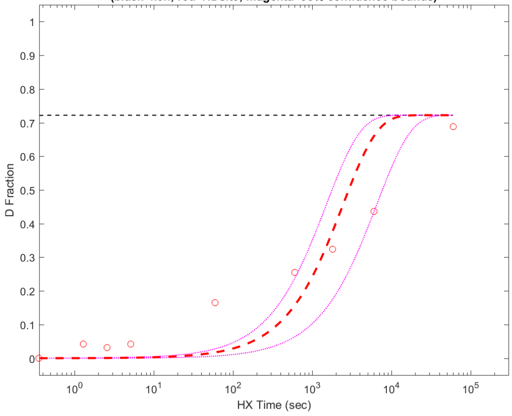


**Residue #88: HDsite kex=4.08e-04**
**(black=kch; red=HDsite; magenta=95% confidence bounds)**

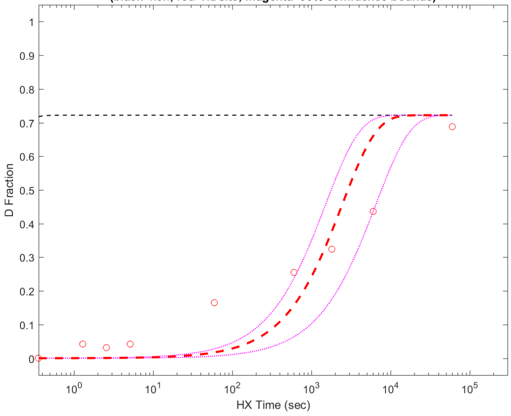


**Residue #89: HDsite kex=4.08e-04**
**(black=kch; red=HDsite; magenta=95% confidence bounds)**

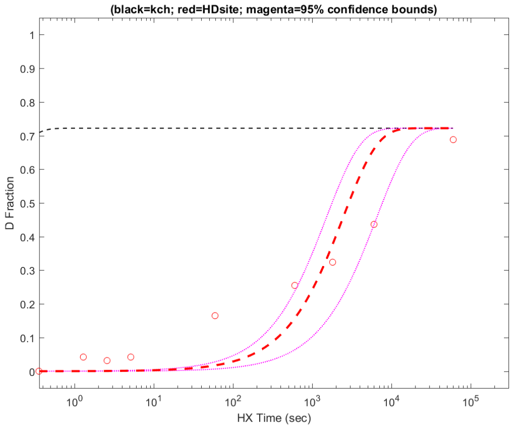


**Residue #90: HDsite kex=3.70e-04**
**(black=kch; red=HDsite; magenta=95% confidence bounds)**

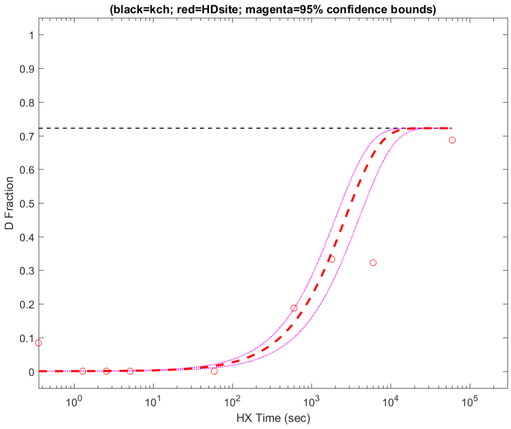

**Residue #91: HDsite kex=3.70e-04**
**(black=kch; red=HDsite; magenta=95% confidence bounds)**

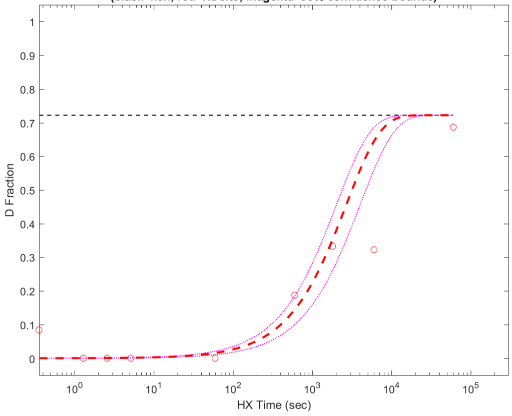


**Residue #92: HDsite kex=3.16e-04**
**(black=kch; red=HDsite; magenta=95% confidence bounds)**

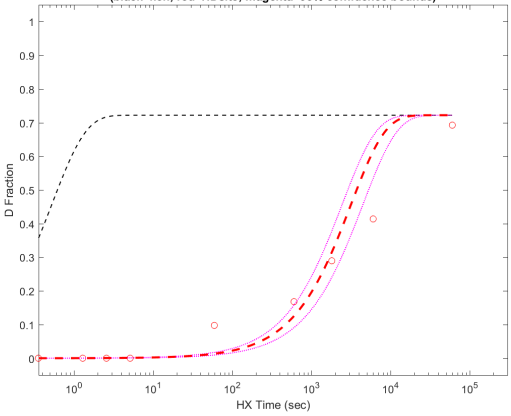


**Residue #93: HDsite kex=3.16e-04**
**(black=kch; red=HDsite; magenta=95% confidence bounds)**

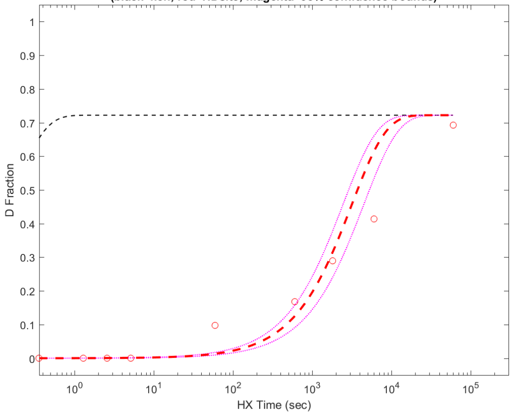


**Residue #94: HDsite kex=2.49e-04**
**(black=kch; red=HDsite; magenta=95% confidence bounds)**

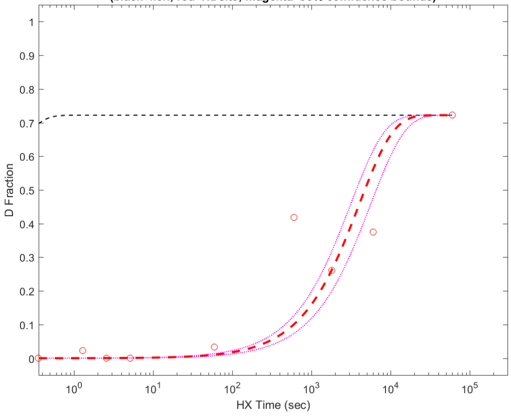


**Residue #95: HDsite kex=4.85e-06**
**(black=kch; red=HDsite; magenta=95% confidence bounds)**

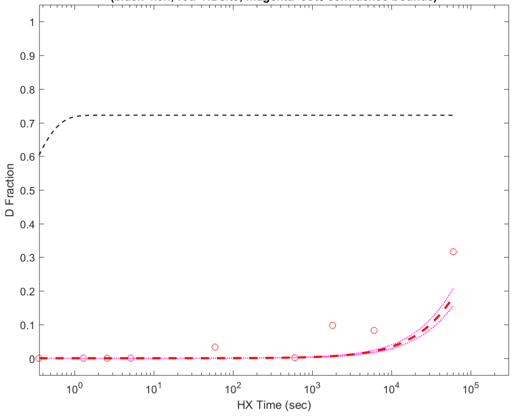


**Residue #96: HDsite kex=4.85e-06**
**(black=kch; red=HDsite; magenta=95% confidence bounds)**

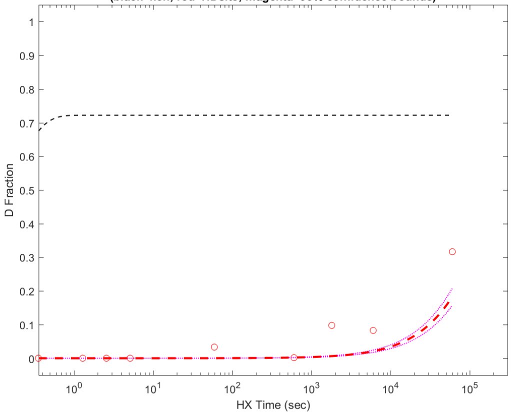


**Residue #97: HDsite kex=4.85e-06**
**(black=kch; red=HDsite; magenta=95% confidence bounds)**

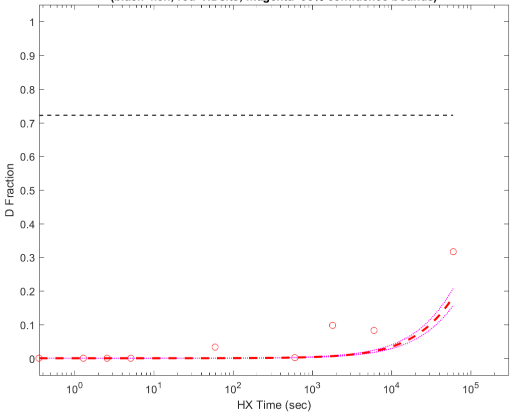


**Residue #98: HDsite kex=4.74e-05**
**(black=kch; red=HDsite; magenta=95% confidence bounds)**

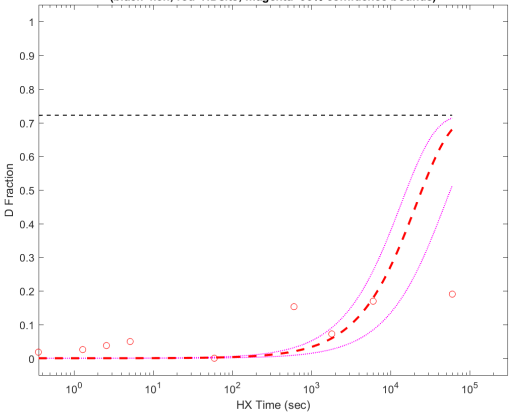

**Residue #99: HDsite kex=4.74e-05**
**(black=kch; red=HDsite; magenta=95% confidence bounds)**

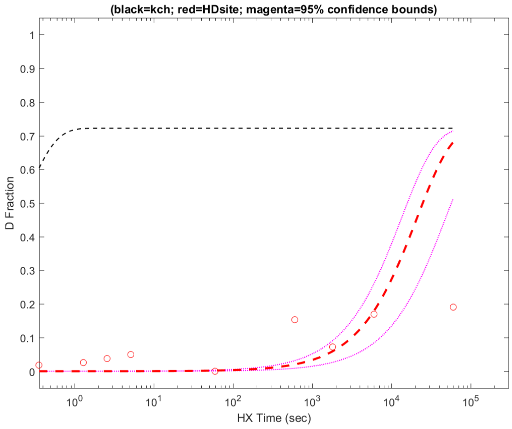


**Residue #100: HDsite kex=3.96e-05**
**(black=kch; red=HDsite; magenta=95% confidence bounds)**

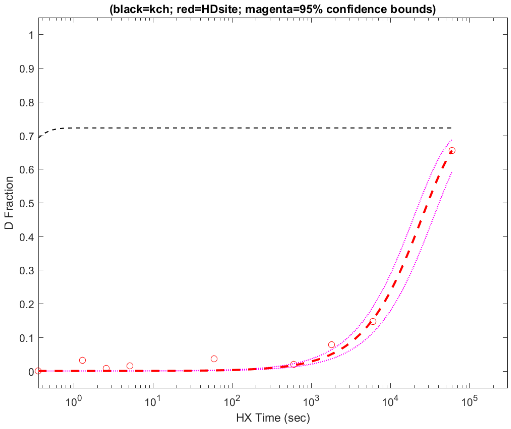


**Residue #101: HDsite kex=3.96e-05**
**(black=kch; red=HDsite; magenta=95% confidence bounds)**

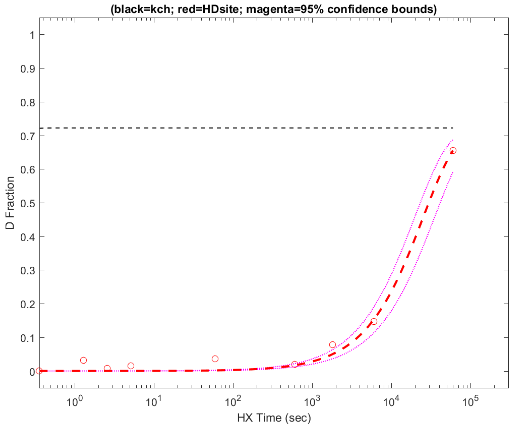


**Residue #102: HDsite kex=3.96e-05**
**(black=kch; red=HDsite; magenta=95% confidence bounds)**

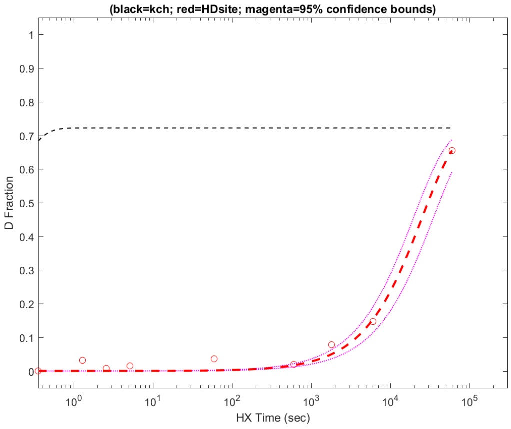


**Residue #103: HDsite kex=3.96e-05**
**(black=kch; red=HDsite; magenta=95% confidence bounds)**

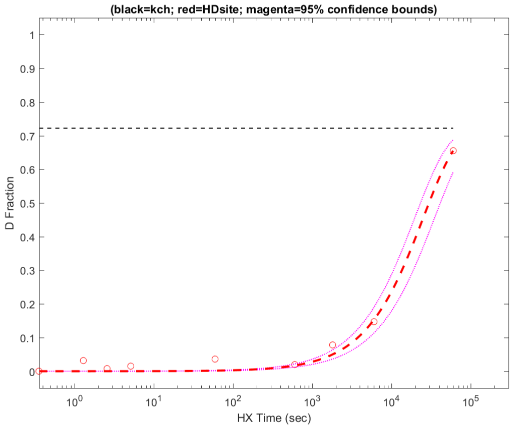


**Residue #104: HDsite kex=1.04e-06**
**(black=kch; red=HDsite; magenta=95% confidence bounds)**

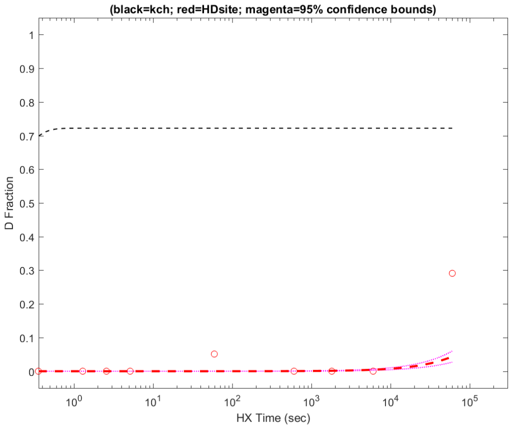


**Residue #105: HDsite kex=1.04e-06**
**(black=kch; red=HDsite; magenta=95% confidence bounds)**

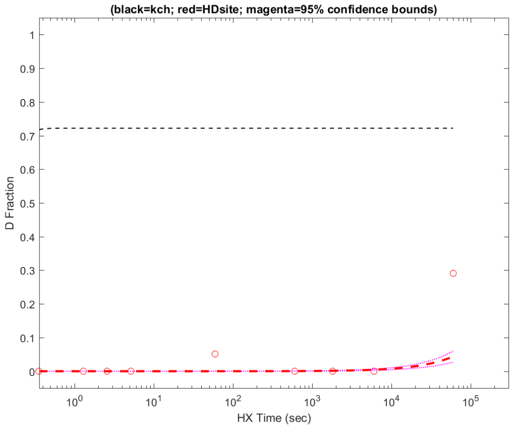


**Residue #106: HDsite kex=1.04e-06**
**(black=kch; red=HDsite; magenta=95% confidence bounds)**

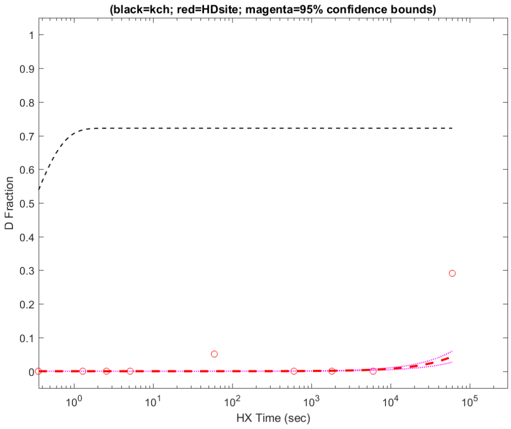

**Residue #109: HDsite kex=1.12e-04**
**(black=kch; red=HDsite; magenta=95% confidence bounds)**

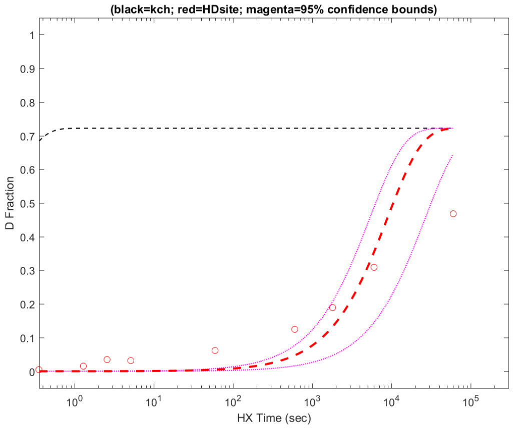


**Residue #110: HDsite kex=1.12e-04**
**(black=kch; red=HDsite; magenta=95% confidence bounds)**

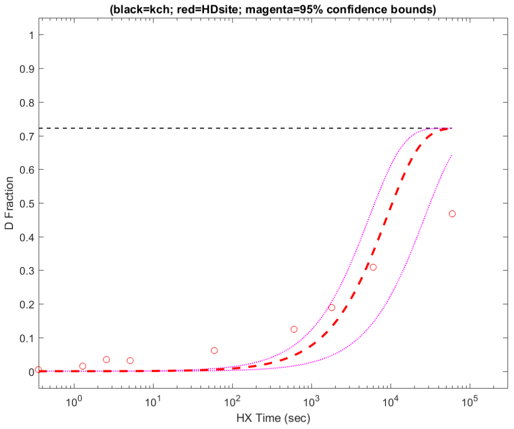


**Residue #111: HDsite kex=1.12e-04**
**(black=kch; red=HDsite; magenta=95% confidence bounds)**

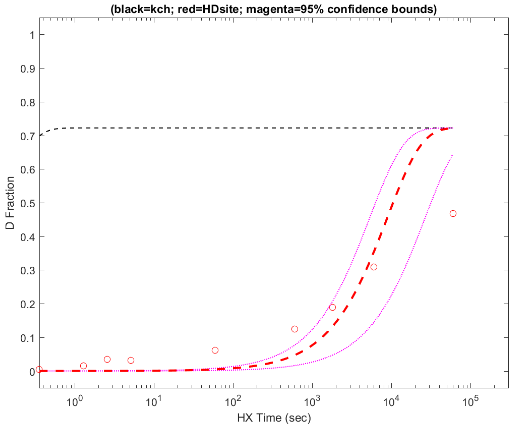


**Residue #112: HDsite kex=1.12e-04**
**(black=kch; red=HDsite; magenta=95% confidence bounds)**

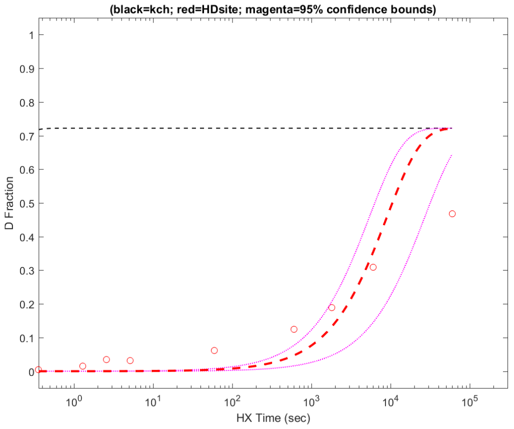


**Residue #113: HDsite kex=1.12e-04**
**(black=kch; red=HDsite; magenta=95% confidence bounds)**

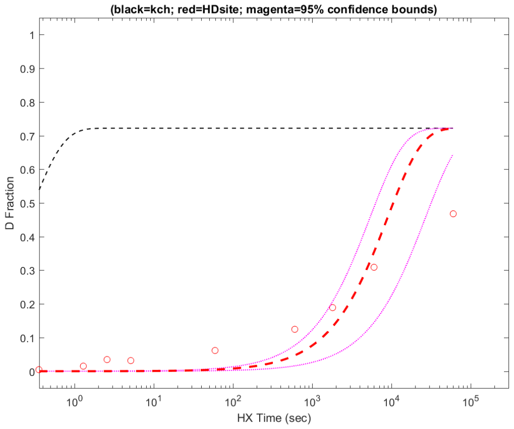


**Residue #114: HDsite kex=1.28e-06**
**(black=kch; red=HDsite; magenta=95% confidence bounds)**

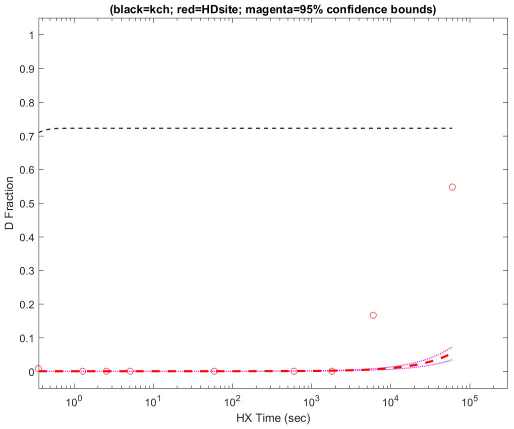


**Residue #115: HDsite kex=1.02e-03**
**(black=kch; red=HDsite; magenta=95% confidence bounds)**

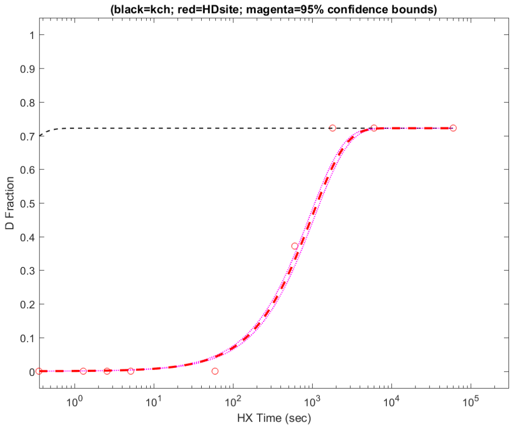


**Residue #116: HDsite kex=2.01e-04**
**(black=kch; red=HDsite; magenta=95% confidence bounds)**

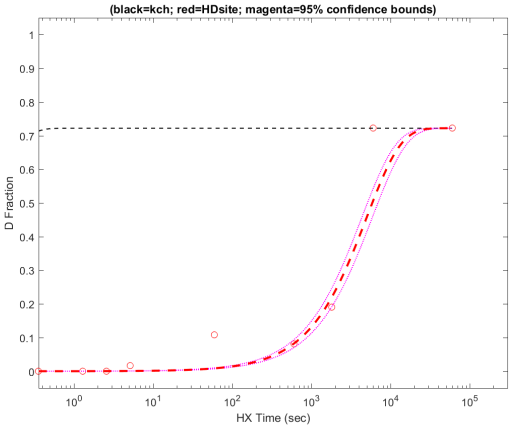

**Residue #117: HDsite kex=3.32e-03**
**(black=kch; red=HDsite; magenta=95% confidence bounds)**

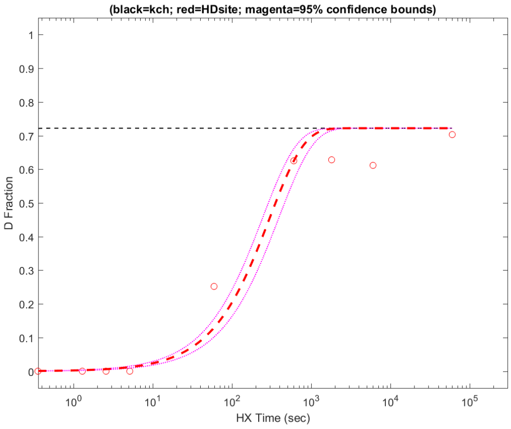


**Residue #118: HDsite kex=3.32e-03**
**(black=kch; red=HDsite; magenta=95% confidence bounds)**

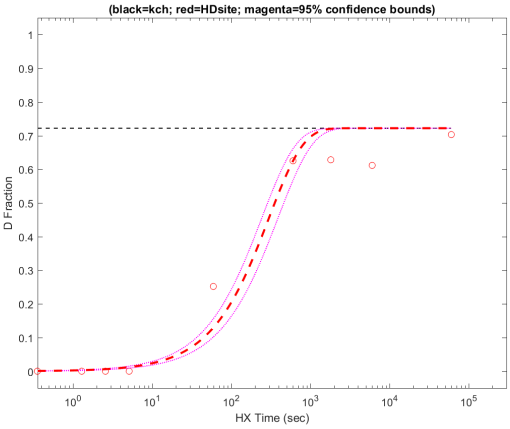


**Residue #119: HDsite kex=6.09e-04**
**(black=kch; red=HDsite; magenta=95% confidence bounds)**

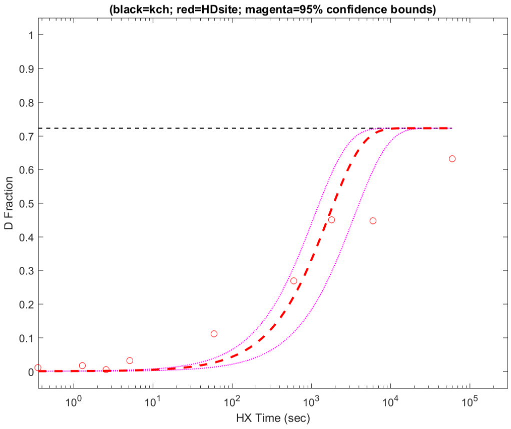


**Residue #120: HDsite kex=6.09e-04**
**(black=kch; red=HDsite; magenta=95% confidence bounds)**

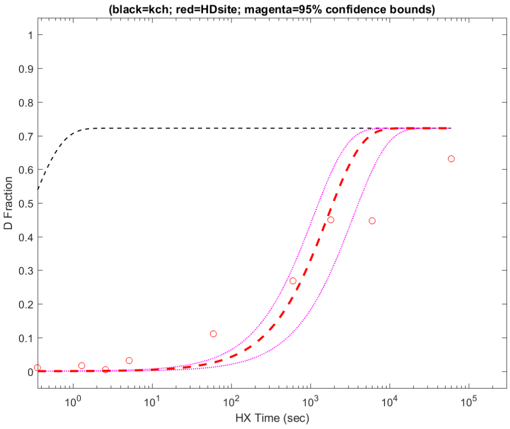


**Residue #121: HDsite kex=6.09e-04**
**(black=kch; red=HDsite; magenta=95% confidence bounds)**

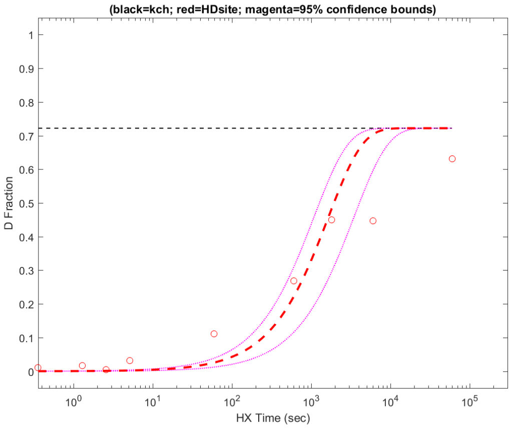


**Residue #122: HDsite kex=6.09e-04**
**(black=kch; red=HDsite; magenta=95% confidence bounds)**

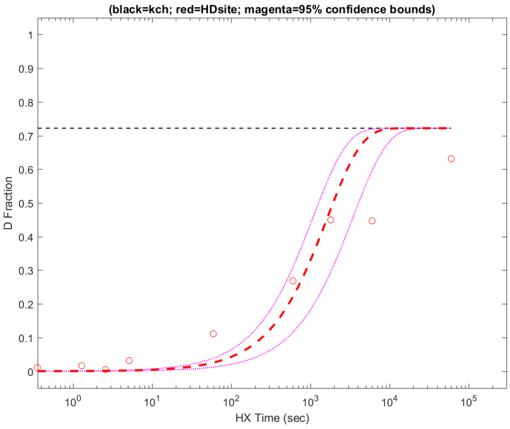


**Residue #123: HDsite kex=1.98e-03**
**(black=kch; red=HDsite; magenta=95% confidence bounds)**

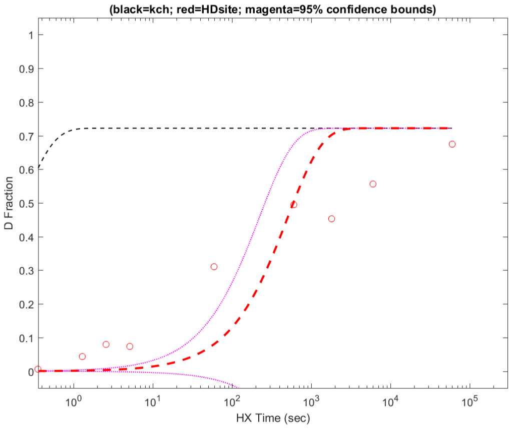


**Residue #124: HDsite kex=1.98e-03**
**(black=kch; red=HDsite; magenta=95% confidence bounds)**

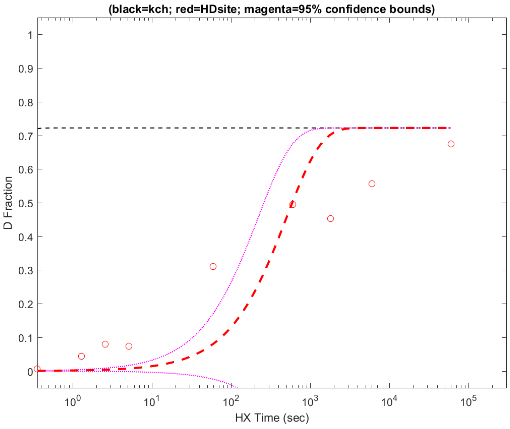

**Residue #125: HDsite kex=1.98e-03**
**(black=kch; red=HDsite; magenta=95% confidence bounds)**

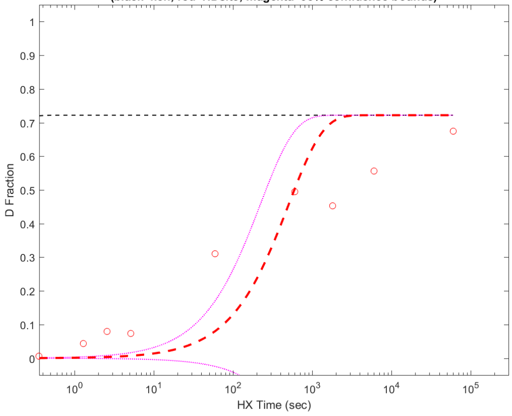


**Residue #126: HDsite kex=1.98e-03**
**(black=kch; red=HDsite; magenta=95% confidence bounds)**

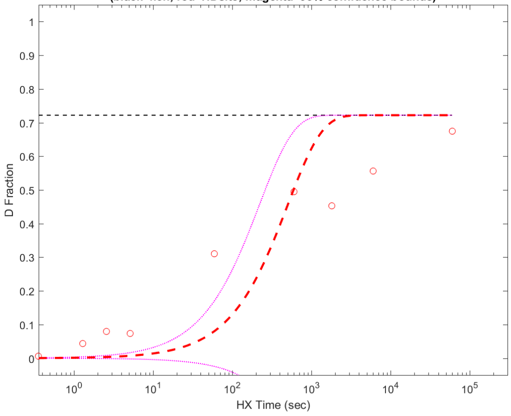


**Residue #127: HDsite kex=9.09e-03**
**(black=kch; red=HDsite; magenta=95% confidence bounds)**

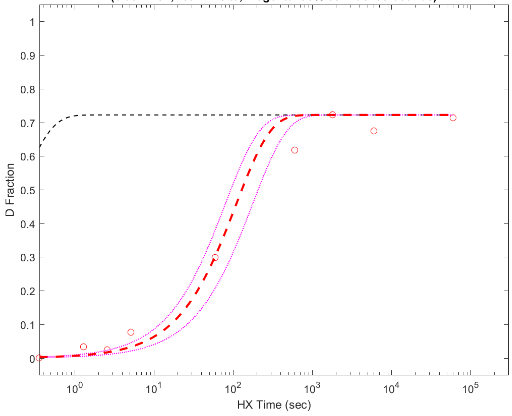


**Residue #128: HDsite kex=9.09e-03**
**(black=kch; red=HDsite; magenta=95% confidence bounds)**

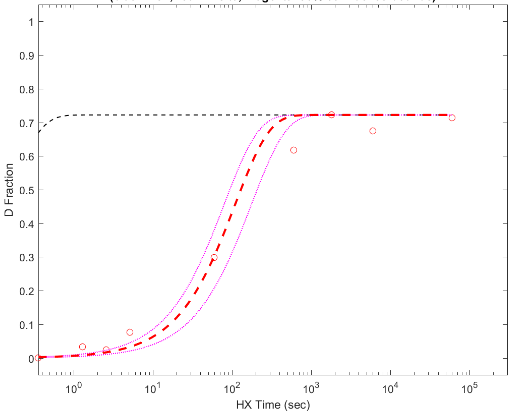


**Residue #129: HDsite kex=9.09e-03**
**(black=kch; red=HDsite; magenta=95% confidence bounds)**

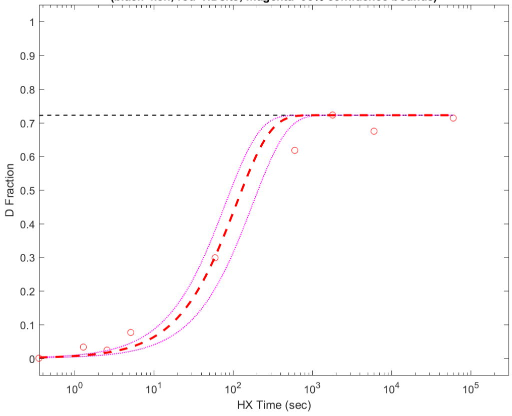


**Residue #130: HDsite kex=1.17e-02**
**(black=kch; red=HDsite; magenta=95% confidence bounds)**

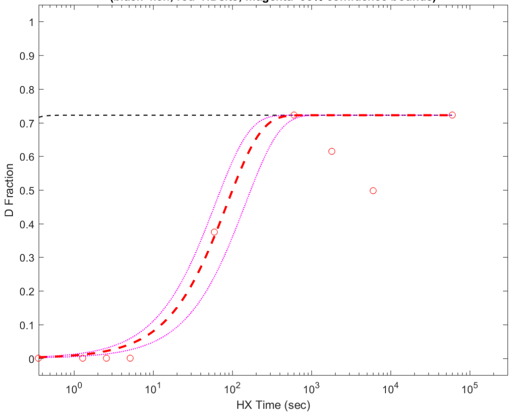


**Residue #132: HDsite kex=1.08e-03**
**(black=kch; red=HDsite; magenta=95% confidence bounds)**

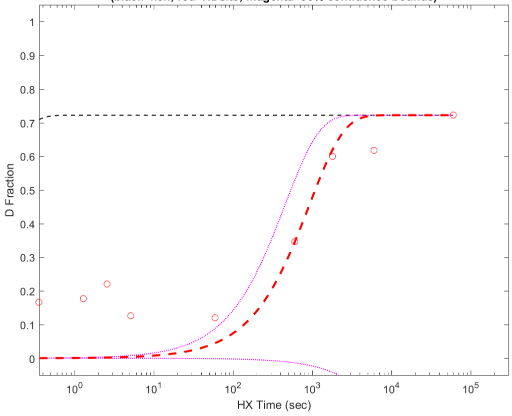


**Residue #133: HDsite kex=1.43e-05**
**(black=kch; red=HDsite; magenta=95% confidence bounds)**

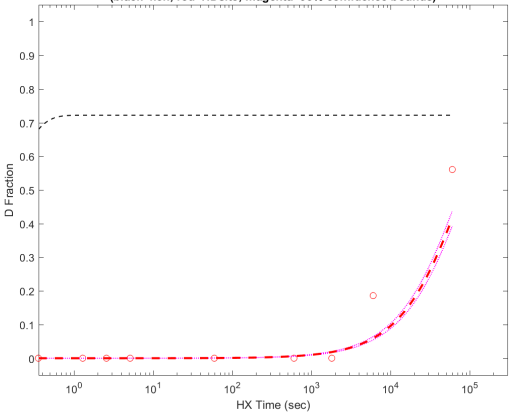

**Residue #134: HDsite kex=2.02e-04**
**(black=kch; red=HDsite; magenta=95% confidence bounds)**

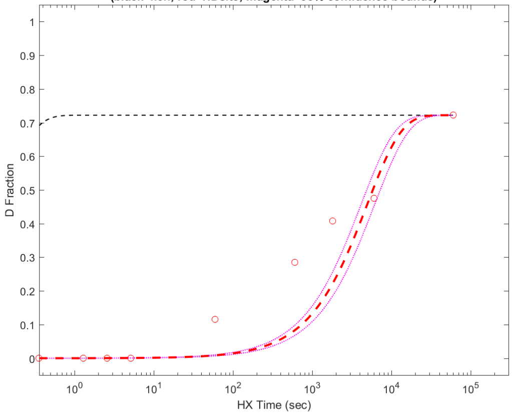


**Residue #135: HDsite kex=1.83e-04**
**(black=kch; red=HDsite; magenta=95% confidence bounds)**

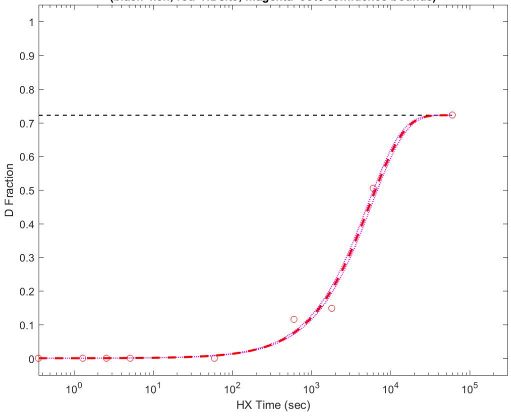


**Residue #136: HDsite kex=1.61e-04**
**(black=kch; red=HDsite; magenta=95% confidence bounds)**

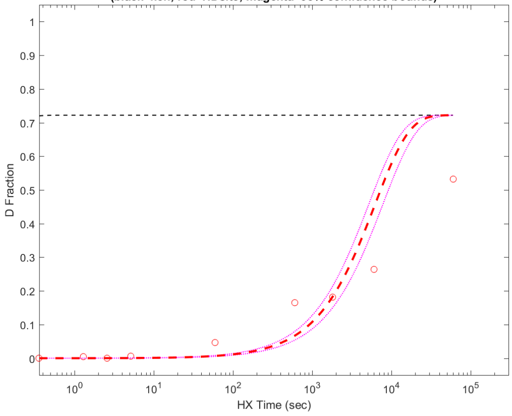


**Residue #137: HDsite kex=1.61e-04**
**(black=kch; red=HDsite; magenta=95% confidence bounds)**

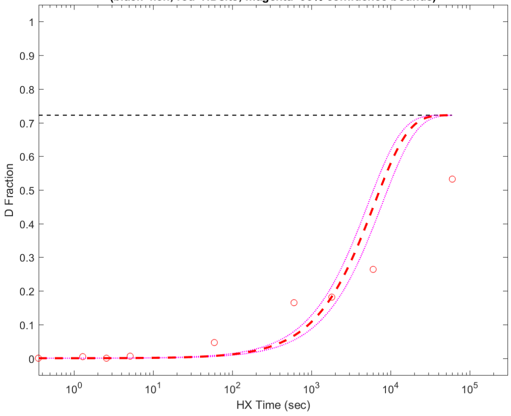


**Residue #138: HDsite kex=1.61e-04**
**(black=kch; red=HDsite; magenta=95% confidence bounds)**

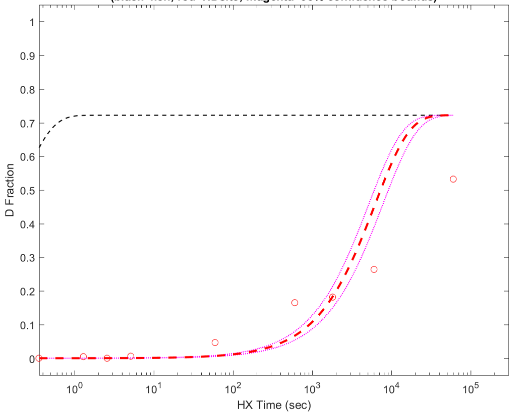


**Residue #139: HDsite kex=1.61e-04**
**(black=kch; red=HDsite; magenta=95% confidence bounds)**

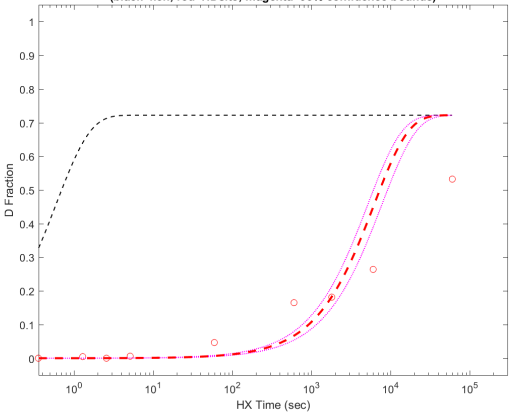


**Residue #140: HDsite kex=1.61e-04**
**(black=kch; red=HDsite; magenta=95% confidence bounds)**

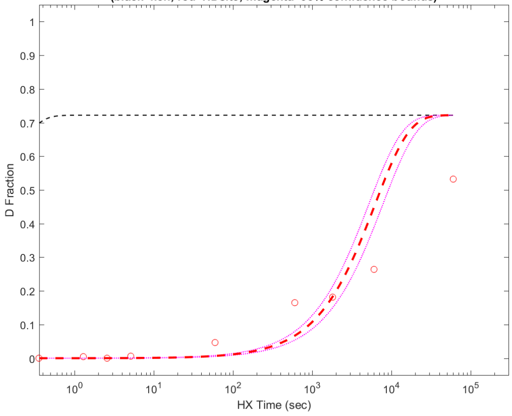


**Residue #141: HDsite kex=3.10e-06**
**(black=kch; red=HDsite; magenta=95% confidence bounds)**

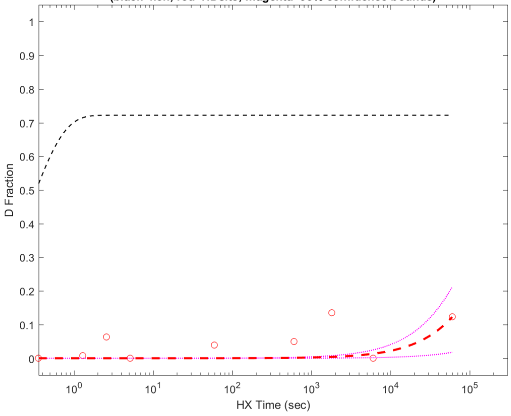

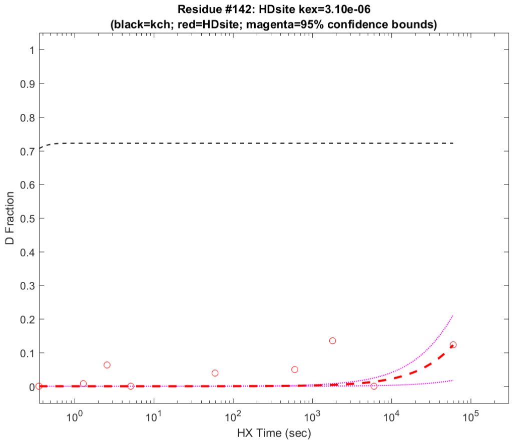
Residue #142: HDsite kex=3.10e-06
(black=kch; red=HDsite; magenta=95% confidence bounds)
D Fraction
HX Time (sec)

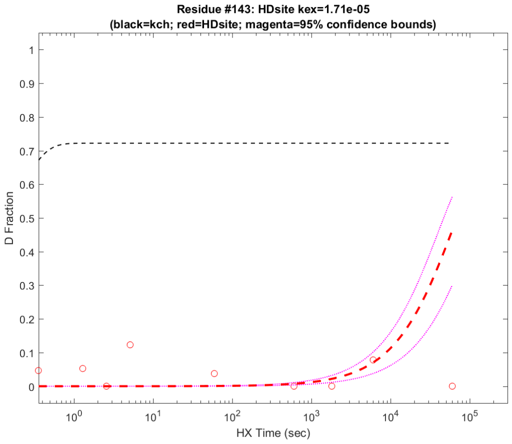
Residue #143: HDsite kex=1.71e-05
(black=kch; red=HDsite; magenta=95% confidence bounds)
D Fraction
HX Time (sec)

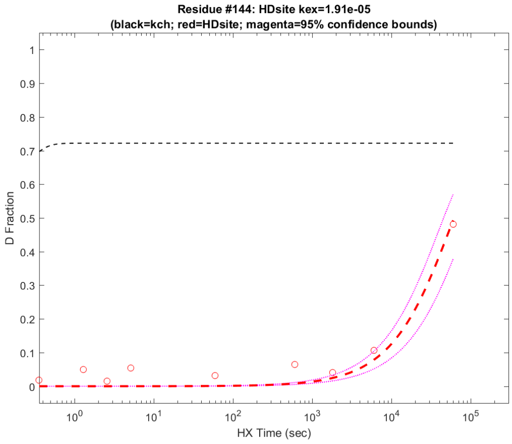
Residue #144: HDsite kex=1.91e-05
(black=kch; red=HDsite; magenta=95% confidence bounds)
D Fraction
HX Time (sec)

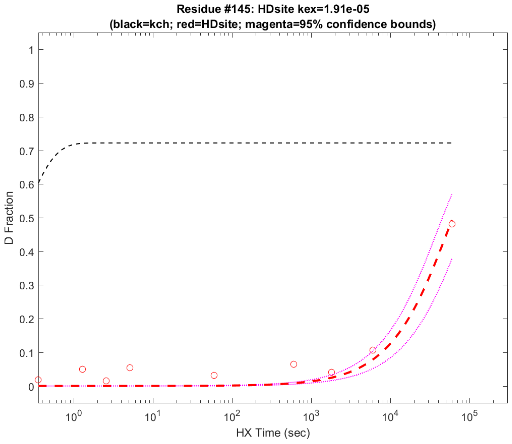
Residue #145: HDsite kex=1.91e-05
(black=kch; red=HDsite; magenta=95% confidence bounds)
D Fraction
HX Time (sec)

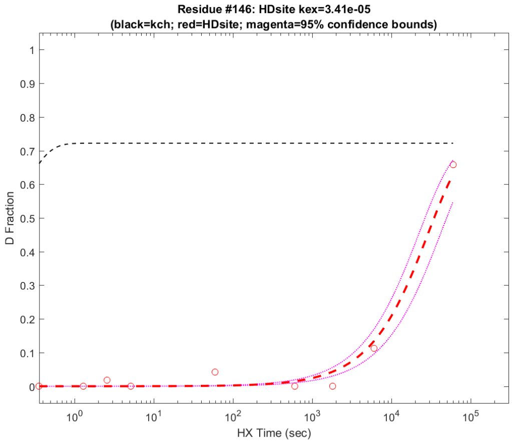
Residue #146: HDsite kex=3.41e-05
(black=kch; red=HDsite; magenta=95% confidence bounds)
D Fraction
HX Time (sec)

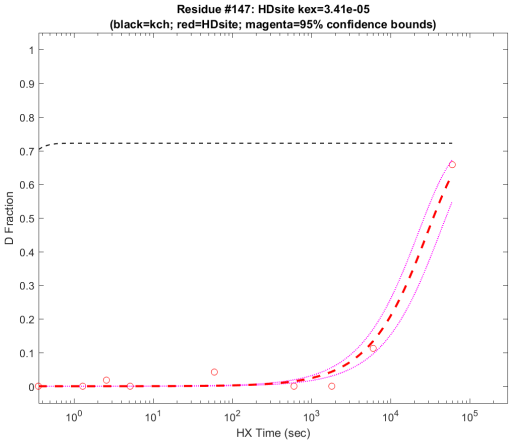
Residue #147: HDsite kex=3.41e-05
(black=kch; red=HDsite; magenta=95% confidence bounds)
D Fraction
HX Time (sec)

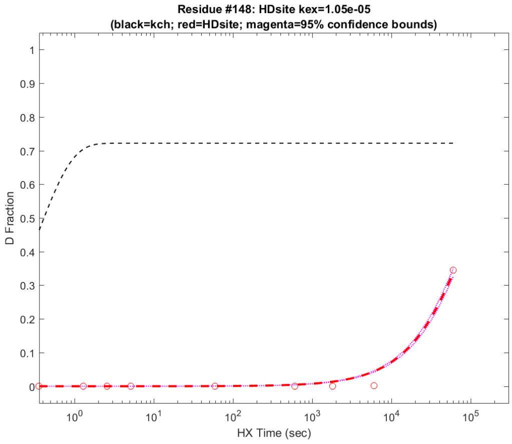
Residue #148: HDsite kex=1.05e-05
(black=kch; red=HDsite; magenta=95% confidence bounds)
D Fraction
HX Time (sec)

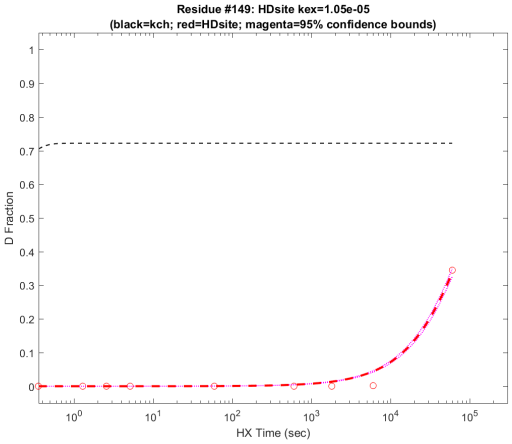
Residue #149: HDsite kex=1.05e-05
(black=kch; red=HDsite; magenta=95% confidence bounds)
D Fraction
HX Time (sec)

**Residue #150: HDsite kex=4.14e-04**
**(black=kch; red=HDsite; magenta=95% confidence bounds)**

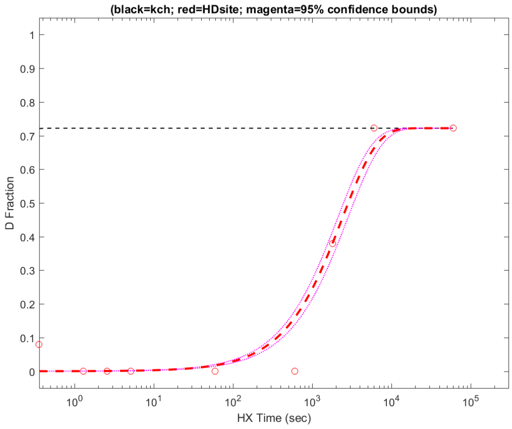


**Residue #151: HDsite kex=2.44e-04**
**(black=kch; red=HDsite; magenta=95% confidence bounds)**

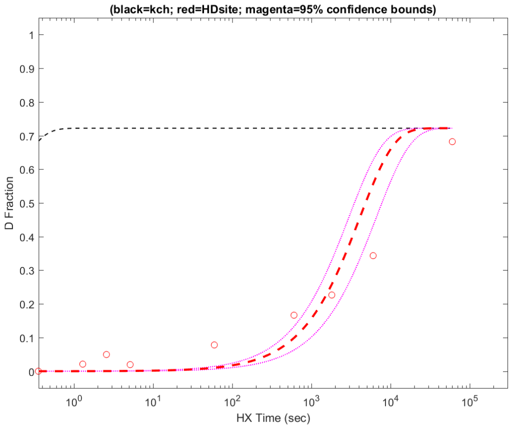


**Residue #152: HDsite kex=2.44e-04**
**(black=kch; red=HDsite; magenta=95% confidence bounds)**

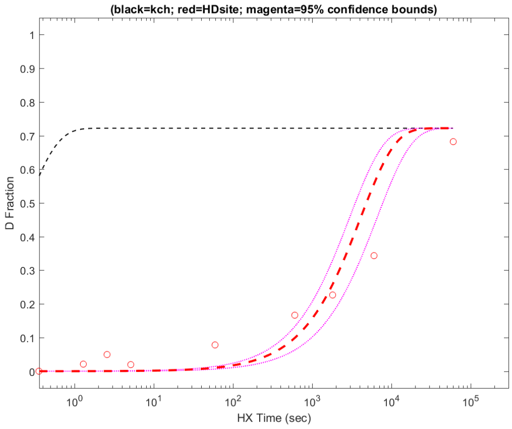


**Residue #153: HDsite kex=2.44e-04**
**(black=kch; red=HDsite; magenta=95% confidence bounds)**

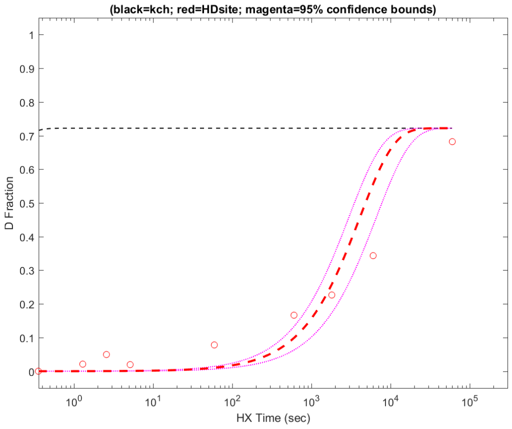


**Residue #154: HDsite kex=1.27e-04**
**(black=kch; red=HDsite; magenta=95% confidence bounds)**

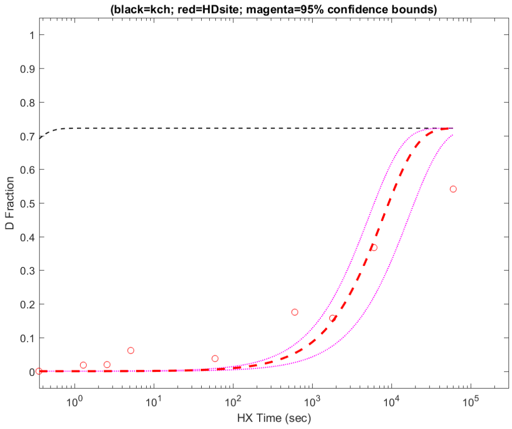


**Residue #155: HDsite kex=1.27e-04**
**(black=kch; red=HDsite; magenta=95% confidence bounds)**

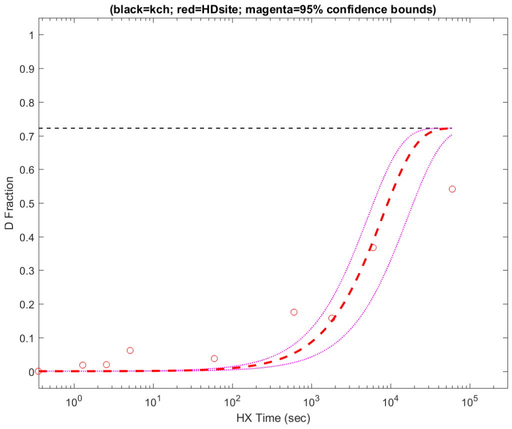


**Residue #156: HDsite kex=1.27e-04**
**(black=kch; red=HDsite; magenta=95% confidence bounds)**

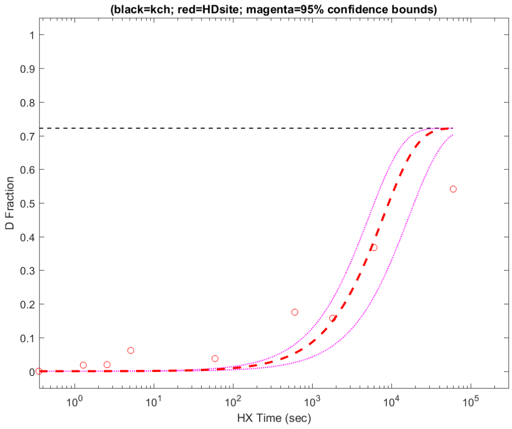


**Residue #157: HDsite kex=1.27e-04**
**(black=kch; red=HDsite; magenta=95% confidence bounds)**

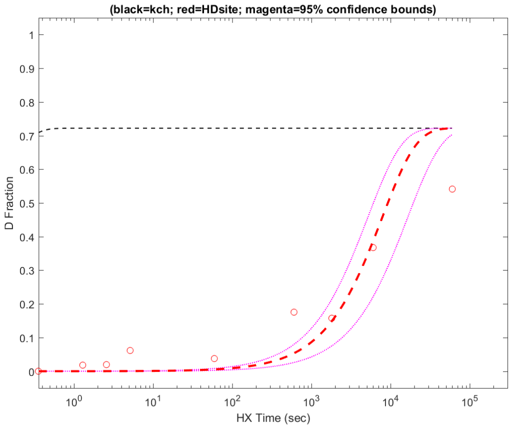

**Residue #158: HDsite kex=1.27e-04**
**(black=kch; red=HDsite; magenta=95% confidence bounds)**

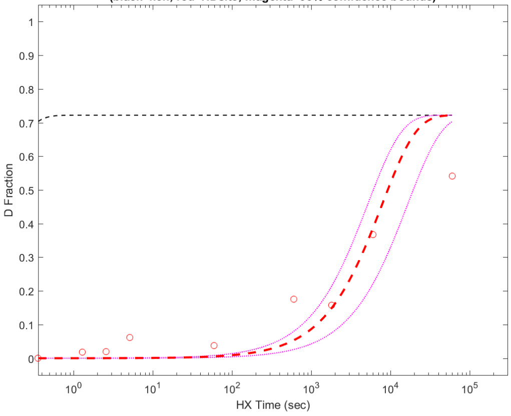


**Residue #159: HDsite kex=3.54e-04**
**(black=kch; red=HDsite; magenta=95% confidence bounds)**

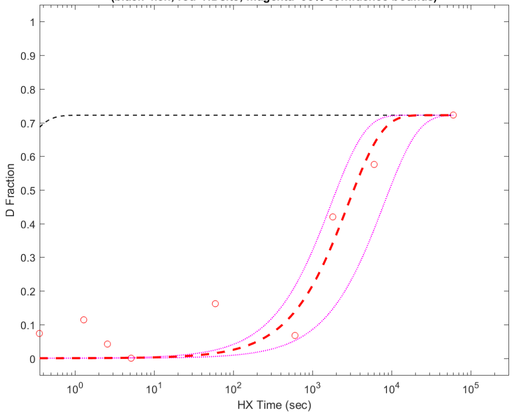


**Residue #160: HDsite kex=7.91e-01**
**(black=kch; red=HDsite; magenta=95% confidence bounds)**

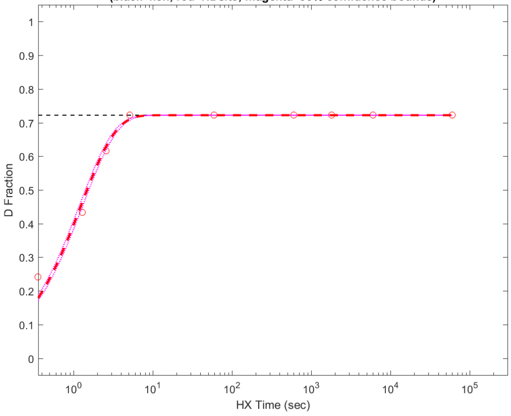


**Residue #161: HDsite kex=8.41e-03**
**(black=kch; red=HDsite; magenta=95% confidence bounds)**

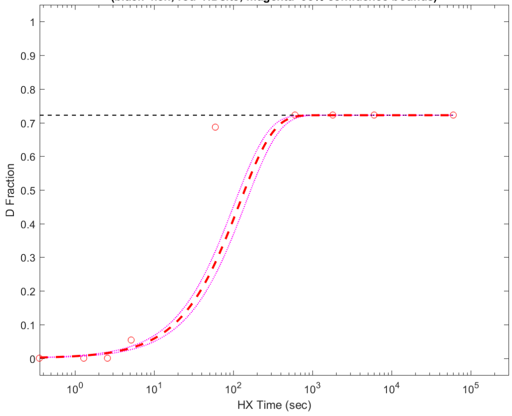


**Residue #162: HDsite kex=5.90e-03**
**(black=kch; red=HDsite; magenta=95% confidence bounds)**

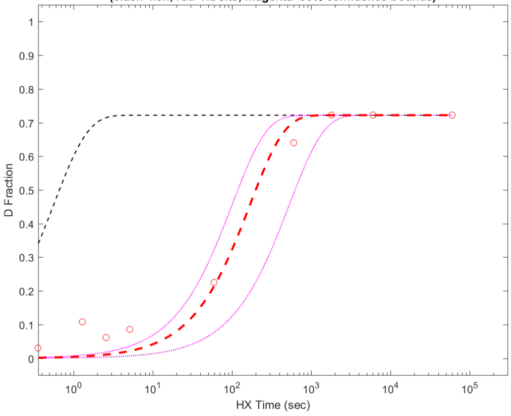


**Residue #163: HDsite kex=5.90e-03**
**(black=kch; red=HDsite; magenta=95% confidence bounds)**

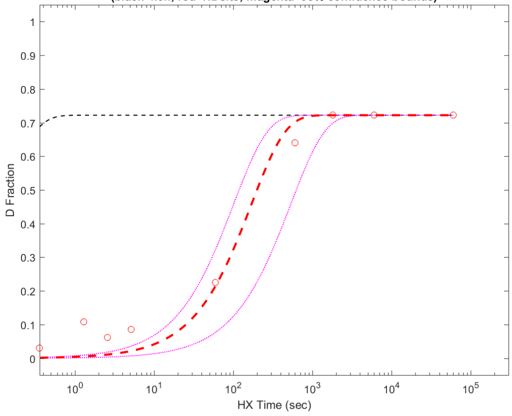


**Residue #164: HDsite kex=1.35e-03**
**(black=kch; red=HDsite; magenta=95% confidence bounds)**

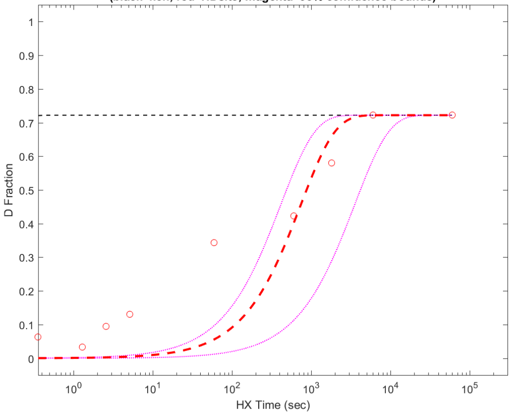


**Residue #165: HDsite kex=1.35e-03**
**(black=kch; red=HDsite; magenta=95% confidence bounds)**

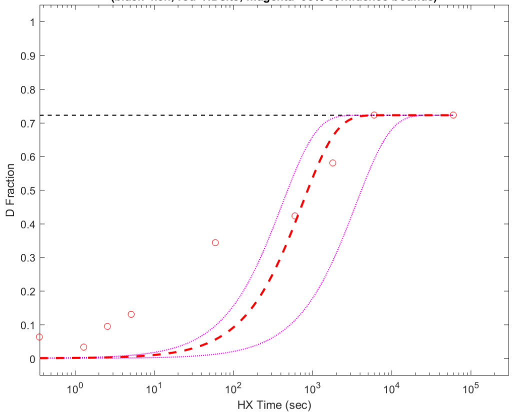

**Residue #166: HDsite kex=1.35e-03**
**(black=kch; red=HDsite; magenta=95% confidence bounds)**

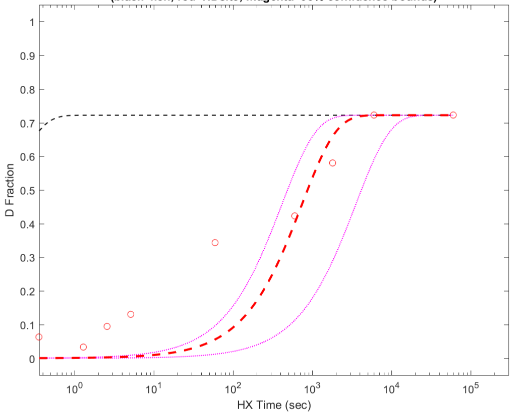


**Residue #167: HDsite kex=2.71e-03**
**(black=kch; red=HDsite; magenta=95% confidence bounds)**

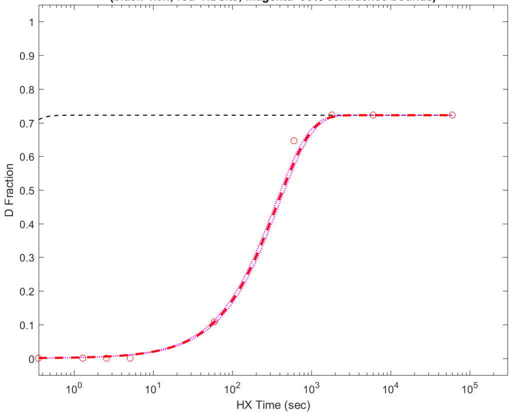


**Residue #168: HDsite kex=2.71e-03**
**(black=kch; red=HDsite; magenta=95% confidence bounds)**

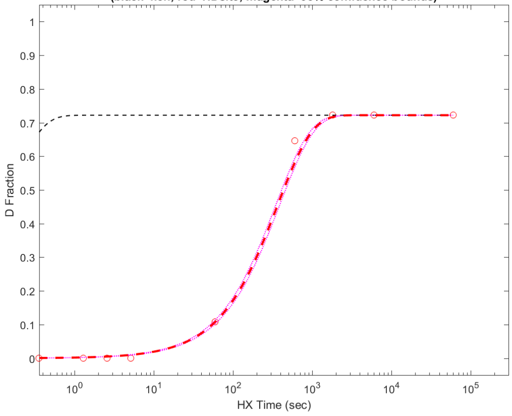


**Residue #169: HDsite kex=1.53e-04**
**(black=kch; red=HDsite; magenta=95% confidence bounds)**

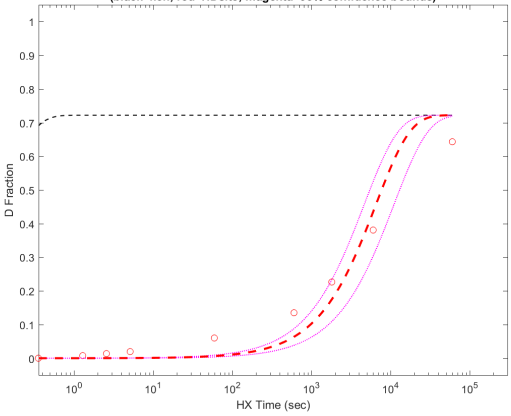


**Residue #170: HDsite kex=1.53e-04**
**(black=kch; red=HDsite; magenta=95% confidence bounds)**

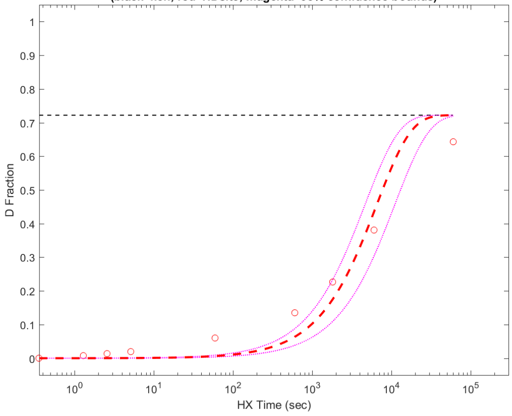


**Residue #171: HDsite kex=1.53e-04**
**(black=kch; red=HDsite; magenta=95% confidence bounds)**

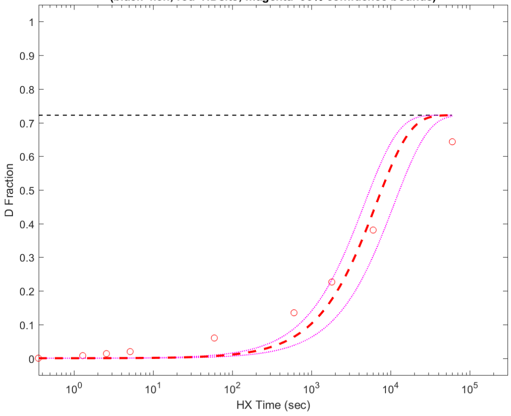


**Residue #172: HDsite kex=1.53e-04**
**(black=kch; red=HDsite; magenta=95% confidence bounds)**

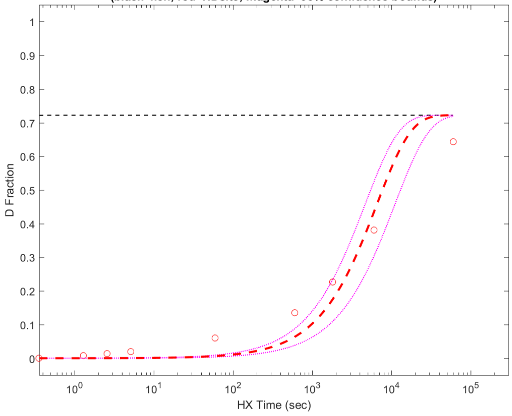


**Residue #173: HDsite kex=1.53e-04**
**(black=kch; red=HDsite; magenta=95% confidence bounds)**

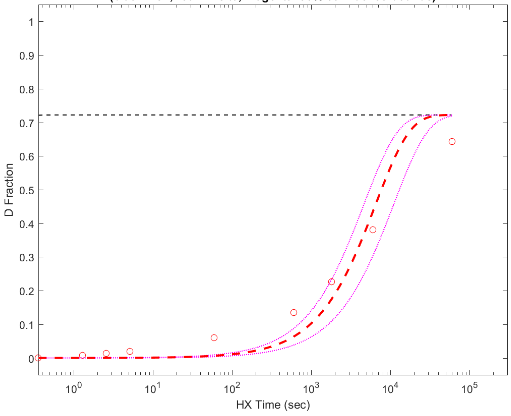

**Residue #174: HDsite kex=1.53e-04**
**(black=kch; red=HDsite; magenta=95% confidence bounds)**

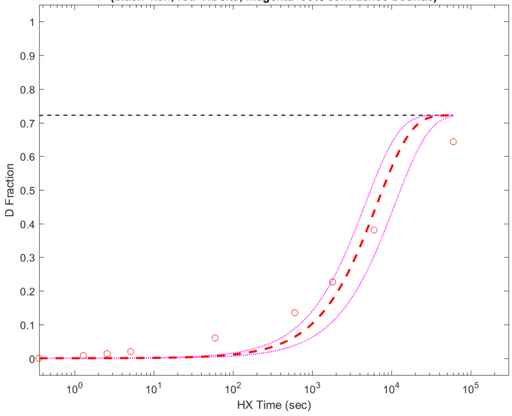


**Residue #175: HDsite kex=1.53e-04**
**(black=kch; red=HDsite; magenta=95% confidence bounds)**

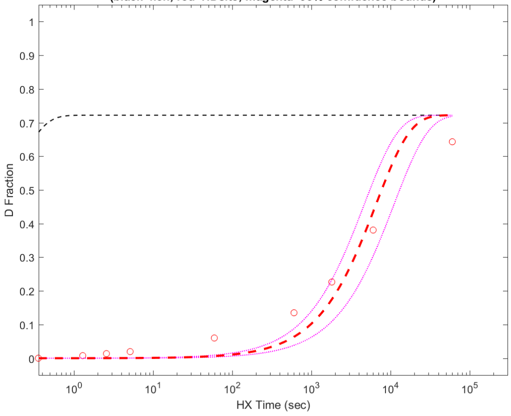


**Residue #176: HDsite kex=8.56e-07**
**(black=kch; red=HDsite; magenta=95% confidence bounds)**

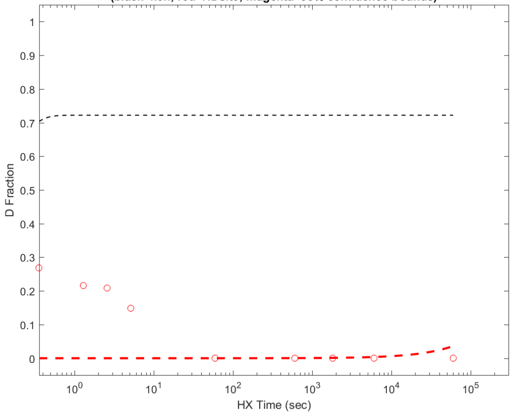


**Residue #177: HDsite kex=8.55e-07**
**(black=kch; red=HDsite; magenta=95% confidence bounds)**

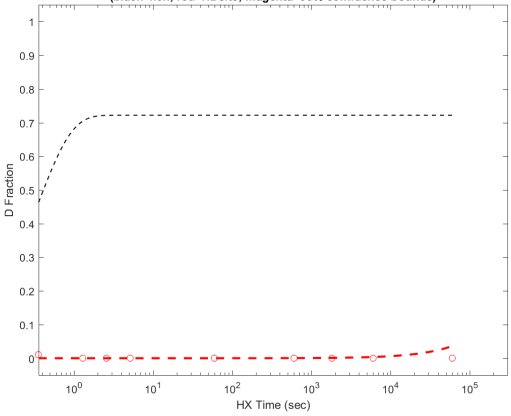


**Residue #178: HDsite kex=8.72e-07**
**(black=kch; red=HDsite; magenta=95% confidence bounds)**

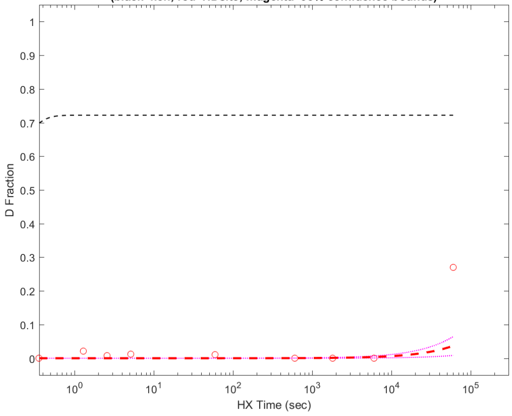


**Residue #179: HDsite kex=8.72e-07**
**(black=kch; red=HDsite; magenta=95% confidence bounds)**

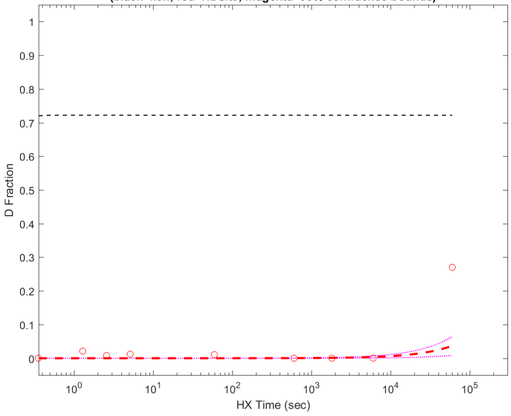


**Residue #180: HDsite kex=8.72e-07**
**(black=kch; red=HDsite; magenta=95% confidence bounds)**

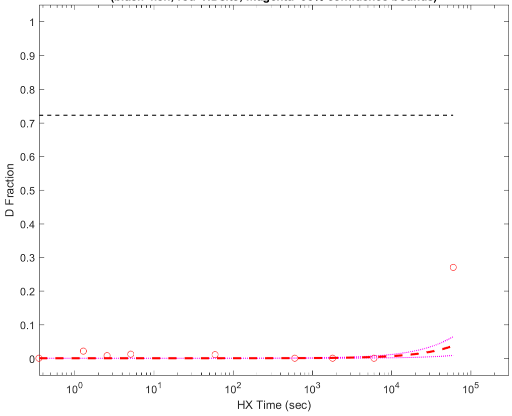


**Residue #181: HDsite kex=8.72e-07**
**(black=kch; red=HDsite; magenta=95% confidence bounds)**

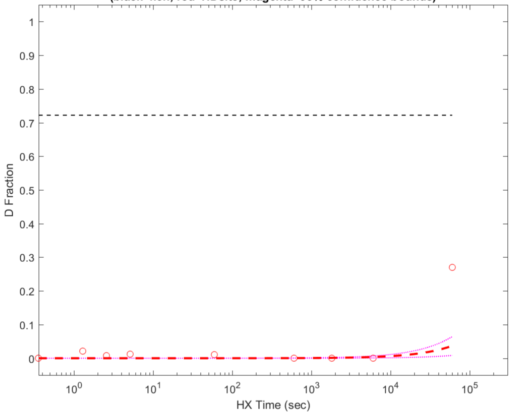

**Residue #182: HDsite kex=8.72e-07**
**(black=kch; red=HDsite; magenta=95% confidence bounds)**

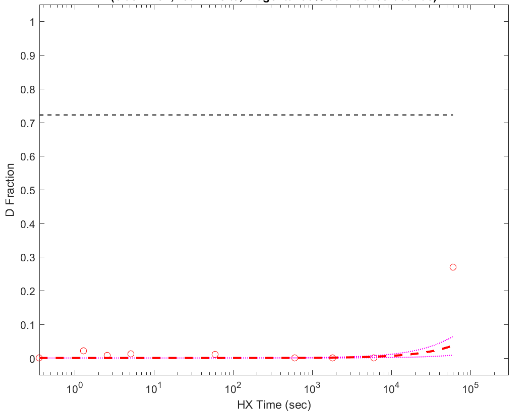


**Residue #183: HDsite kex=6.90e-05**
**(black=kch; red=HDsite; magenta=95% confidence bounds)**

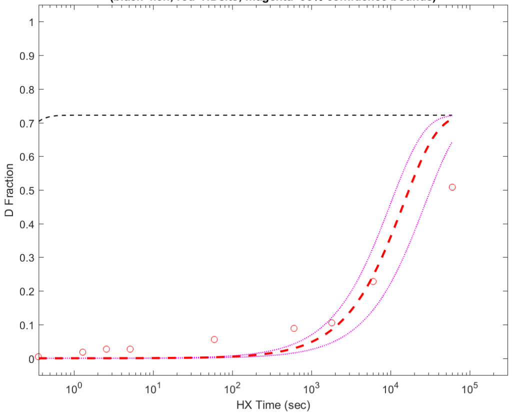


**Residue #184: HDsite kex=6.90e-05**
**(black=kch; red=HDsite; magenta=95% confidence bounds)**

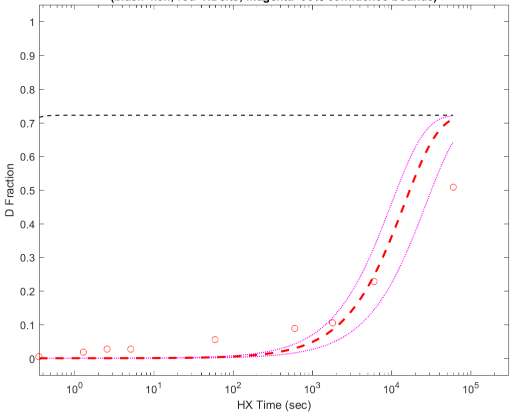


**Residue #185: HDsite kex=6.90e-05**
**(black=kch; red=HDsite; magenta=95% confidence bounds)**

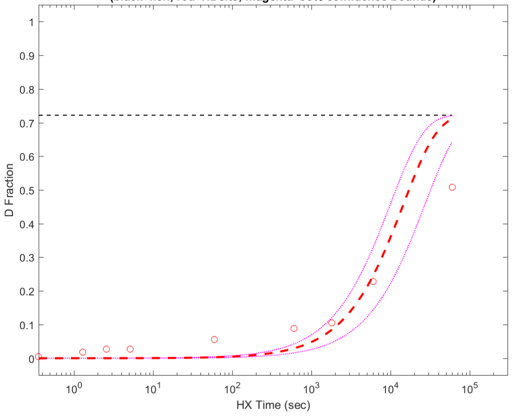


**Residue #186: HDsite kex=6.90e-05**
**(black=kch; red=HDsite; magenta=95% confidence bounds)**

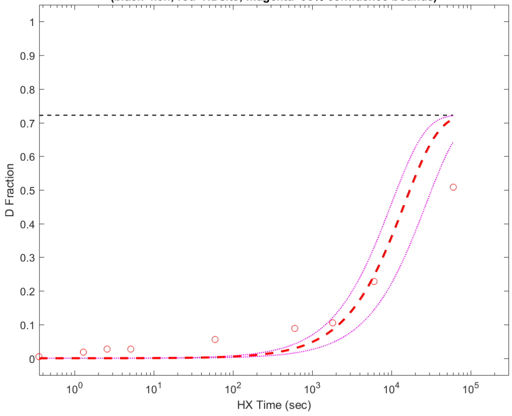


**Residue #187: HDsite kex=6.90e-05**
**(black=kch; red=HDsite; magenta=95% confidence bounds)**

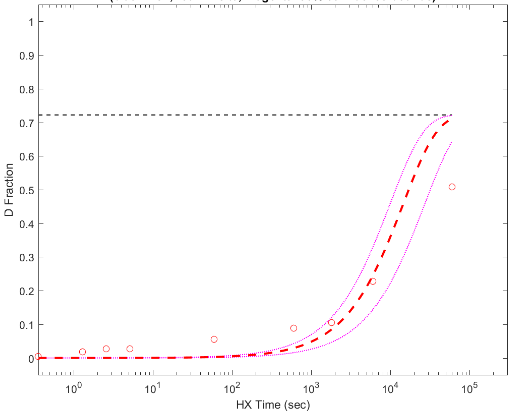


**Residue #188: HDsite kex=6.90e-05**
**(black=kch; red=HDsite; magenta=95% confidence bounds)**

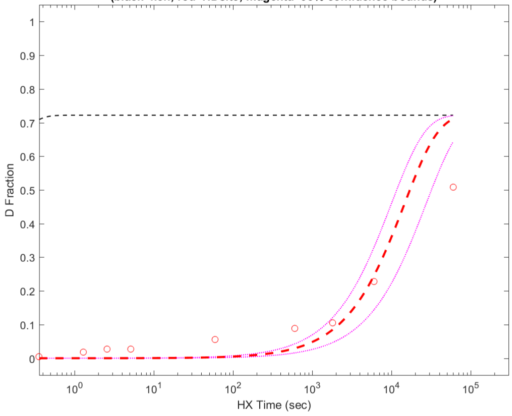


**Residue #189: HDsite kex=1.70e-05**
**(black=kch; red=HDsite; magenta=95% confidence bounds)**

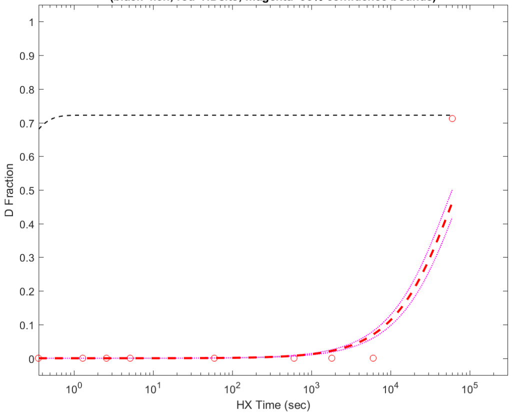

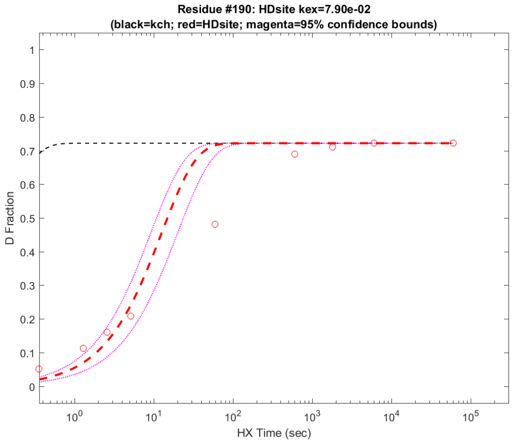
Residue #190: HDsite kex=7.90e-02
(black=kch; red=HDsite; magenta=95% confidence bounds)
D Fraction
HX Time (sec)

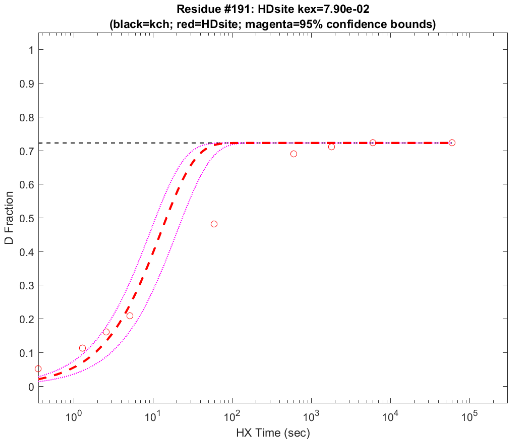
Residue #191: HDsite kex=7.90e-02
(black=kch; red=HDsite; magenta=95% confidence bounds)
D Fraction
HX Time (sec)

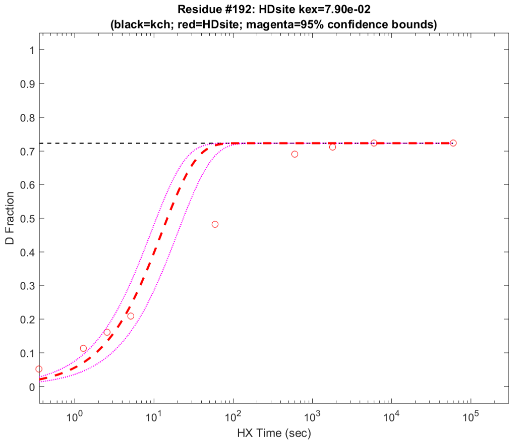
Residue #192: HDsite kex=7.90e-02
(black=kch; red=HDsite; magenta=95% confidence bounds)
D Fraction
HX Time (sec)

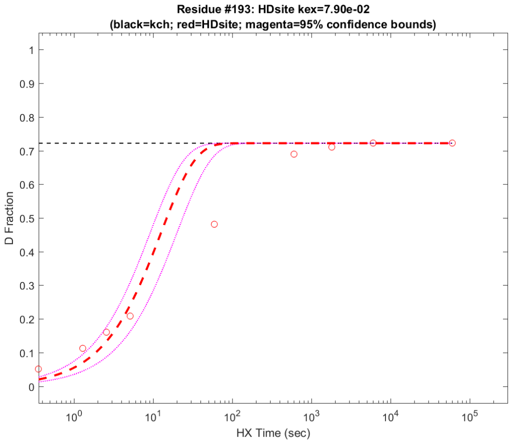
Residue #193: HDsite kex=7.90e-02
(black=kch; red=HDsite; magenta=95% confidence bounds)
D Fraction
HX Time (sec)

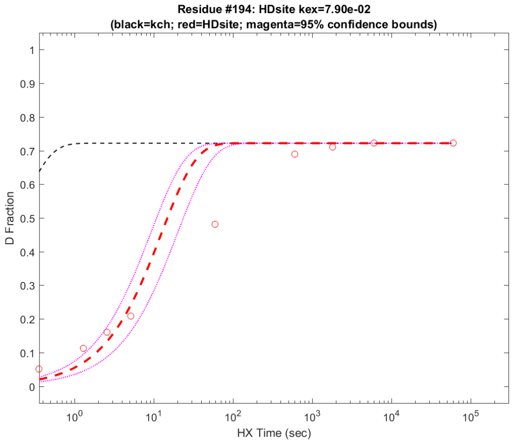
Residue #194: HDsite kex=7.90e-02
(black=kch; red=HDsite; magenta=95% confidence bounds)
D Fraction
HX Time (sec)

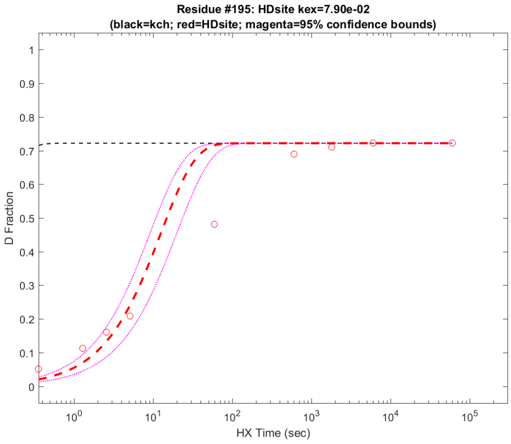
Residue #195: HDsite kex=7.90e-02
(black=kch; red=HDsite; magenta=95% confidence bounds)
D Fraction
HX Time (sec)

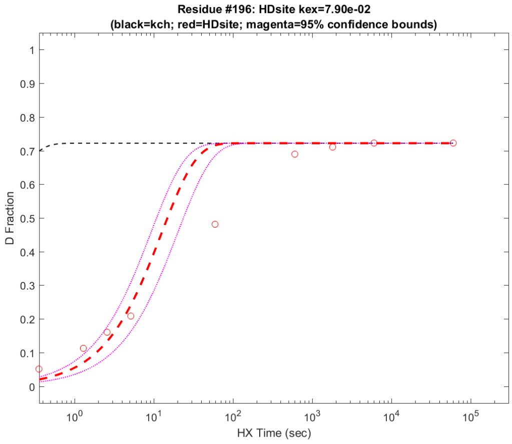
Residue #196: HDsite kex=7.90e-02
(black=kch; red=HDsite; magenta=95% confidence bounds)
D Fraction
HX Time (sec)

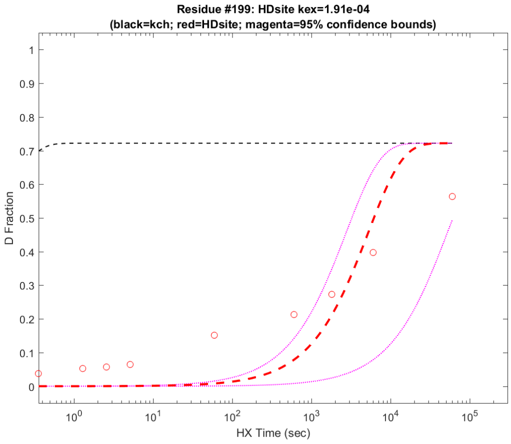
Residue #199: HDsite kex=1.91e-04
(black=kch; red=HDsite; magenta=95% confidence bounds)
D Fraction
HX Time (sec)

**Residue #200: HDsite kex=1.91e-04**
**(black=kch; red=HDsite; magenta=95% confidence bounds)**

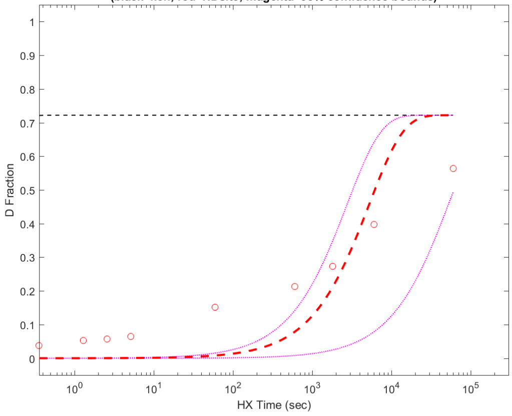


**Residue #201: HDsite kex=1.91e-04**
**(black=kch; red=HDsite; magenta=95% confidence bounds)**

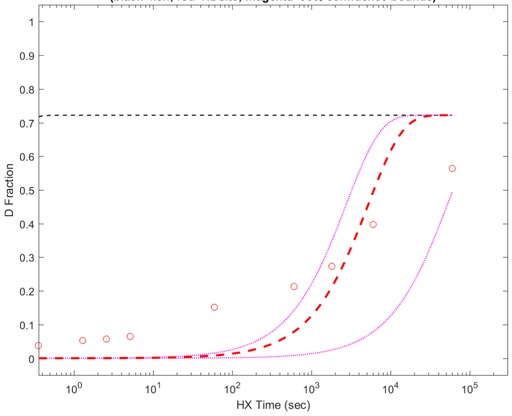


**Residue #202: HDsite kex=1.91e-04**
**(black=kch; red=HDsite; magenta=95% confidence bounds)**

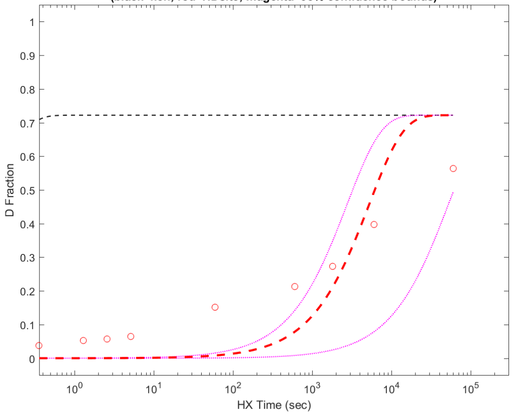


**Residue #203: HDsite kex=1.91e-04**
**(black=kch; red=HDsite; magenta=95% confidence bounds)**

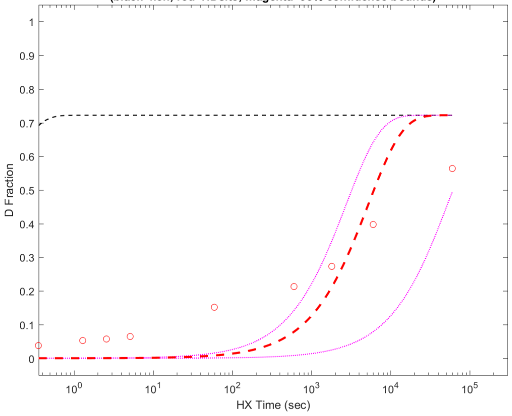


**Residue #204: HDsite kex=1.91e-04**
**(black=kch; red=HDsite; magenta=95% confidence bounds)**

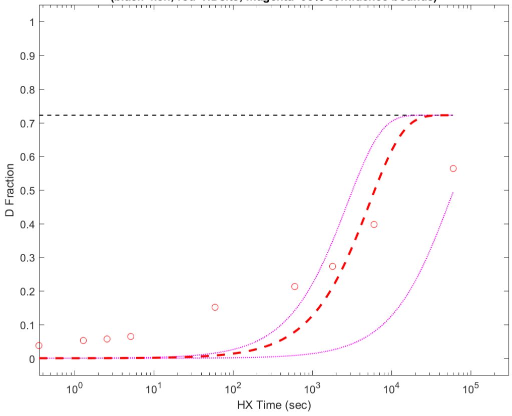


**Residue #205: HDsite kex=1.91e-04**
**(black=kch; red=HDsite; magenta=95% confidence bounds)**

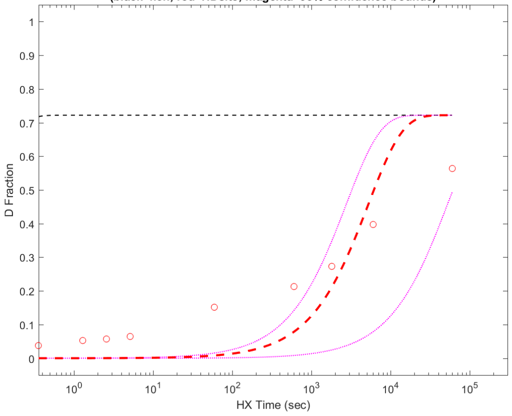


**Residue #206: HDsite kex=1.91e-04**
**(black=kch; red=HDsite; magenta=95% confidence bounds)**

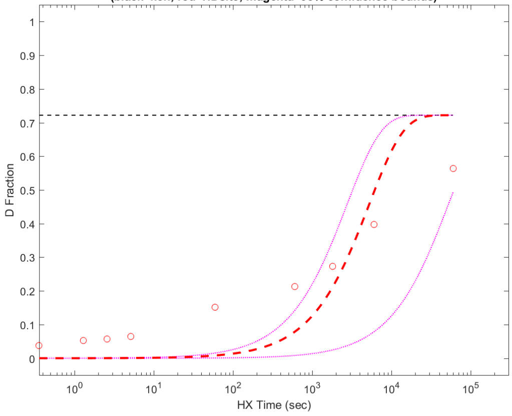


**Residue #207: HDsite kex=1.91e-04**
**(black=kch; red=HDsite; magenta=95% confidence bounds)**

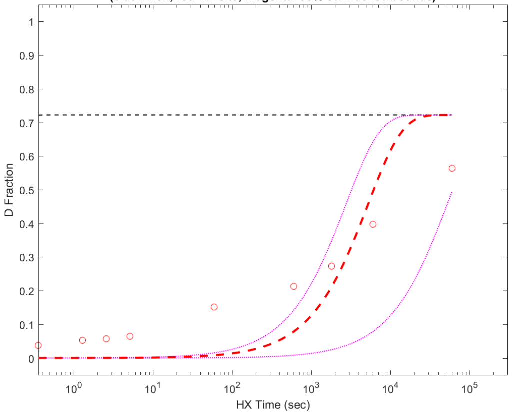

**Residue #208: HDsite kex=1.91e-04**
**(black=kch; red=HDsite; magenta=95% confidence bounds)**

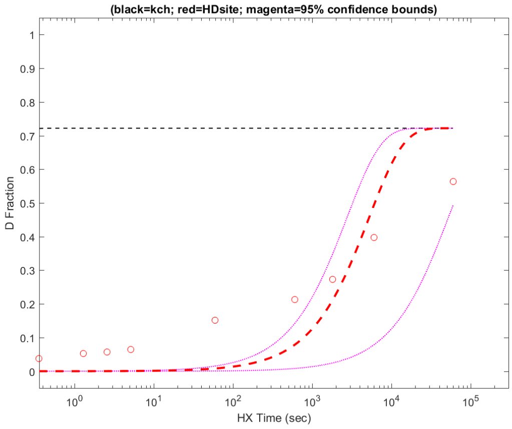


**Residue #209: HDsite kex=1.91e-04**
**(black=kch; red=HDsite; magenta=95% confidence bounds)**

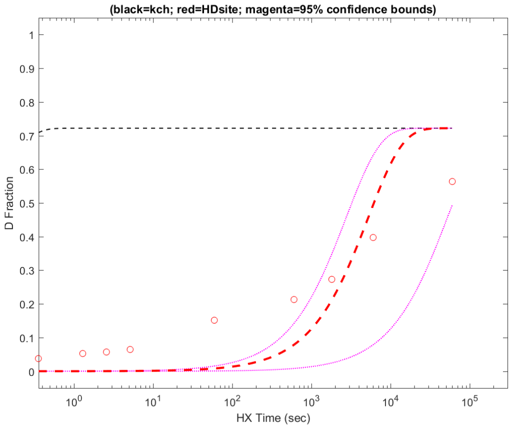


**Residue #210: HDsite kex=1.91e-04**
**(black=kch; red=HDsite; magenta=95% confidence bounds)**

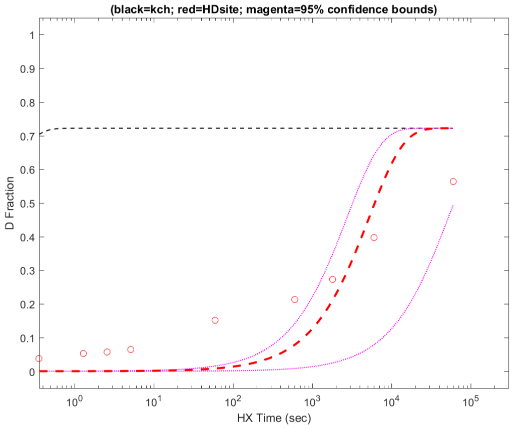


**Residue #211: HDsite kex=1.91e-04**
**(black=kch; red=HDsite; magenta=95% confidence bounds)**

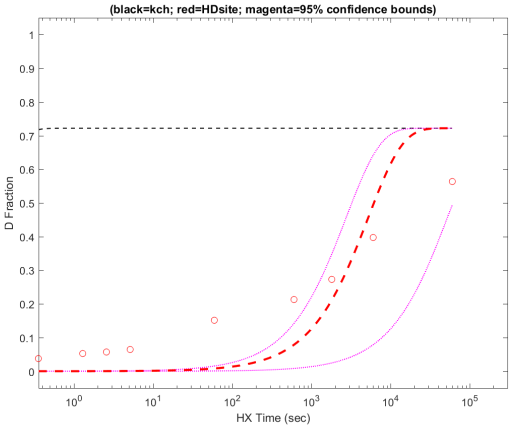


**Residue #212: HDsite kex=4.18e-04**
**(black=kch; red=HDsite; magenta=95% confidence bounds)**

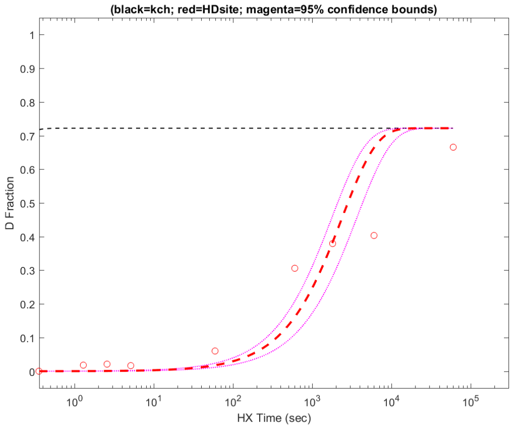


**Residue #213: HDsite kex=4.18e-04**
**(black=kch; red=HDsite; magenta=95% confidence bounds)**

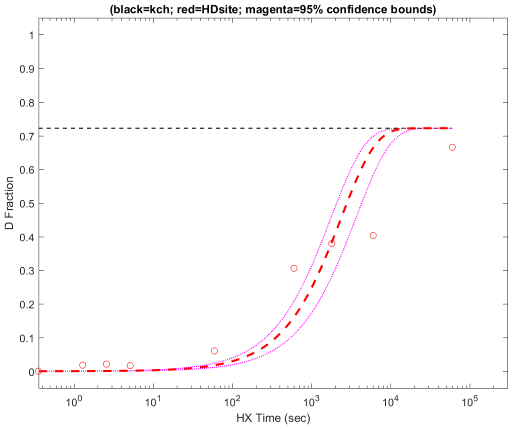


**Residue #214: HDsite kex=4.18e-04**
**(black=kch; red=HDsite; magenta=95% confidence bounds)**

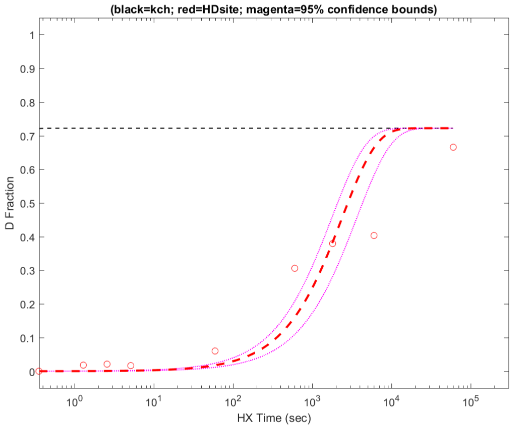


**Residue #215: HDsite kex=4.18e-04**
**(black=kch; red=HDsite; magenta=95% confidence bounds)**

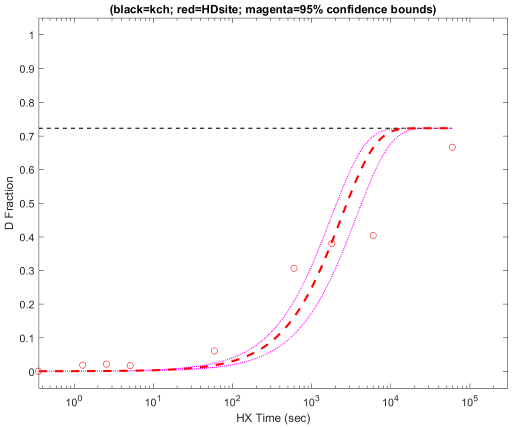

**Residue #216: HDsite kex=4.18e-04**
**(black=kch; red=HDsite; magenta=95% confidence bounds)**

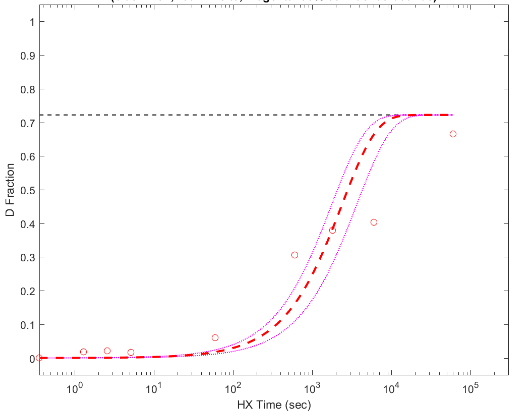


**Residue #217: HDsite kex=4.18e-04**
**(black=kch; red=HDsite; magenta=95% confidence bounds)**

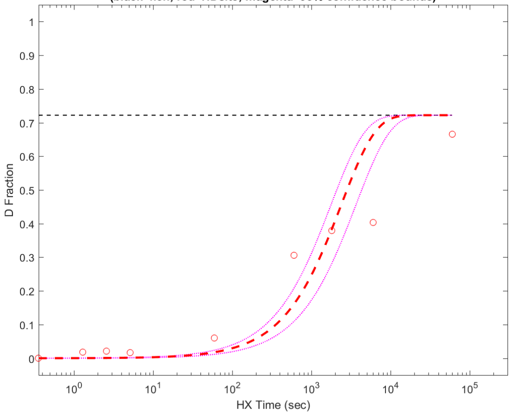


**Residue #218: HDsite kex=4.18e-04**
**(black=kch; red=HDsite; magenta=95% confidence bounds)**

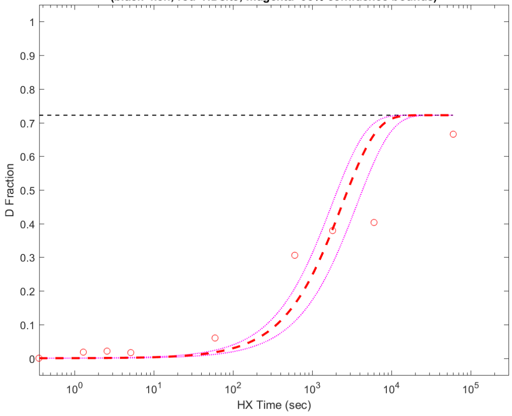


**Residue #219: HDsite kex=4.18e-04**
**(black=kch; red=HDsite; magenta=95% confidence bounds)**

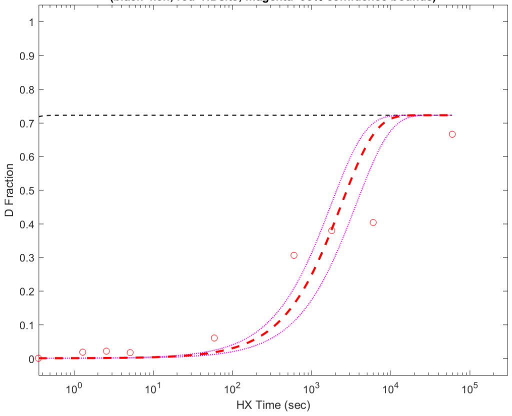


**Residue #220: HDsite kex=2.69e-05**
**(black=kch; red=HDsite; magenta=95% confidence bounds)**

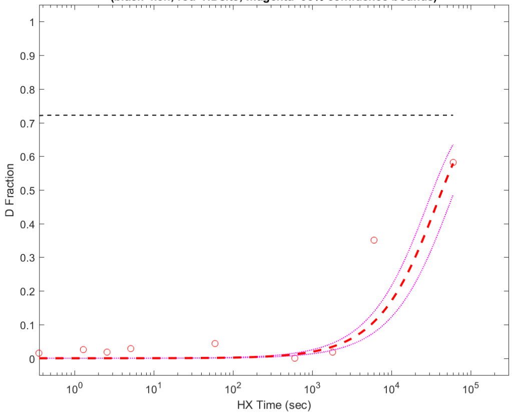


**Residue #221: HDsite kex=2.69e-05**
**(black=kch; red=HDsite; magenta=95% confidence bounds)**

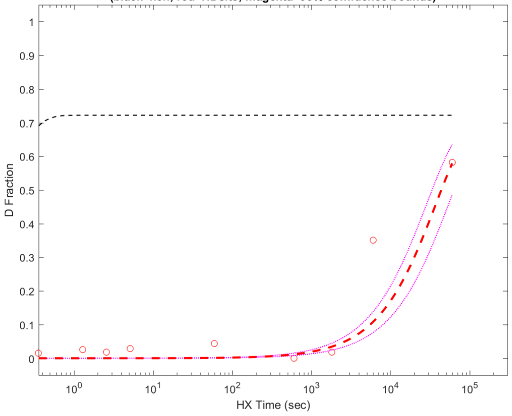


**Residue #222: HDsite kex=2.69e-05**
**(black=kch; red=HDsite; magenta=95% confidence bounds)**

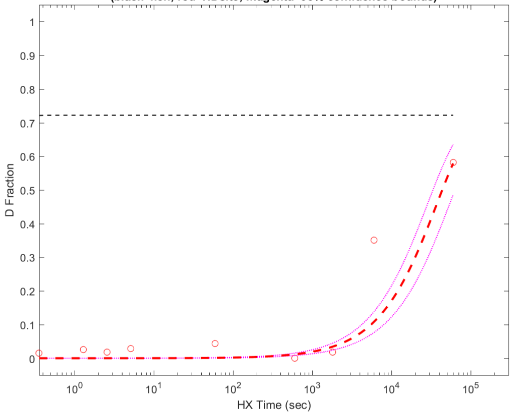


**Residue #223: HDsite kex=2.69e-05**
**(black=kch; red=HDsite; magenta=95% confidence bounds)**

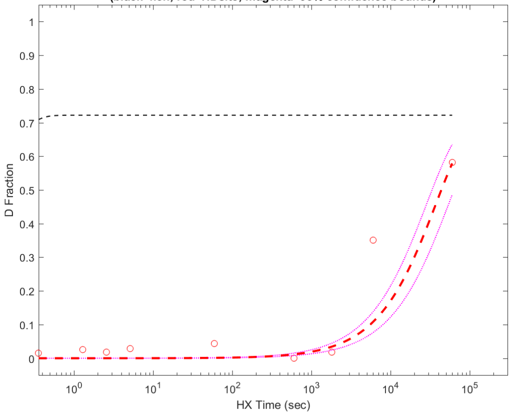

**Residue #224: HDsite kex=2.69e-05**
**(black=kch; red=HDsite; magenta=95% confidence bounds)**

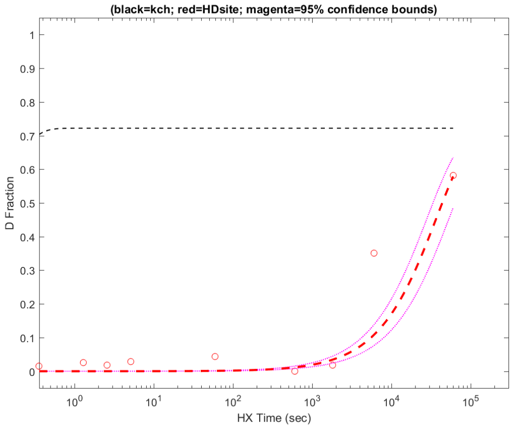


**Residue #225: HDsite kex=1.70e-04**
**(black=kch; red=HDsite; magenta=95% confidence bounds)**

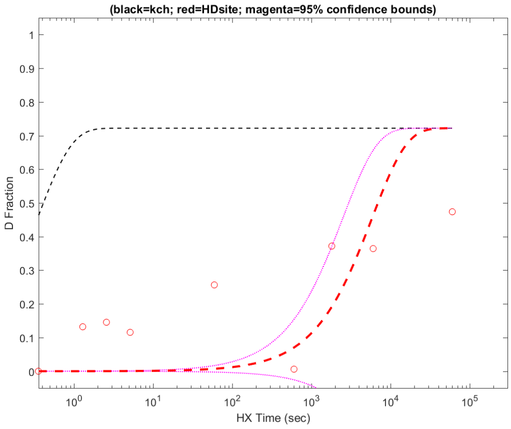


**Residue #227: HDsite kex=9.01e-04**
**(black=kch; red=HDsite; magenta=95% confidence bounds)**

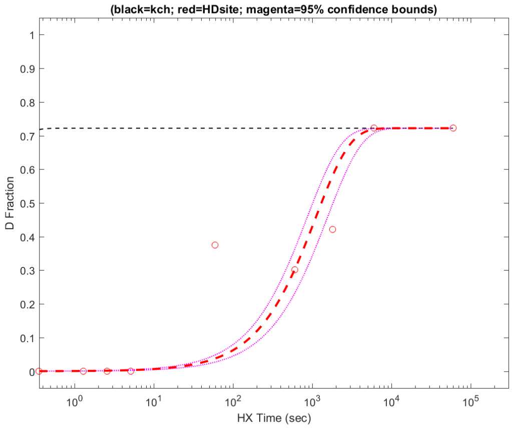


**Residue #228: HDsite kex=9.01e-04**
**(black=kch; red=HDsite; magenta=95% confidence bounds)**

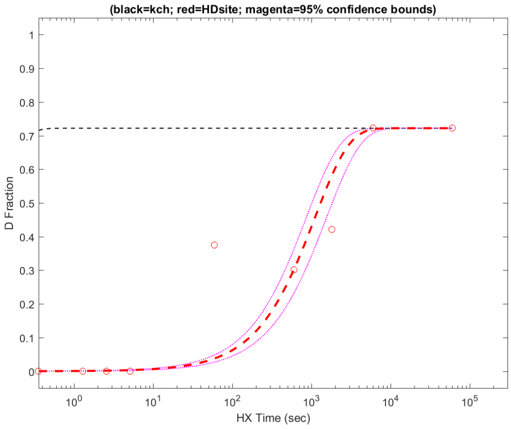


**Residue #229: HDsite kex=7.26e+00**
**(black=kch; red=HDsite; magenta=95% confidence bounds)**

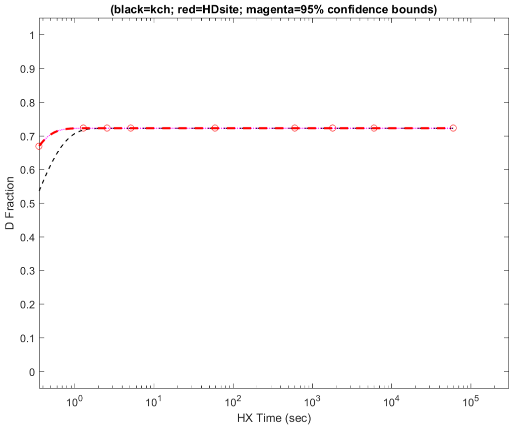


**Residue #230: HDsite kex=2.65e+00**
**(black=kch; red=HDsite; magenta=95% confidence bounds)**

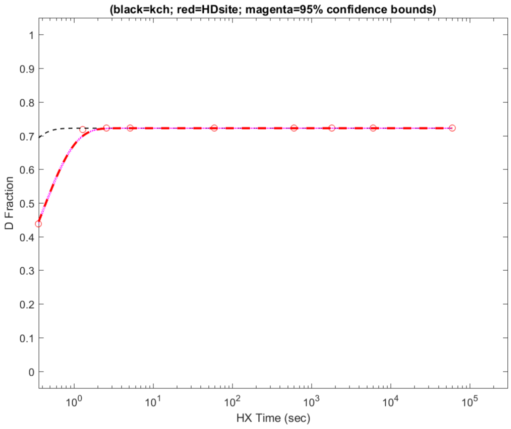


**Residue #231: HDsite kex=2.65e+00**
**(black=kch; red=HDsite; magenta=95% confidence bounds)**

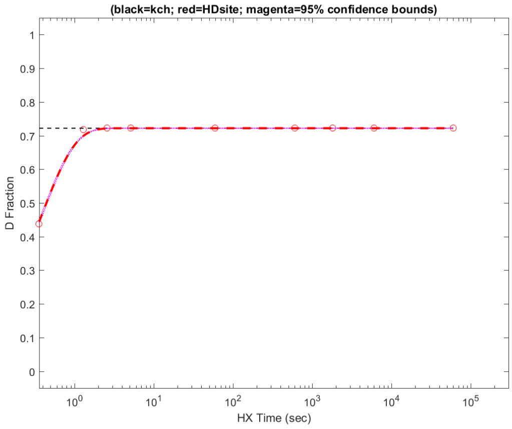


**Residue #232: HDsite kex=9.53e-03**
**(black=kch; red=HDsite; magenta=95% confidence bounds)**

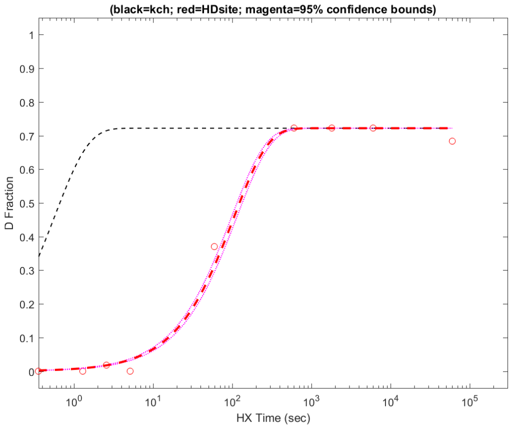

**Residue #233: HDsite kex=9.53e-03**
**(black=kch; red=HDsite; magenta=95% confidence bounds)**

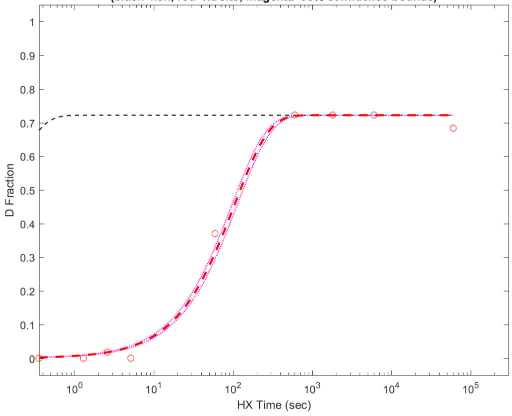


**Residue #234: HDsite kex=5.18e-02**
**(black=kch; red=HDsite; magenta=95% confidence bounds)**

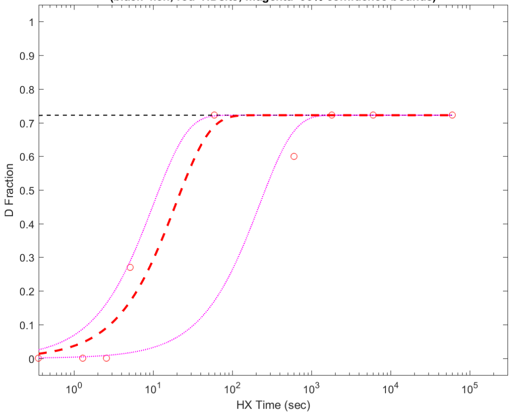


**Residue #235: HDsite kex=5.11e-05**
**(black=kch; red=HDsite; magenta=95% confidence bounds)**

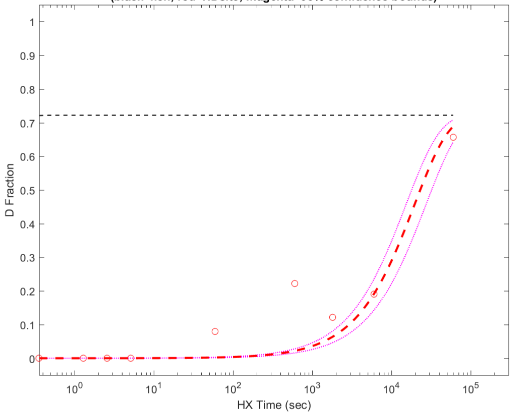


**Residue #236: HDsite kex=5.11e-05**
**(black=kch; red=HDsite; magenta=95% confidence bounds)**

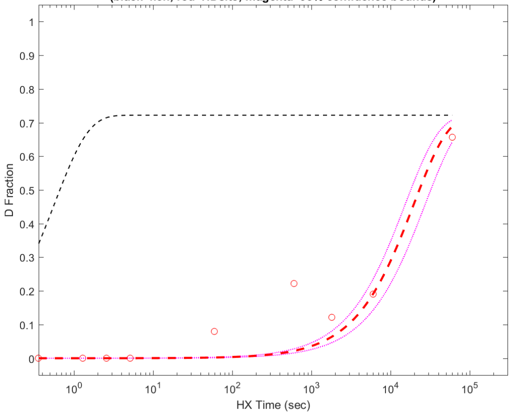


**Residue #237: HDsite kex=3.87e-04**
**(black=kch; red=HDsite; magenta=95% confidence bounds)**

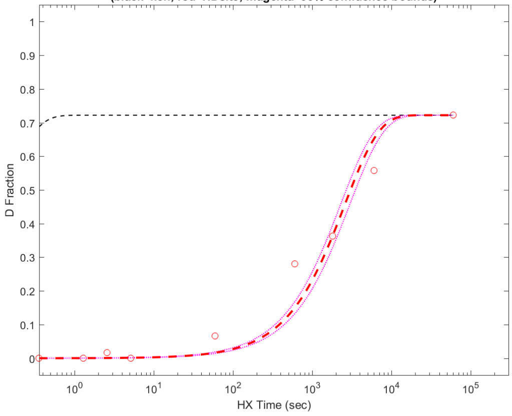


**Residue #238: HDsite kex=3.87e-04**
**(black=kch; red=HDsite; magenta=95% confidence bounds)**

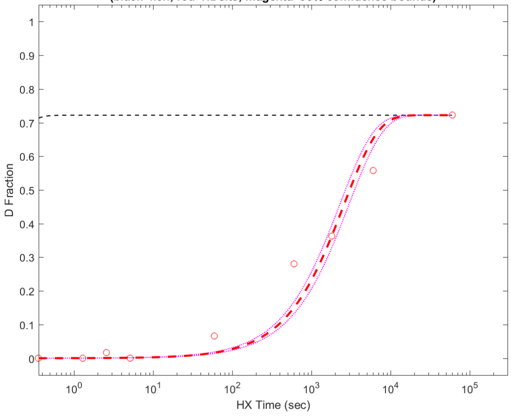


**Residue #239: HDsite kex=3.87e-04**
**(black=kch; red=HDsite; magenta=95% confidence bounds)**

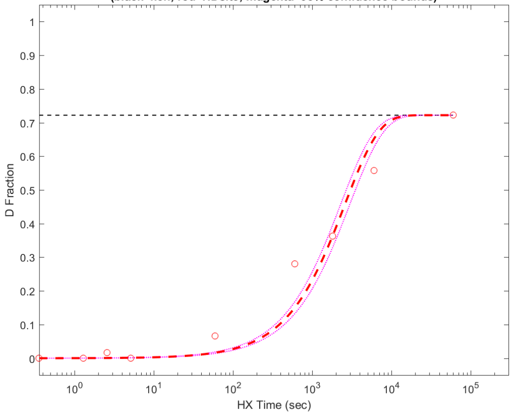


**Residue #240: HDsite kex=3.87e-04**
**(black=kch; red=HDsite; magenta=95% confidence bounds)**

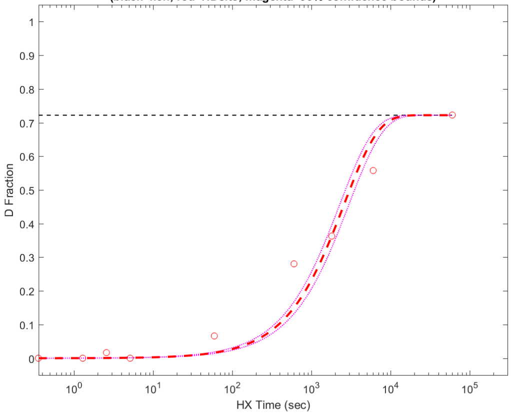

Residue #241: HDsite kex=5.31e-01
(black=kch; red=HDsite; magenta=95% confidence bounds)

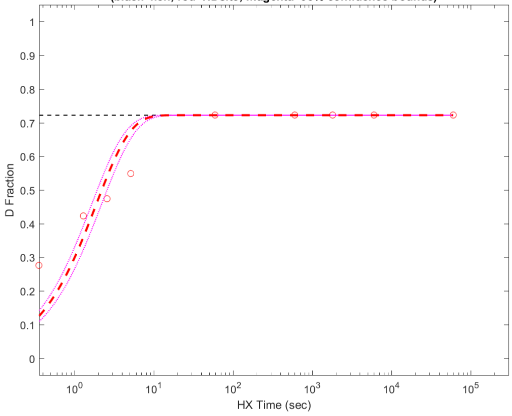


Residue #242: HDsite kex=5.31e-01
(black=kch; red=HDsite; magenta=95% confidence bounds)

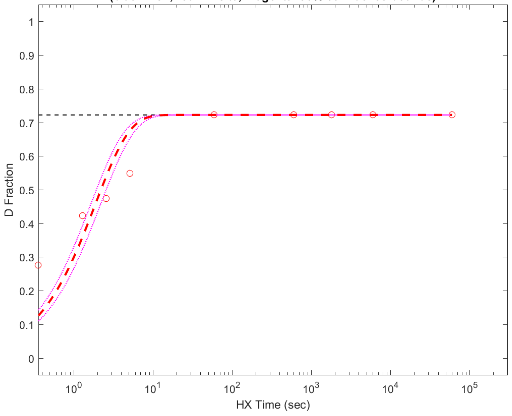


Residue #243: HDsite kex=5.31e-01
(black=kch; red=HDsite; magenta=95% confidence bounds)

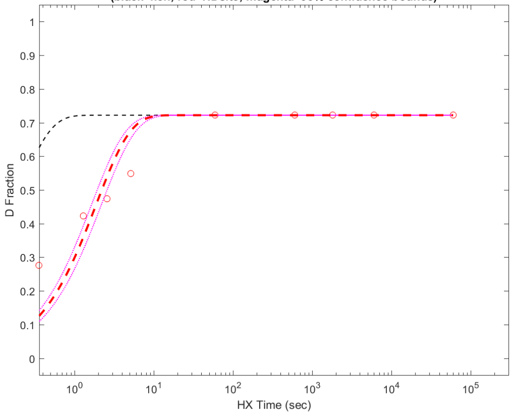


Residue #244: HDsite kex=5.31e-01
(black=kch; red=HDsite; magenta=95% confidence bounds)

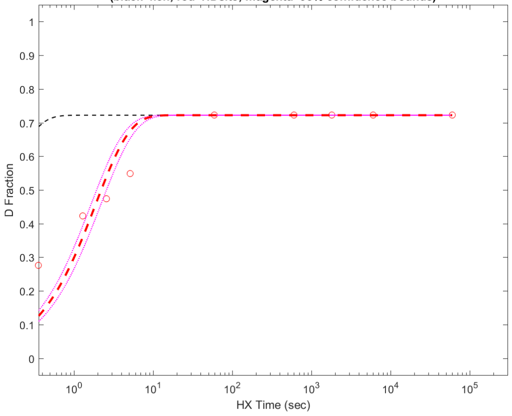


Residue #245: HDsite kex=1.49e-05
(black=kch; red=HDsite; magenta=95% confidence bounds)

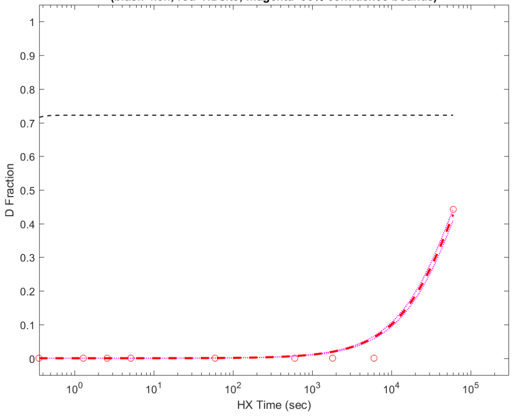


Residue #246: HDsite kex=1.49e-05
(black=kch; red=HDsite; magenta=95% confidence bounds)

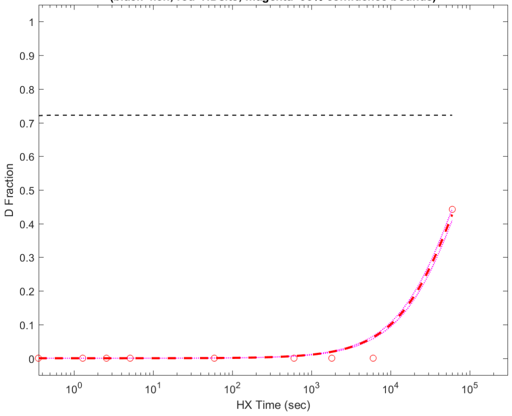


Residue #247: HDsite kex=1.49e-05
(black=kch; red=HDsite; magenta=95% confidence bounds)

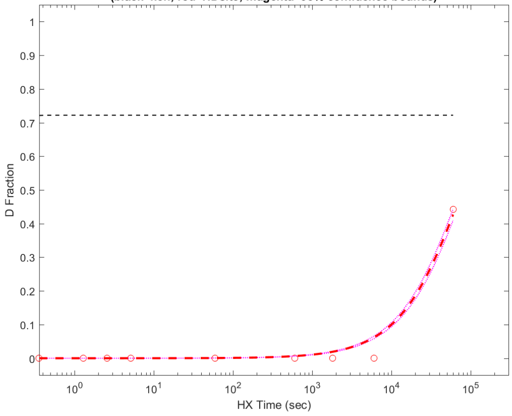


Residue #248: HDsite kex=1.59e-04
(black=kch; red=HDsite; magenta=95% confidence bounds)

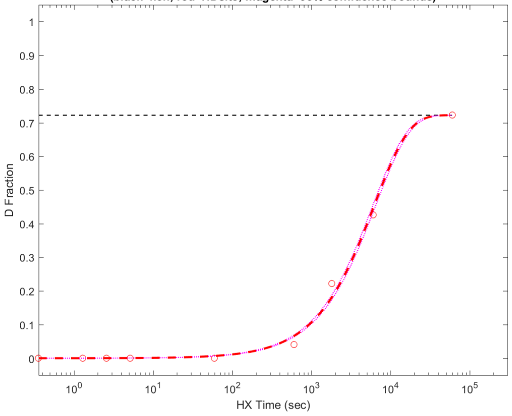

**Residue #249: HDsite kex=1.59e-04**
**(black=kch; red=HDsite; magenta=95% confidence bounds)**

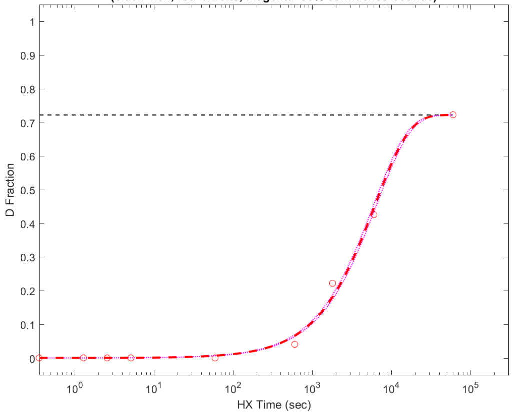


**Residue #250: HDsite kex=1.59e-04**
**(black=kch; red=HDsite; magenta=95% confidence bounds)**

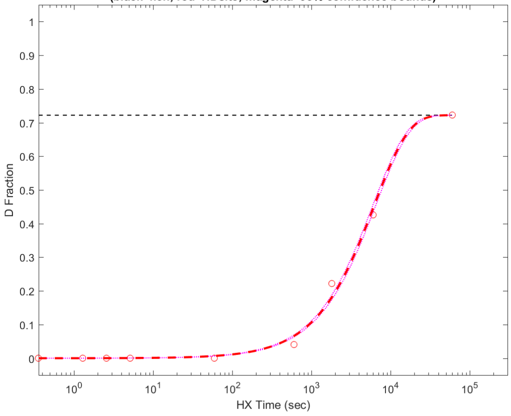


**Residue #251: HDsite kex=1.59e-04**
**(black=kch; red=HDsite; magenta=95% confidence bounds)**

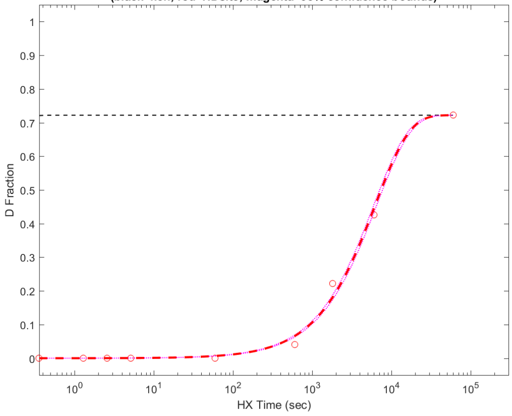


**Residue #252: HDsite kex=4.77e-06**
**(black=kch; red=HDsite; magenta=95% confidence bounds)**

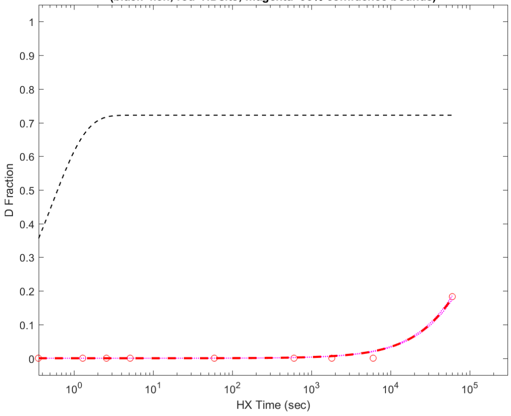


**Residue #253: HDsite kex=1.38e-04**
**(black=kch; red=HDsite; magenta=95% confidence bounds)**

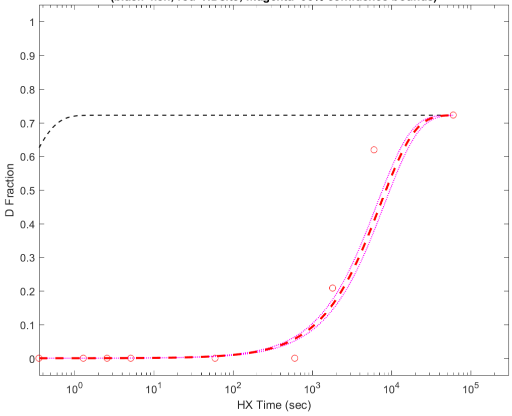


**Residue #254: HDsite kex=2.76e-03**
**(black=kch; red=HDsite; magenta=95% confidence bounds)**

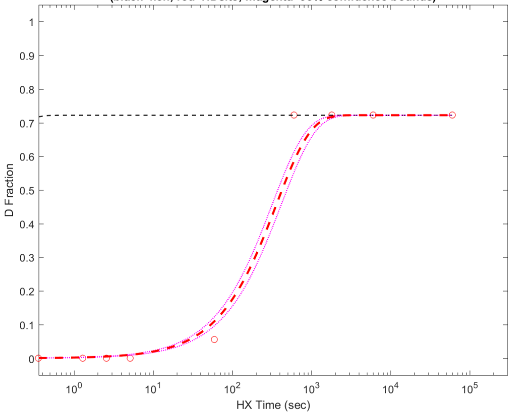


**Residue #255: HDsite kex=2.76e-03**
**(black=kch; red=HDsite; magenta=95% confidence bounds)**

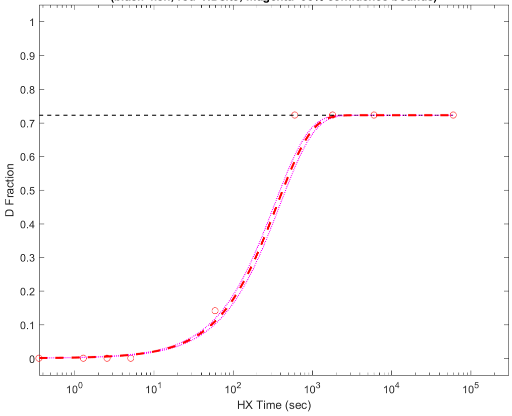


**Residue #256: HDsite kex=7.86e-03**
**(black=kch; red=HDsite; magenta=95% confidence bounds)**

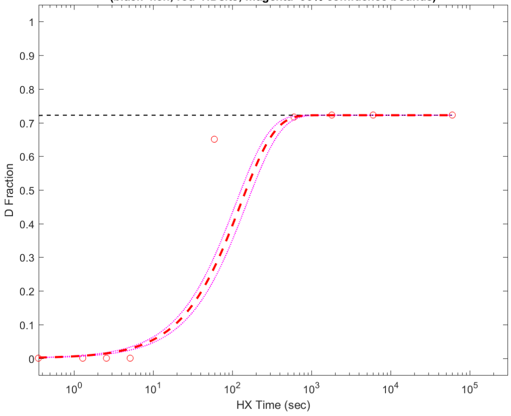

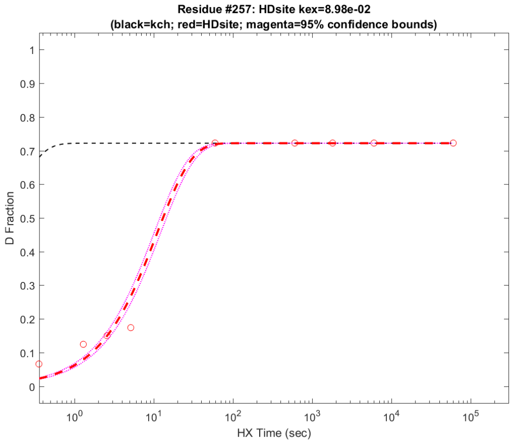
Residue #257: HDsite kex=8.98e-02
(black=kch; red=HDsite; magenta=95% confidence bounds)
D Fraction
HX Time (sec)

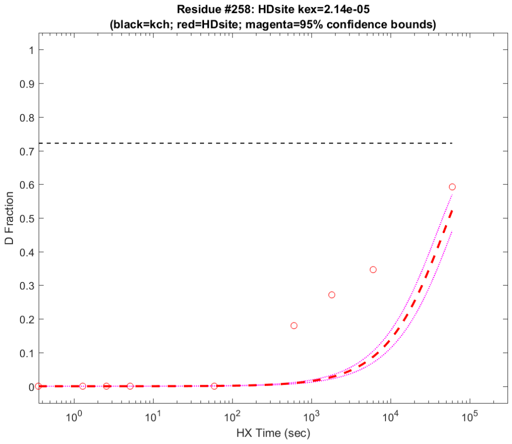
Residue #258: HDsite kex=2.14e-05
(black=kch; red=HDsite; magenta=95% confidence bounds)
D Fraction
HX Time (sec)

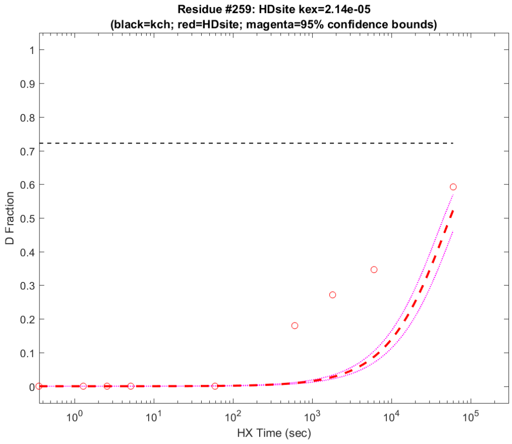
Residue #259: HDsite kex=2.14e-05
(black=kch; red=HDsite; magenta=95% confidence bounds)
D Fraction
HX Time (sec)

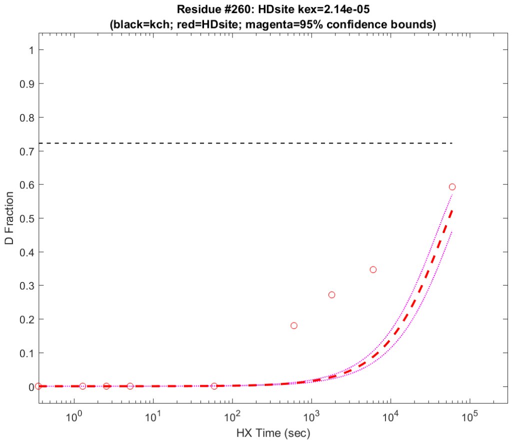
Residue #260: HDsite kex=2.14e-05
(black=kch; red=HDsite; magenta=95% confidence bounds)
D Fraction
HX Time (sec)

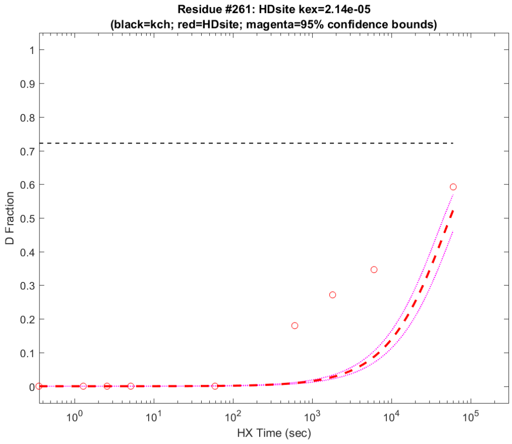
Residue #261: HDsite kex=2.14e-05
(black=kch; red=HDsite; magenta=95% confidence bounds)
D Fraction
HX Time (sec)

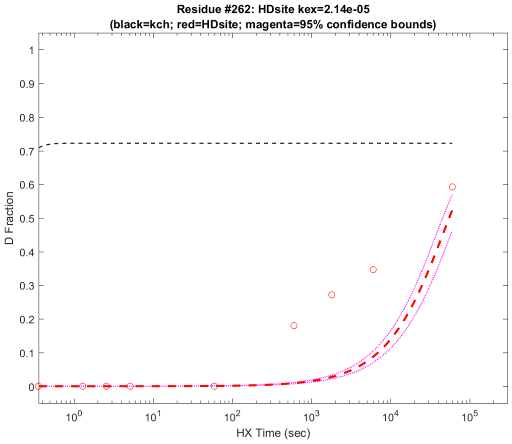
Residue #262: HDsite kex=2.14e-05
(black=kch; red=HDsite; magenta=95% confidence bounds)
D Fraction
HX Time (sec)

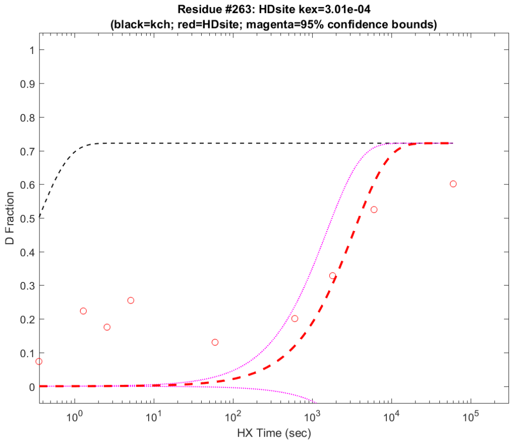
Residue #263: HDsite kex=3.01e-04
(black=kch; red=HDsite; magenta=95% confidence bounds)
D Fraction
HX Time (sec)

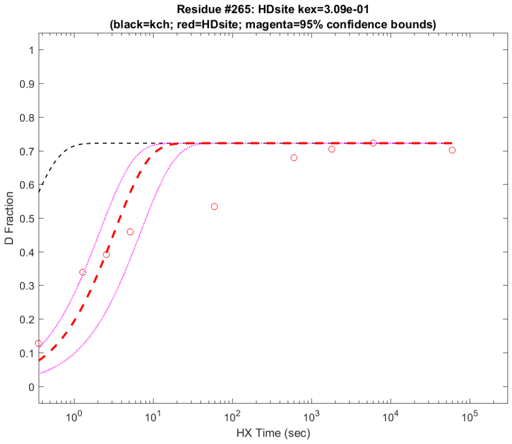
Residue #265: HDsite kex=3.09e-01
(black=kch; red=HDsite; magenta=95% confidence bounds)
D Fraction
HX Time (sec)

**Residue #266: HDsite kex=3.09e-01**
**(black=kch; red=HDsite; magenta=95% confidence bounds)**

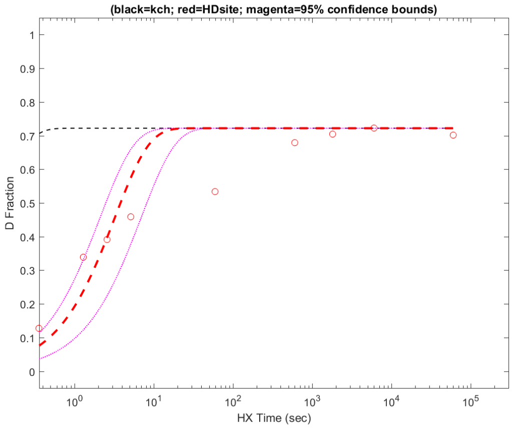


**Residue #267: HDsite kex=3.09e-01**
**(black=kch; red=HDsite; magenta=95% confidence bounds)**

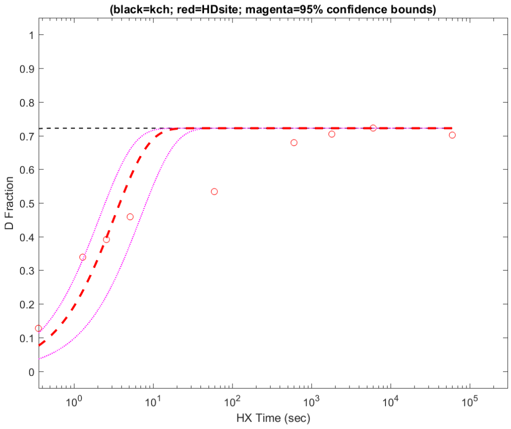


**Residue #268: HDsite kex=3.09e-01**
**(black=kch; red=HDsite; magenta=95% confidence bounds)**

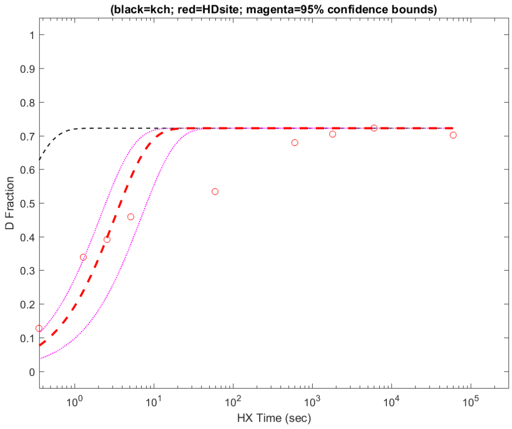


**Residue #269: HDsite kex=3.09e-01**
**(black=kch; red=HDsite; magenta=95% confidence bounds)**

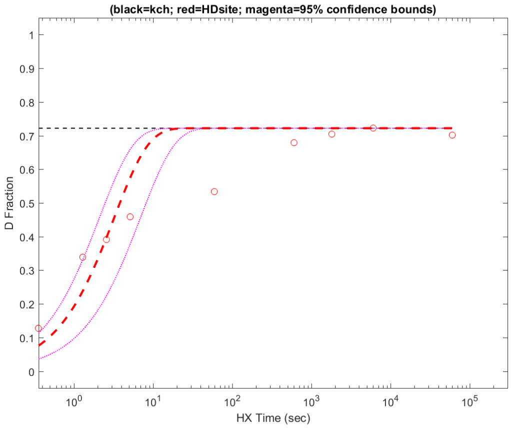


**Residue #270: HDsite kex=3.09e-01**
**(black=kch; red=HDsite; magenta=95% confidence bounds)**

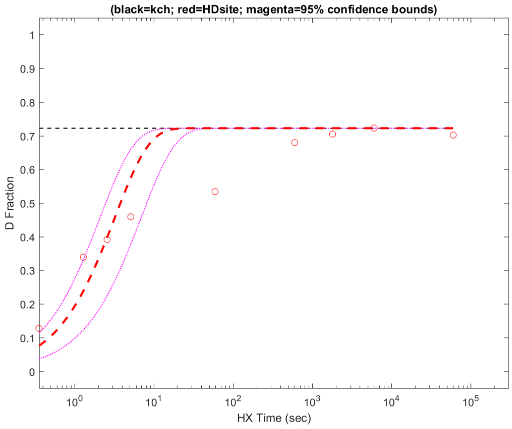


**Residue #271: HDsite kex=3.32e+00**
**(black=kch; red=HDsite; magenta=95% confidence bounds)**

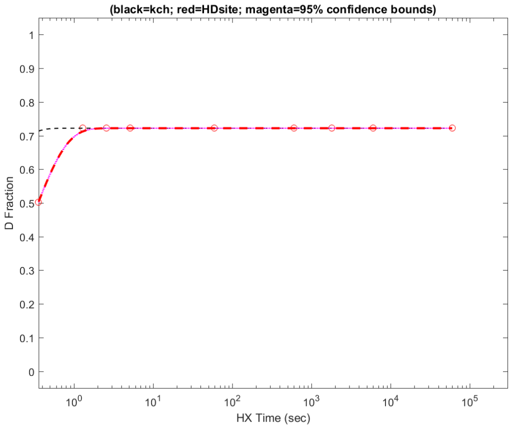


**Residue #272: HDsite kex=2.64e-03**
**(black=kch; red=HDsite; magenta=95% confidence bounds)**

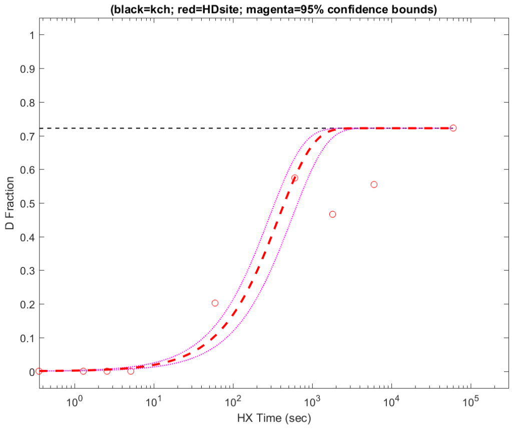


**Residue #273: HDsite kex=1.10e+00**
**(black=kch; red=HDsite; magenta=95% confidence bounds)**

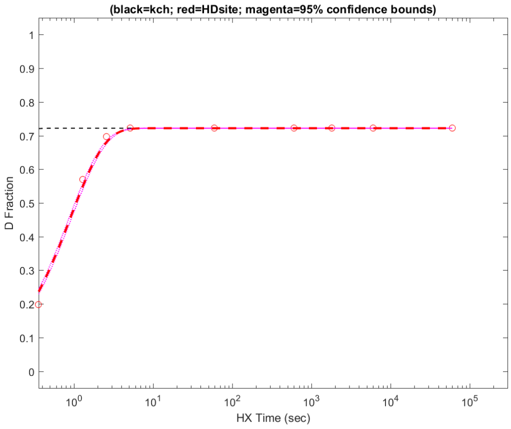

**Residue #274: HDsite kex=1.10e+00**
**(black=kch; red=HDsite; magenta=95% confidence bounds)**

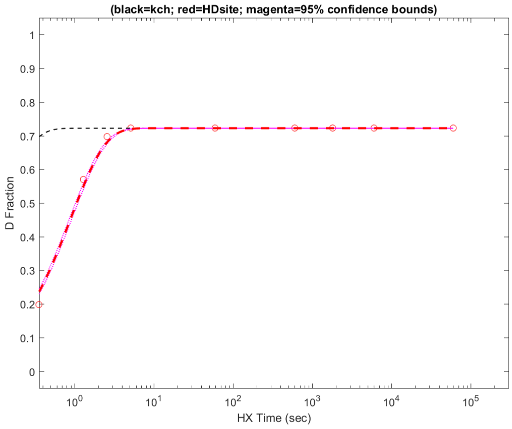


**Residue #275: HDsite kex=1.10e+00**
**(black=kch; red=HDsite; magenta=95% confidence bounds)**

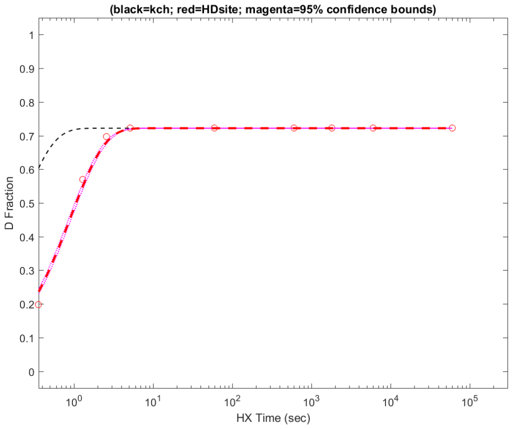


**Residue #276: HDsite kex=8.86e-03**
**(black=kch; red=HDsite; magenta=95% confidence bounds)**

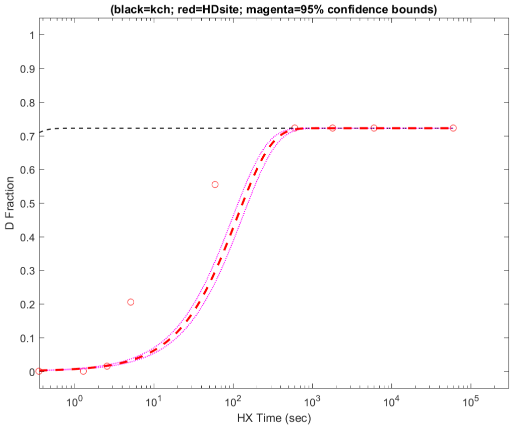


**Residue #277: HDsite kex=8.86e-03**
**(black=kch; red=HDsite; magenta=95% confidence bounds)**

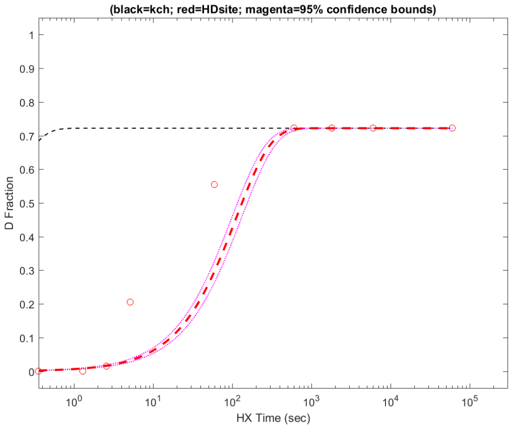


**Residue #278: HDsite kex=4.16e-04**
**(black=kch; red=HDsite; magenta=95% confidence bounds)**

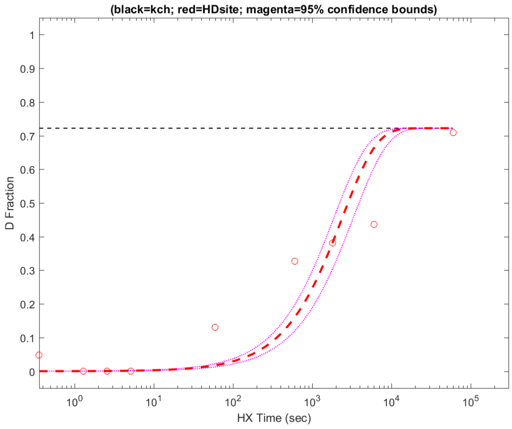


**Residue #279: HDsite kex=4.16e-04**
**(black=kch; red=HDsite; magenta=95% confidence bounds)**

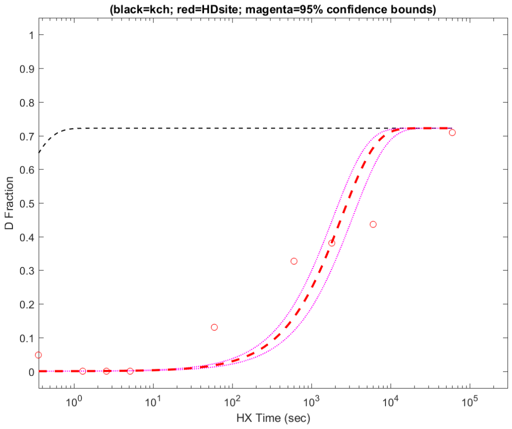


**Residue #280: HDsite kex=1.75e-05**
**(black=kch; red=HDsite; magenta=95% confidence bounds)**

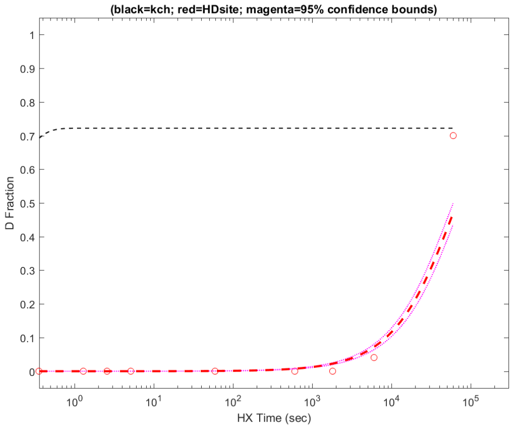


**Residue #281: HDsite kex=6.97e-06**
**(black=kch; red=HDsite; magenta=95% confidence bounds)**

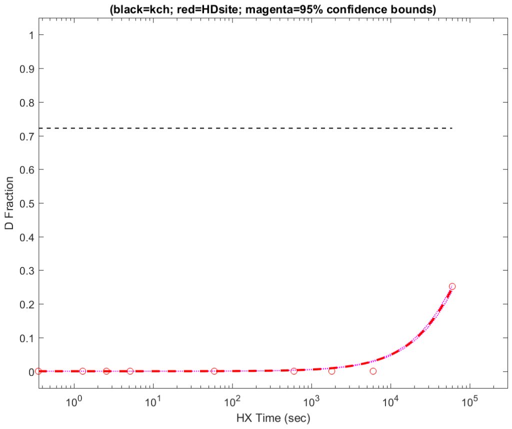

**Residue #282: HDsite kex=8.35e-04**
**(black=kch; red=HDsite; magenta=95% confidence bounds)**

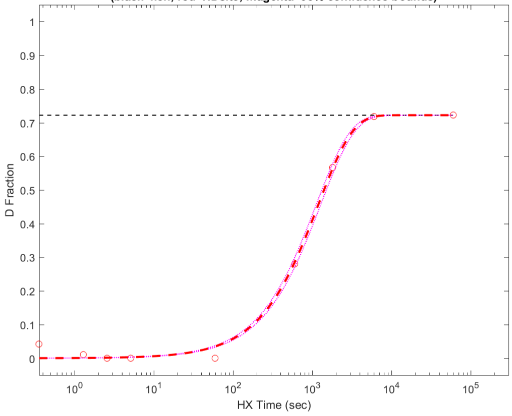


**Residue #283: HDsite kex=1.97e-05**
**(black=kch; red=HDsite; magenta=95% confidence bounds)**

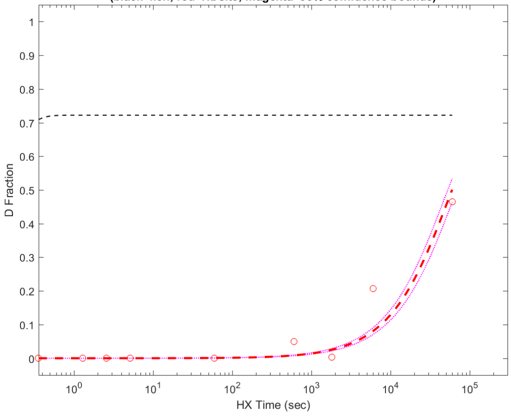


**Residue #284: HDsite kex=1.03e-04**
**(black=kch; red=HDsite; magenta=95% confidence bounds)**

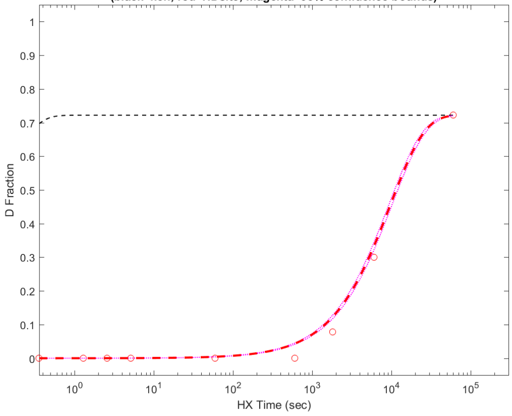


**Residue #285: HDsite kex=8.55e-07**
**(black=kch; red=HDsite; magenta=95% confidence bounds)**

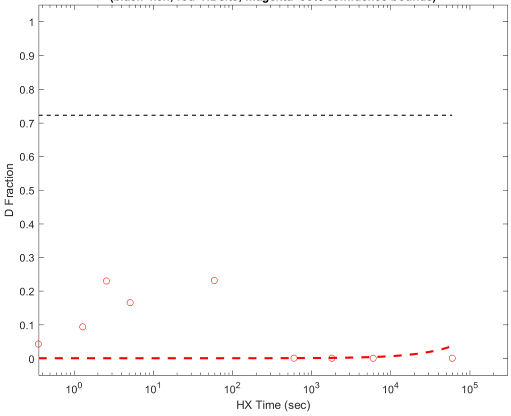


**Residue #286: HDsite kex=1.96e-05**
**(black=kch; red=HDsite; magenta=95% confidence bounds)**

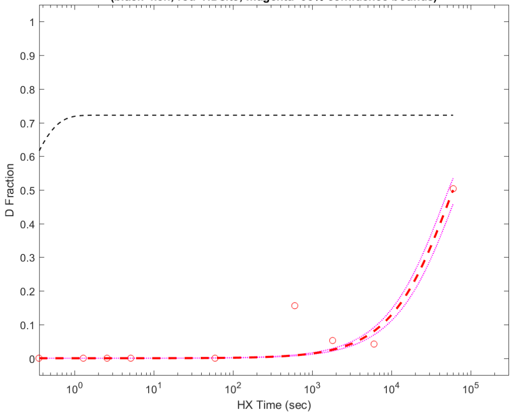


**Residue #287: HDsite kex=1.13e-04**
**(black=kch; red=HDsite; magenta=95% confidence bounds)**

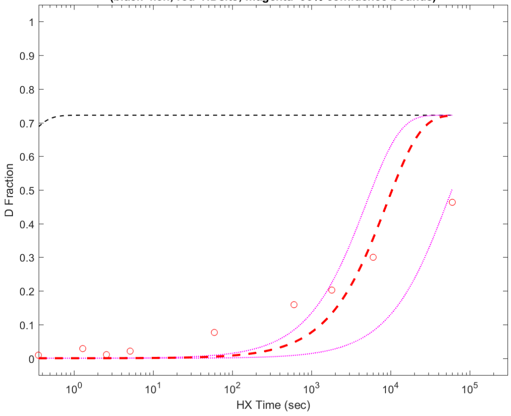


**Residue #288: HDsite kex=1.13e-04**
**(black=kch; red=HDsite; magenta=95% confidence bounds)**

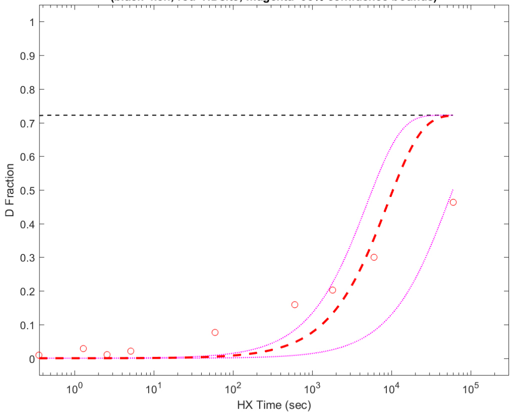


**Residue #289: HDsite kex=1.13e-04**
**(black=kch; red=HDsite; magenta=95% confidence bounds)**

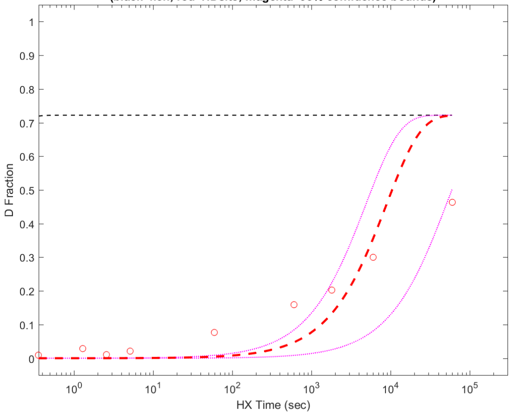

**Residue #290: HDsite kex=1.13e-04**
**(black=kch; red=HDsite; magenta=95% confidence bounds)**

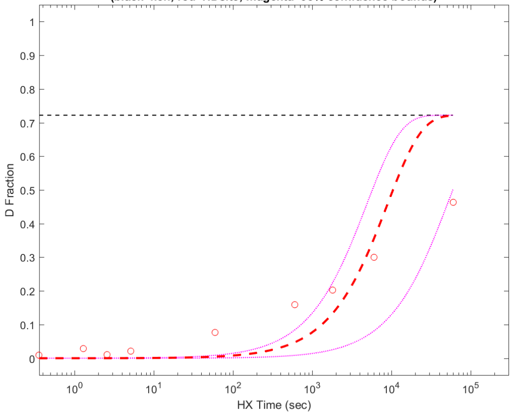


**Residue #291: HDsite kex=1.13e-04**
**(black=kch; red=HDsite; magenta=95% confidence bounds)**

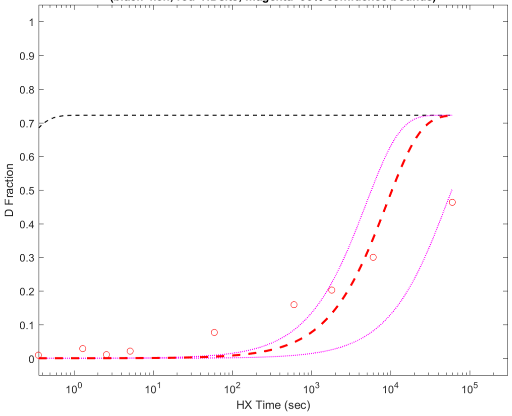


**Residue #292: HDsite kex=1.13e-04**
**(black=kch; red=HDsite; magenta=95% confidence bounds)**

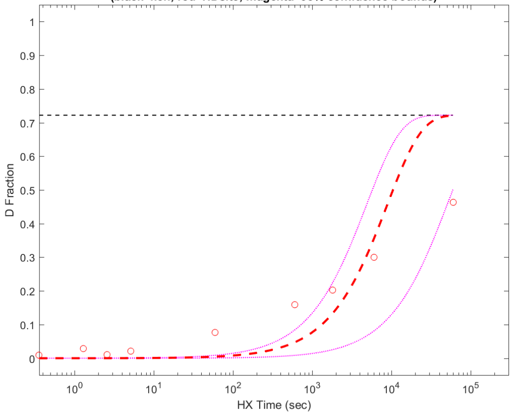


**Residue #293: HDsite kex=1.13e-04**
**(black=kch; red=HDsite; magenta=95% confidence bounds)**

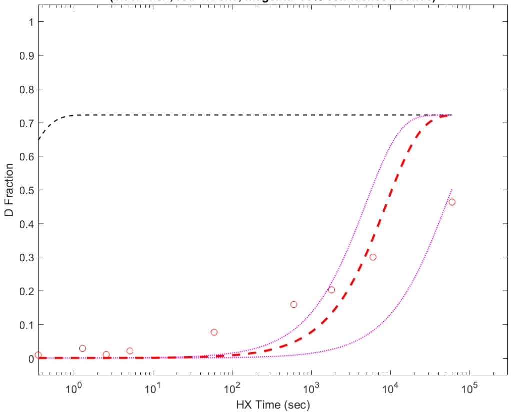


**Residue #294: HDsite kex=1.10e-03**
**(black=kch; red=HDsite; magenta=95% confidence bounds)**

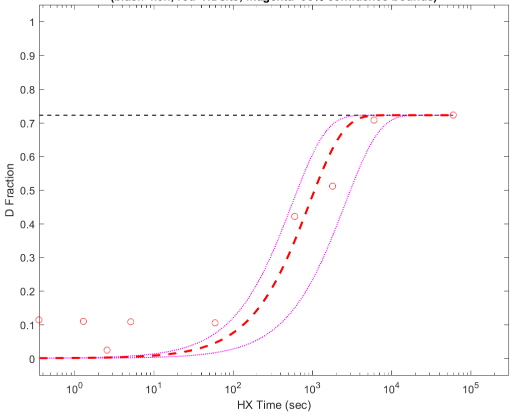


**Residue #295: HDsite kex=1.10e-03**
**(black=kch; red=HDsite; magenta=95% confidence bounds)**

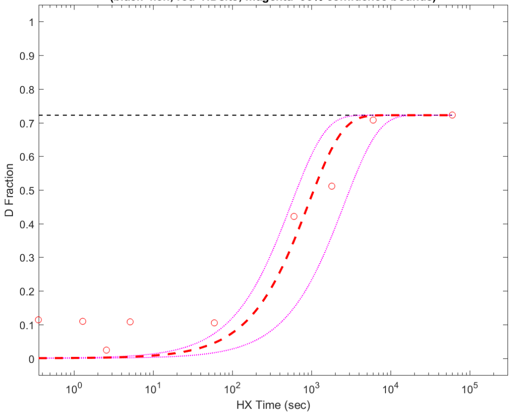


**Residue #296: HDsite kex=4.16e-04**
**(black=kch; red=HDsite; magenta=95% confidence bounds)**

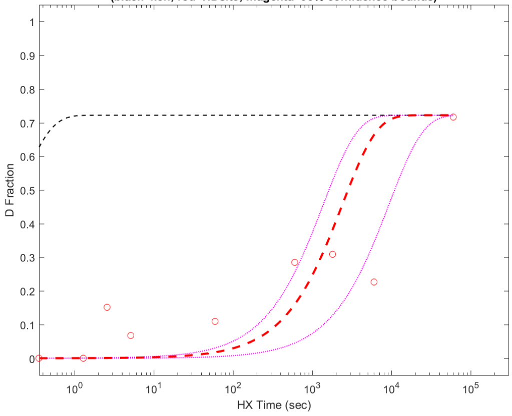


**Residue #297: HDsite kex=1.09e-02**
**(black=kch; red=HDsite; magenta=95% confidence bounds)**

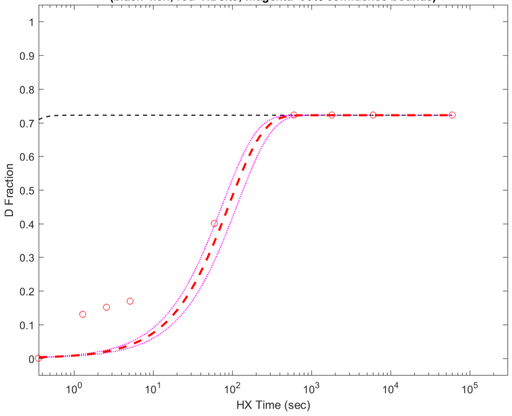

**Residue #298: HDsite kex=1.09e-02**
**(black=kch; red=HDsite; magenta=95% confidence bounds)**

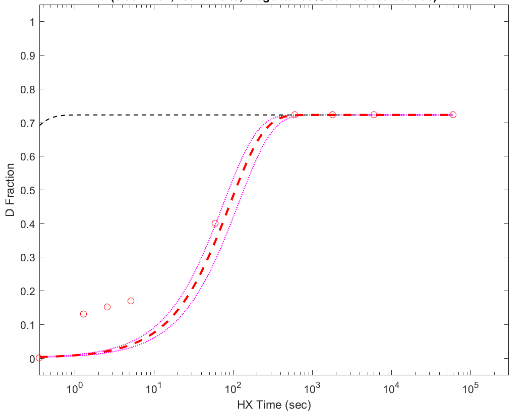


**Residue #299: HDsite kex=1.09e-02**
**(black=kch; red=HDsite; magenta=95% confidence bounds)**

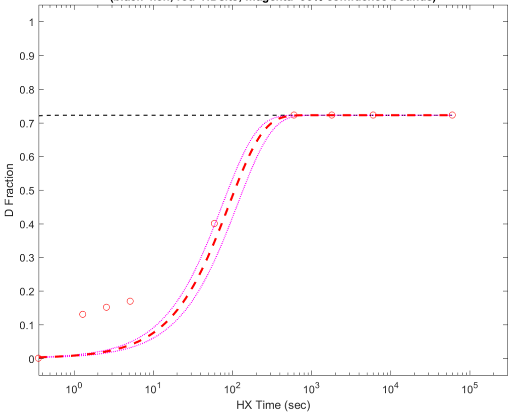


**Residue #300: HDsite kex=2.12e-04**
**(black=kch; red=HDsite; magenta=95% confidence bounds)**

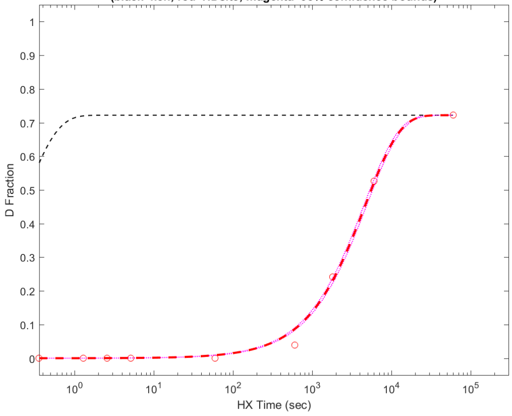


**Residue #301: HDsite kex=2.12e-04**
**(black=kch; red=HDsite; magenta=95% confidence bounds)**

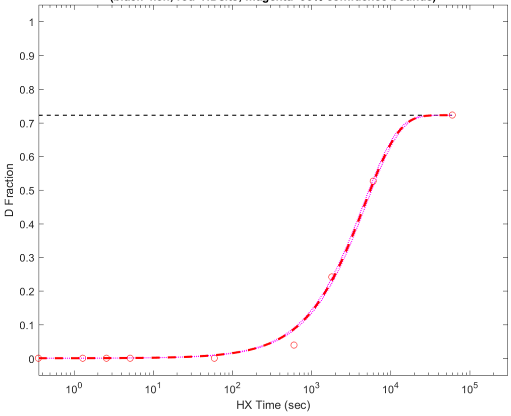


**Residue #302: HDsite kex=2.12e-04**
**(black=kch; red=HDsite; magenta=95% confidence bounds)**

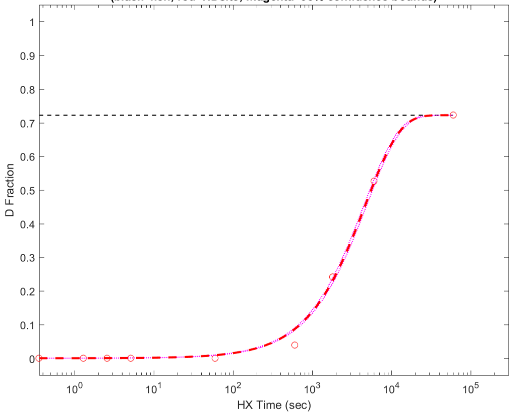


**Residue #303: HDsite kex=1.99e-03**
**(black=kch; red=HDsite; magenta=95% confidence bounds)**

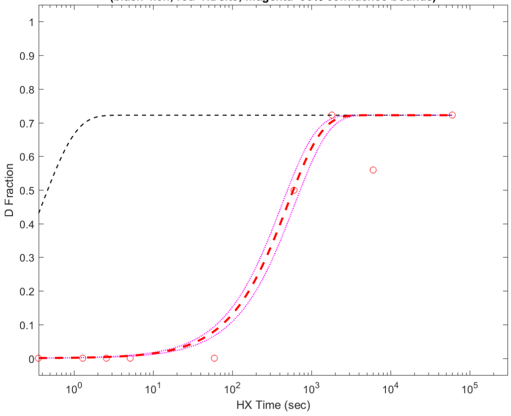


**Residue #304: HDsite kex=7.06e-03**
**(black=kch; red=HDsite; magenta=95% confidence bounds)**

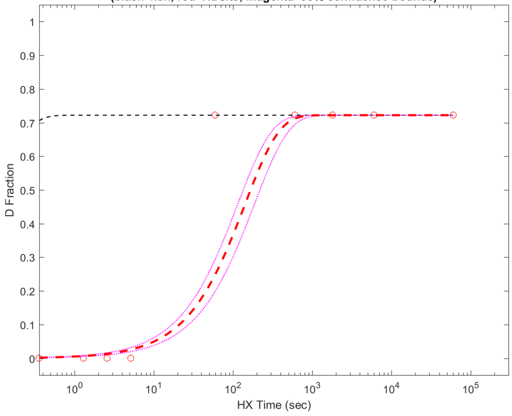


**Residue #305: HDsite kex=1.62e-01**
**(black=kch; red=HDsite; magenta=95% confidence bounds)**

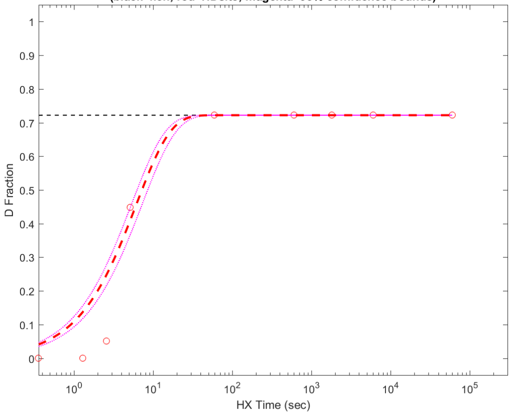

**Residue #306: HDsite kex=1.81e+00**
**(black=kch; red=HDsite; magenta=95% confidence bounds)**

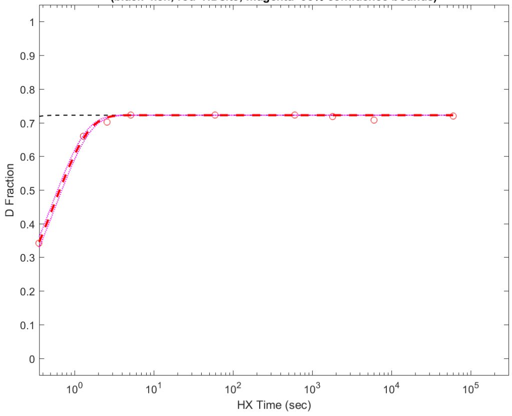


**Residue #307: HDsite kex=1.81e+00**
**(black=kch; red=HDsite; magenta=95% confidence bounds)**

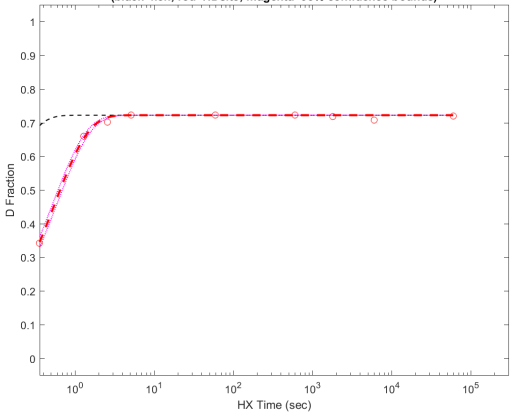


**Residue #308: HDsite kex=1.81e+00**
**(black=kch; red=HDsite; magenta=95% confidence bounds)**

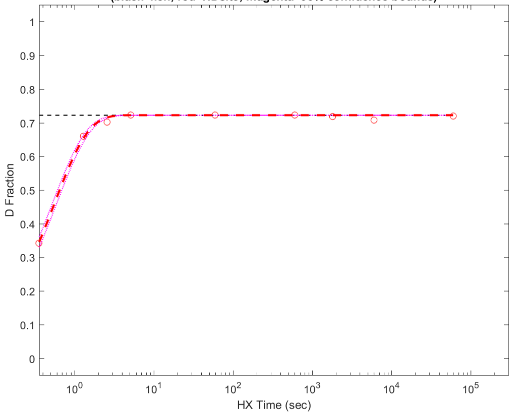


**Residue #309: HDsite kex=1.81e+00**
**(black=kch; red=HDsite; magenta=95% confidence bounds)**

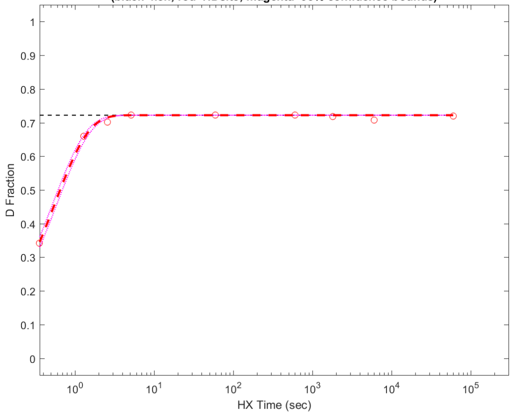


**Residue #310: HDsite kex=1.81e+00**
**(black=kch; red=HDsite; magenta=95% confidence bounds)**

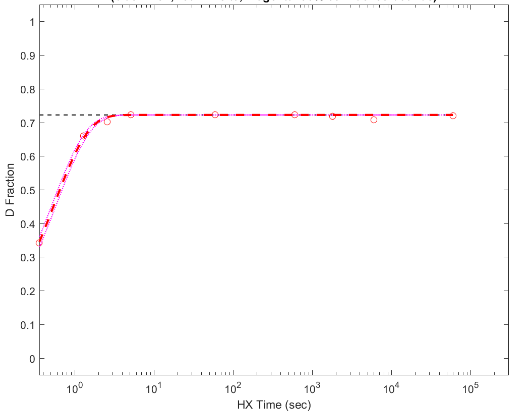


**Residue #311: HDsite kex=1.81e+00**
**(black=kch; red=HDsite; magenta=95% confidence bounds)**

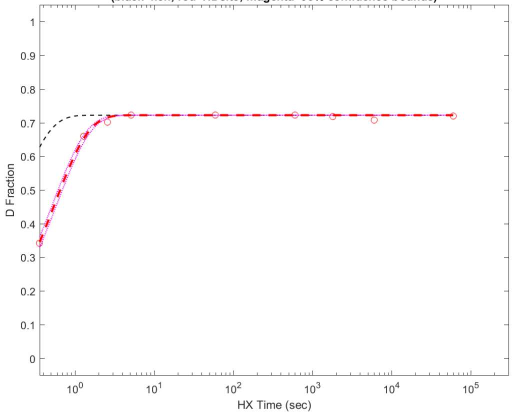


**Residue #312: HDsite kex=1.81e+00**
**(black=kch; red=HDsite; magenta=95% confidence bounds)**

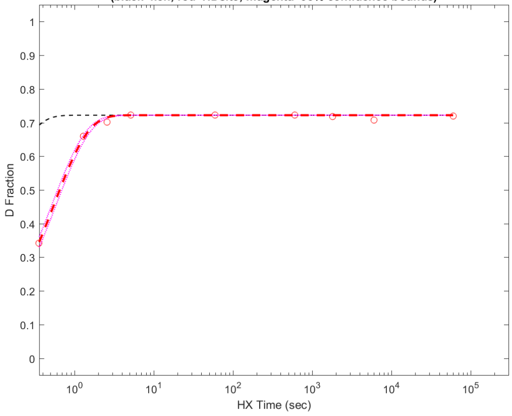


**Residue #313: HDsite kex=1.81e+00**
**(black=kch; red=HDsite; magenta=95% confidence bounds)**

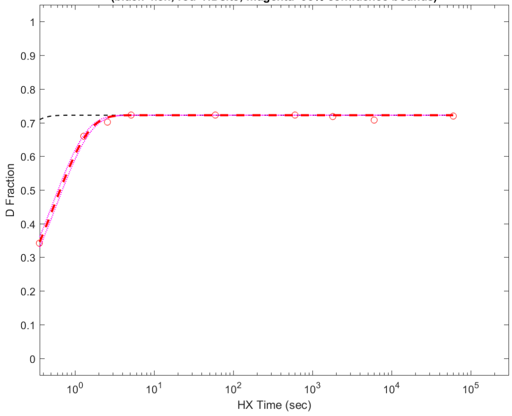

**Residue #314: HDsite kex=1.81e+00**
**(black=kch; red=HDsite; magenta=95% confidence bounds)**

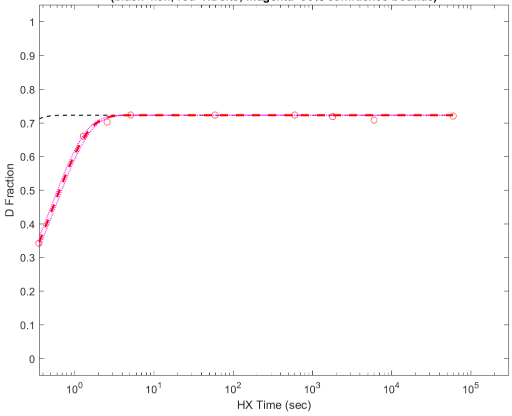


**Residue #315: HDsite kex=4.75e+00**
**(black=kch; red=HDsite; magenta=95% confidence bounds)**

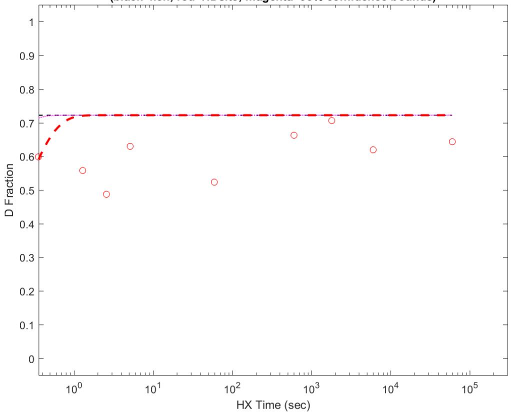


**Residue #316: HDsite kex=8.32e+00**
**(black=kch; red=HDsite; magenta=95% confidence bounds)**

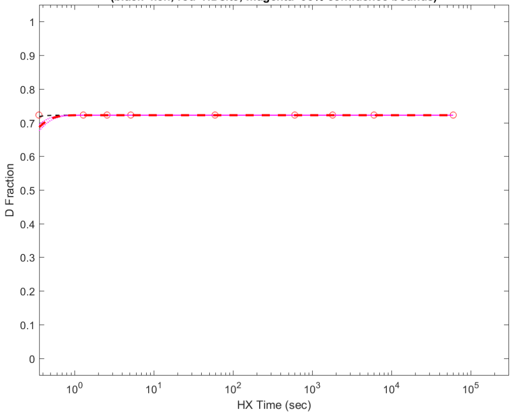


**Residue #317: HDsite kex=1.63e+00**
**(black=kch; red=HDsite; magenta=95% confidence bounds)**

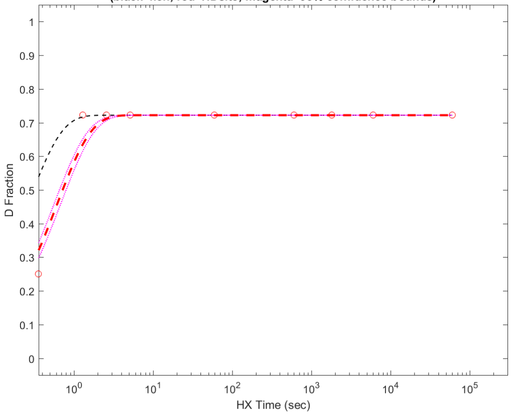


**Residue #318: HDsite kex=8.32e+00**
**(black=kch; red=HDsite; magenta=95% confidence bounds)**

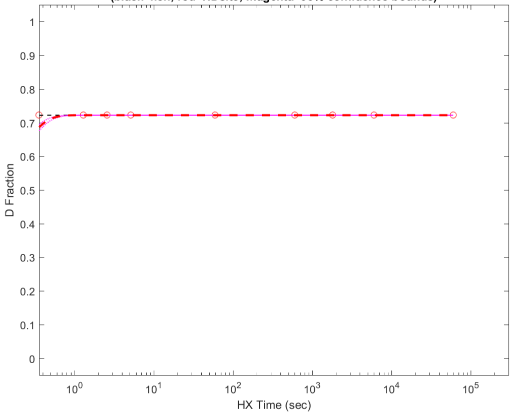


**Residue #319: HDsite kex=8.32e+00**
**(black=kch; red=HDsite; magenta=95% confidence bounds)**

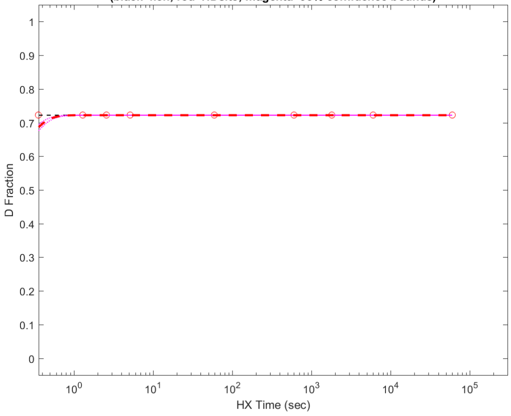


**Residue #320: HDsite kex=1.88e+00**
**(black=kch; red=HDsite; magenta=95% confidence bounds)**

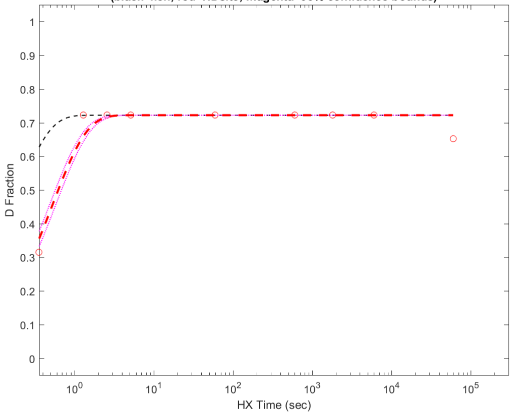


**Residue #321: HDsite kex=6.74e+00**
**(black=kch; red=HDsite; magenta=95% confidence bounds)**

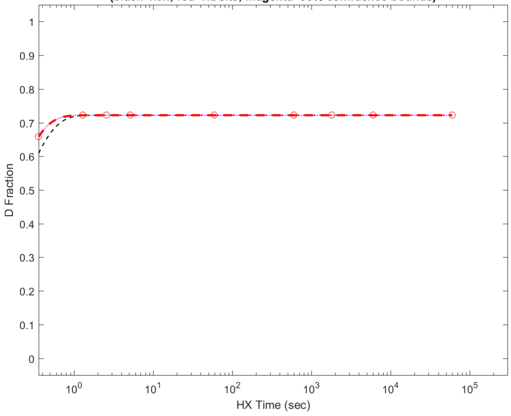

Residue #322: HDsite kex=6.74e+00
(black=kch; red=HDsite; magenta=95% confidence bounds)

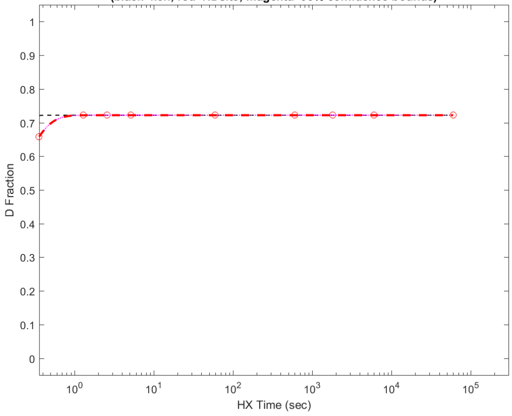


Residue #324: HDsite kex=3.36e+00
(black=kch; red=HDsite; magenta=95% confidence bounds)

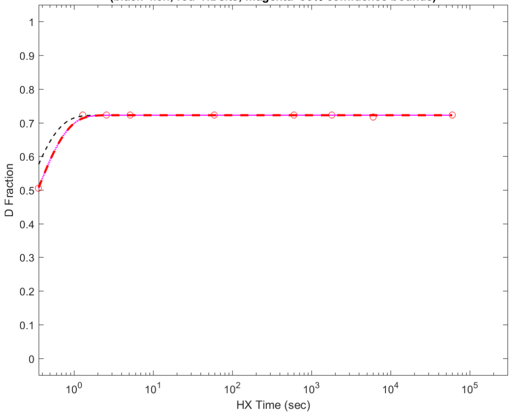


Residue #325: HDsite kex=3.36e+00
(black=kch; red=HDsite; magenta=95% confidence bounds)

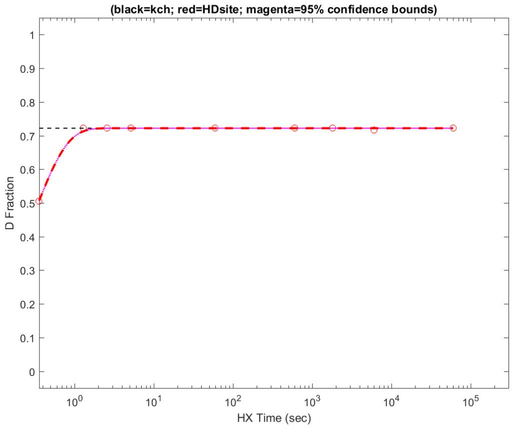


Residue #326: HDsite kex=3.36e+00
(black=kch; red=HDsite; magenta=95% confidence bounds)

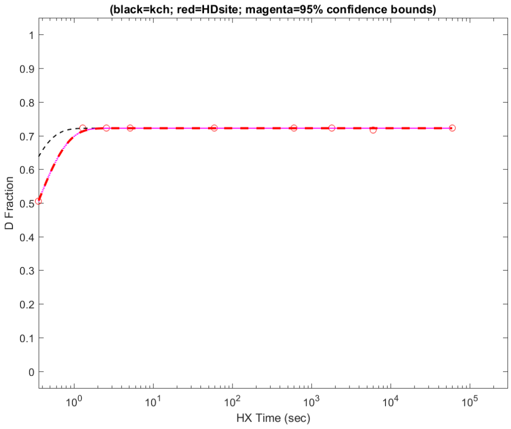


Residue #327: HDsite kex=3.36e+00
(black=kch; red=HDsite; magenta=95% confidence bounds)

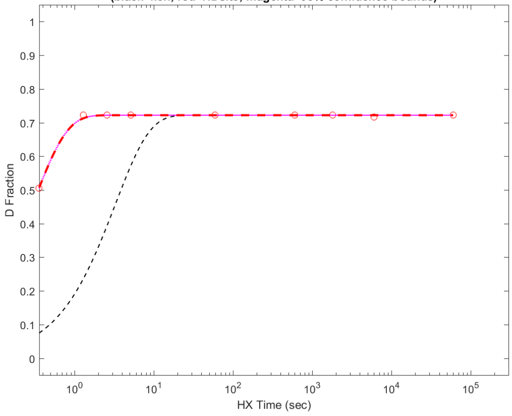
